# Biosynthetic Origin of the Methoxy Group in Quinine and Related Cinchona Alkaloids

**DOI:** 10.1101/2024.09.24.614729

**Authors:** Blaise Kimbadi Lombe, Tingan Zhou, Lorenzo Caputi, Kerstin Ploss, Sarah E. O’Connor

## Abstract

Quinine is a historically important natural product containing a methoxy group that is assumed to be incorporated at a late pathway stage. Here we show that the methoxy group in quinine and related Cinchona alkaloids is introduced onto the starting substrate tryptamine. Feeding studies with Cinchona plantlets definitively show that 5-methoxytryptamine is utilized as a quinine biosynthetic intermediate *in planta*. We discover the biosynthetic genes that encode the responsible oxidase and methyltransferase, and we use these genes to reconstitute the early steps of the Cinchona alkaloid biosynthetic pathway in *Nicotiana benthamiana* to produce a mixture of methoxylated and desmethoxylated Cinchona alkaloid intermediates. Importantly, we show that the co-occurrence of both tryptamine and 5-methoxytryptamine substrates, along with the substrate promiscuity of downstream pathway enzymes, enable parallel formation of both methoxylated and desmethoxylated alkaloids in *Cinchona pubescens*.

---

Quinine (**1**), used for centuries as an antimalarial drug and as a bittering ingredient, is a member of the Cinchona alkaloids, a structurally diverse group of alkaloids produced primarily by *Cinchona* plants.^[1]^ In addition to quinine (**1**), the quinine stereoisomer, quinidine (**2**), is commonly prescribed as an antiarrhythmic medication.^[2]^ Desmethoxylated analogs (*e*.*g*. cinchonidine **3** and cinchonine **4**) also have established pharmacological properties^[1d, 3]^ and, moreover, are used in synthetic chemistry as chiral catalysts and additives.^[4]^ Congeners with a single bond at C-10 – C-11 (dihydro alkaloids; for structures, see **5-8** in Scheme S1 or Figure S1) also occur in several *Cinchona* species and have similar properties as **1-4**.^[1d,3a]^ While total syntheses of many Cinchona alkaloids have been reported,^[5]^ the biosynthesis of these compounds is still largely unelucidated.

Early investigations established that Cinchona alkaloids belong to the large family of monoterpenoid indole alkaloids.^[6]^ These alkaloids are derived from strictosidine (**12**), which is formed from the Pictet-Spengler condensation of a tryptophan-derived moiety, tryptamine (**10**), and the iridoid-type monoterpene secologanin (**11**).^[7]^ It has long been hypothesized that strictosidine is converted to cinchonidinone (**15a**) and cinchoninone (**15b**), compounds that are then methoxylated to form quinine (**1**) and quinidine (**2**), respectively (Scheme 1).^[8]^ Here we show through feeding experiments that methoxylation instead occurs on the early precursor tryptamine (**10**). We show that both tryptamine and 5-methoxytryptamine (**21**) are carried through the downstream pathway, leading to mixtures of methoxy (*e*.*g*. **1, 2**) and desmethoxy (*e*.*g*. **3, 4**) Cinchona alkaloids *in planta*. Additionally, we report the discovery of the biosynthetic genes that are responsible for this methoxylation, allowing reconstruction of the early biosynthetic steps of desmethoxy and methoxy Cinchona alkaloid intermediates.

Methoxylation of cinchonidinone (**15a**) and cinchoninone (**15b**) was first proposed^[8a]^ based on the incorporation of the radio-labeled compounds corynantheal (**13**) and cinchonidinone (**15a**) into quinine (**1**), though the incorporation was marginal (ca. 0.002%).^[8b, 8c]^ Additionally, methoxylated ketone **17a** has been detected in *C. ledgeriana*,^[8c]^ and crude extracts of a cell culture derived from this species showed the presence of NADPH-dependent reduction activity that specifically reduced **17a/b** to **1** and **2**.^[8f]^ Moreover, a recently discovered *O*-methyltransferase from *C. pubescens* can methylate 6′-hydroxycinchoninone (**16a/16b**).^[8d]^ Together, these data are consistent with – though do not prove – a pathway order of hydroxylation of **15a/b**, *O*-methylation, followed by keto-reduction to form **1** and **2** (Scheme 1). We set out to identify the missing oxidase that would hydroxylate cinchonidinone/cinchoninone (**15a/b**). In *C. pubescens*, **1** and **2**, along with the C10 – C11 dihydro analogs (**5** and **6**), accumulate only in roots and stem,^[8d]^ leading us to hypothesize that the **15a/b**-hydroxylase would be specifically expressed in these two tissues. RNA-seq data from *C. pubescens* were therefore mined and a total of 150 putative oxidases (P450s, BBE, and PPOs) that were enriched in these tissues were selected, cloned into *Agrobacterium tumefaciens* and transiently expressed in *N. benthamiana* leaves. Infiltration of **15a/b** into the treated *N. benthamiana* leaves, followed by LC-MS analysis of the crude transformed leaf extracts, indicated that none of the enzyme candidates could hydroxylate or otherwise metabolize **15a/b**.

This failure to identify **15a/b**-hydroxylase, along with previously reported unsuccessful attempts,^[8d]^ prompted us to consider that methoxylation may occur at an earlier stage of the biosynthetic pathway. Strictosidine (**12**), dihydrocorynantheal (**18**), and cinchonamine (**19**) were tested as substrates with the oxidase gene candidates as described above, but no activity could be detected. Therefore, to pinpoint the stage at which methoxylation occurs, we performed untargeted and targeted metabolomics analyses of young tissue from *C. pubescens* leaves, stem, and root. These analyses revealed the presence of the putative intermediates tryptamine (**10**), strictosidine (**12**), corynantheal (**13**), dihydrocorynantheal (**18**), cinchoni(di)none (**15a/b**) (along with cinchonamine (**19**) and tetrahydroalstonine (**20**), which are not predicted to be on pathway for **1** and **2**) (Figures S2-S7). Notably, for each of these metabolites a compound corresponding to the methoxylated analog was also detected (Figures S2-S7). These observations clearly indicated that two parallel pathways, one with methoxy and one with desmethoxy substrates, operate in *C. pubescens* (Scheme 2). The detection of 5-methoxytryptamine (**21**) in *C. pubescens* suggested that the aromatic hydroxylation occurs either on tryptamine (**10**) or tryptophan (**9**). However, the hydroxylated or methoxylated derivatives of tryptophan (**9**) were not detected in these metabolomic analyses.^[9]^ Additionally, no orthologs of any tryptophan-5-hydroxylase could be detected in the *C. pubescens* transcriptome. Therefore, we speculated that 5-methoxytryptamine (**21**) derives from hydroxylation of tryptamine (**10**) to form serotonin (**27**), which is then *O*-methylated to form **21** (Scheme 2).

**Scheme 1.**
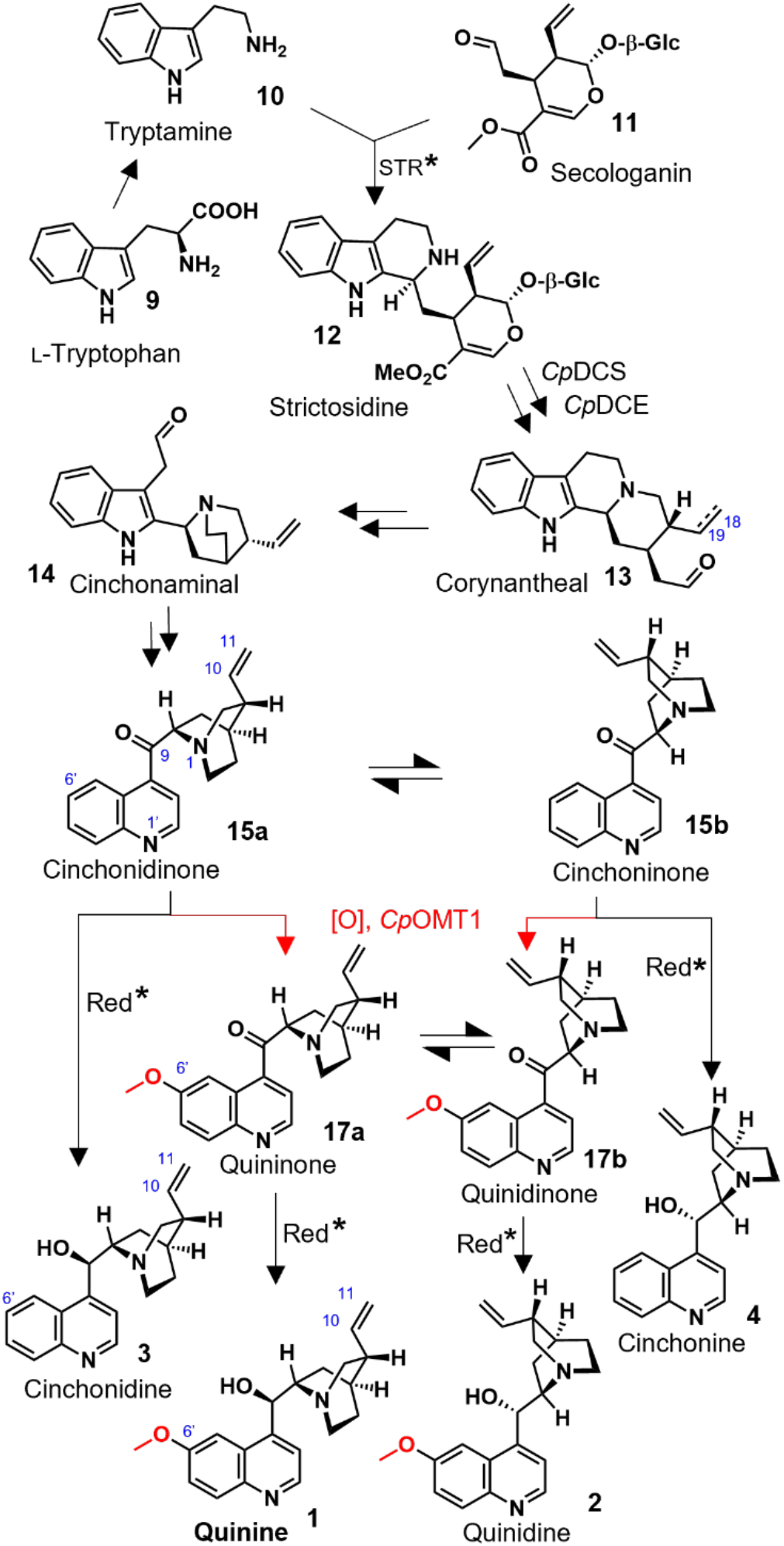
Currently accepted hypothetical biosynthetic pathway to quinine (**1**) and congeners **2-4**. Highlighted in red are the putative steps installing the methoxy group. The asterisk designates previously reported proteins that have been biochemically characterized from crude *Cinchona* tissues, but the corresponding gene remains undiscovered. The dashed double bond at C18-C19 in **13** indicates that this compound also occurs in the dihydro form (i.e., with a single bond, see **18**).

**Scheme 2.**
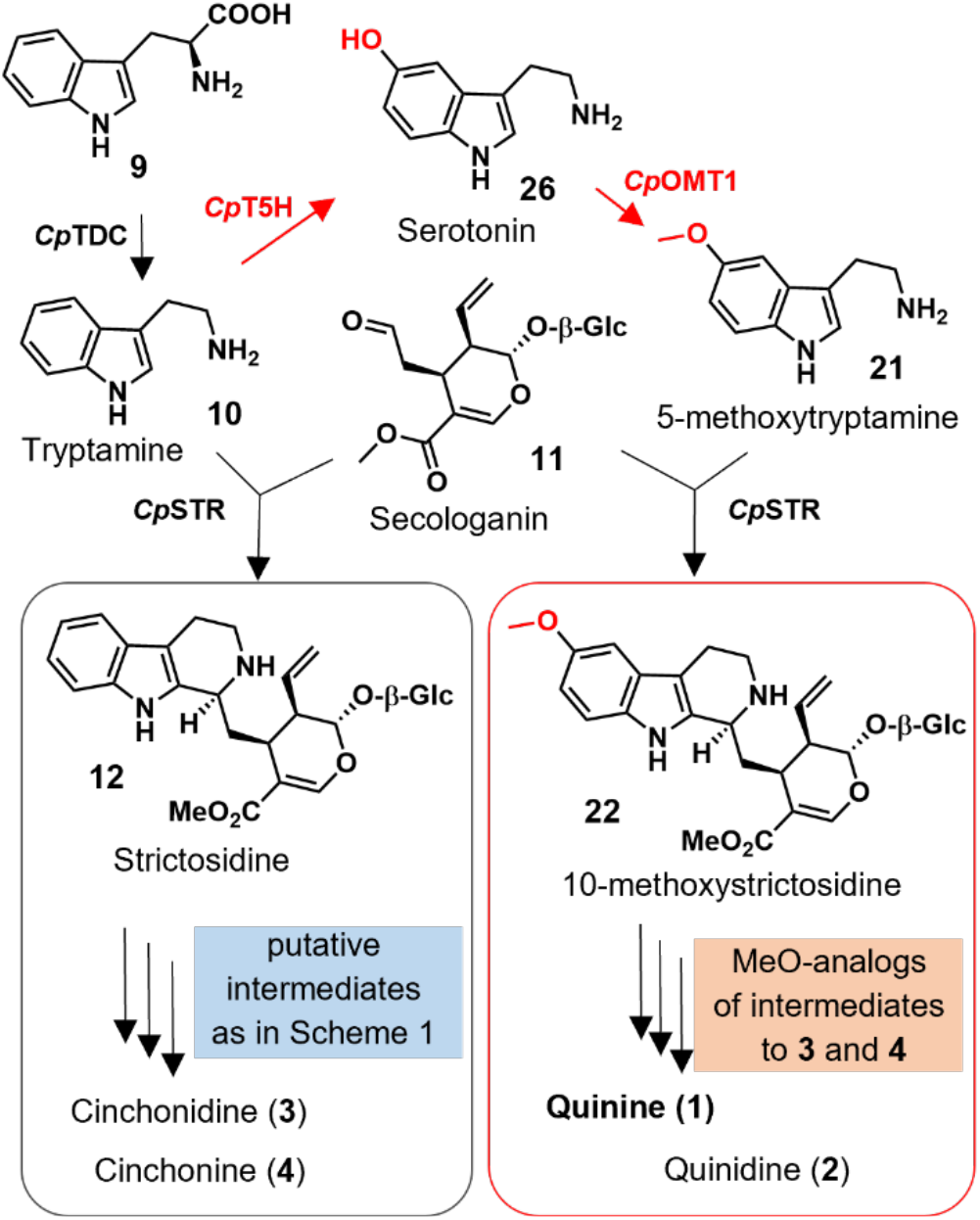
Revised biosynthetic pathway for Cinchona alkaloids. The key steps and involved enzymes that introduce the aromatic methoxy group and leading to parallel pathways are highlighted in red. For a detailed scheme see Scheme S1.

To identify *C. pubescens* enzyme candidates that catalyze formation of 5-hydroxytryptamine (**27**) from tryptamine (**10**), we BLAST-searched for orthologs of CYP71P1 from *Oryza sativa*, the only reported functionally characterized tryptamine-5-hydroxylase.^[10]^ We transiently expressed the 15 top candidates in *N. benthamiana* leaves and infiltrated tryptamine substrate **10** into these infected leaves. LC–MS analysis of the resulting leaf extracts indicated the formation of **27** (*m/z* 177.102 [M+H]^+^) only in *N. benthamiana* leaves expressing either of two highly similar candidate enzymes (97.9% identical amino acids, Figure S8), which we thus named *Cp*T5H1 and *Cp*T5H2 (Figure 1a). Notably, *Cp*T5H1 and *Cp*T5H2 have complementary gene expression profiles (Figure 1b), consistent with the presence of different methoxylated alkaloid types (quinoline-, indole-, yohimbine-, and corynanthe-type) across different *C. pubescens* tissues (Figures S2-S7, and S9).

**Figure 1.**
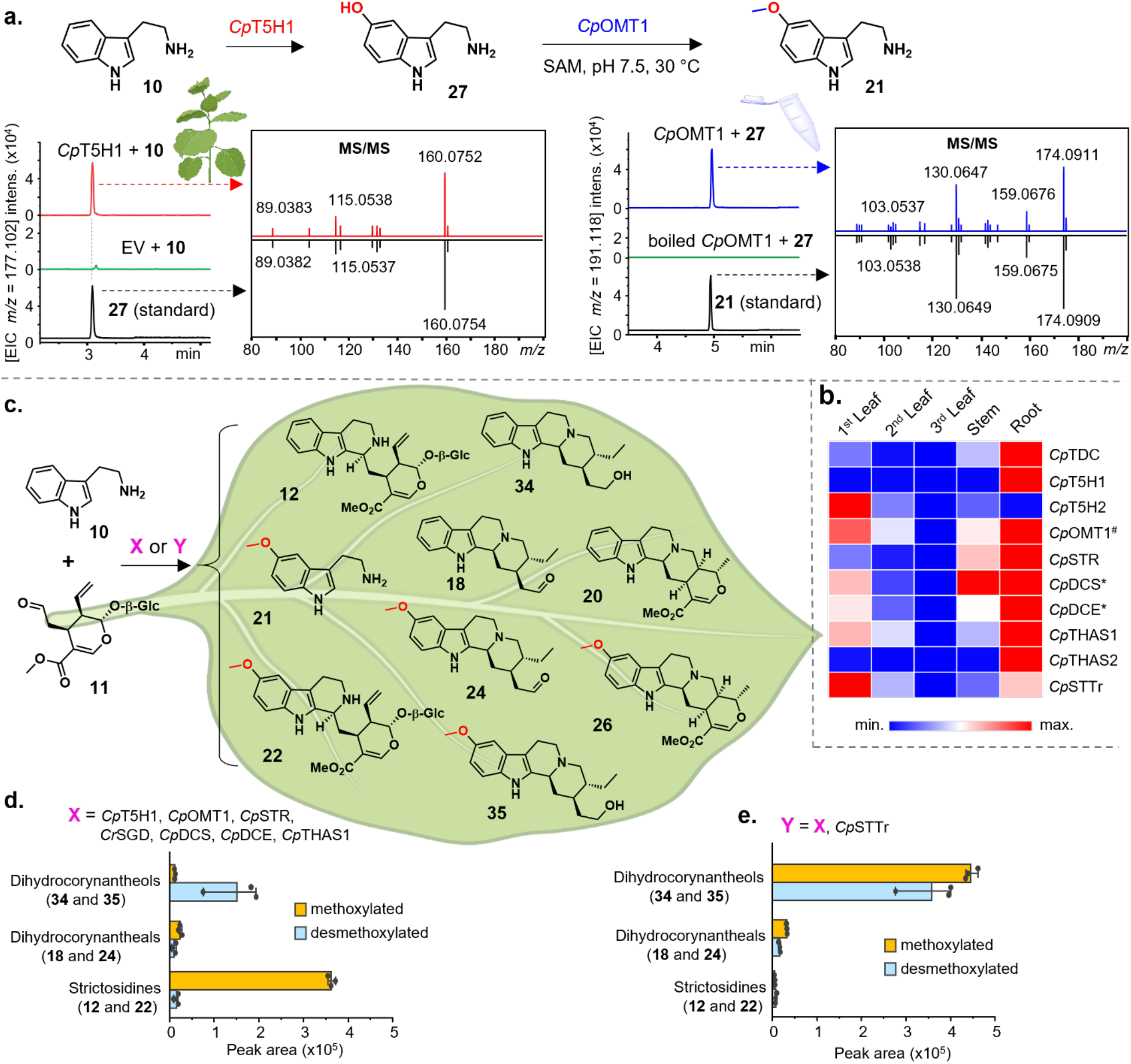
**(a)** Identification and functional characterization of 5-methoxytryptamine (**21**) biosynthetic genes; **(b)** expression profiles of identified genes in *C. pubescens* (^**\***^ indicates previously reported genes; ^#^ denotes previously reported but here functionally repurposed gene); **(c)** reconstruction of the pathways to early Cinchona alkaloids in *N. benthamiana* (for further details see Figure S16); **(d)** peak areas of key metabolites, strictosidines and corynantheal-type alkaloids, produced when only biosynthetic genes are co-expressed in *N. benthamiana* leaves; and **(e)** accumulation of the same key metabolites when the transporter gene *Cp*STTr is co-expressed with the biosynthetic genes in *N. benthamiana* leaves.

Subsequent *O*-methylation of serotonin (**27**) is required to install the methoxy group on 5-methoxytryptamine (**21**). To identify candidate enzymes that methylate the hydroxyl function of **27**, we searched for transcripts annotated as hydroxyindole-*O*-methyltransferase genes in the *C. pubescens* transcriptome, focusing on those that were expressed in all tissues (FPKM > 4) and having a similar co-expression profile with either *Cp*T5H1 or *Cp*T5H2 (Pearson correlation coefficient > 0.5). From this process, seven candidates were identified, expressed in *Escherichia coli*, purified, and assayed *in vitro* using **27** as a substrate. Among these candidates was the previously^[8d]^ identified 6′-hydroxycinchoninone-*O*-methyltransferase (*Cp*OMT1) mentioned above. Surprisingly, only *Cp*OMT1 showed methyltransferase activity with **27**, and the methylated product was identical to an authentic standard of 5-methoxytryptamine (**21**) (Figure 1a). This enzyme more efficiently consumed 5-hydroxytryptamine (**27**) than 6′-hydroxycinchoninone (**16a/16b**) (substrate conversion after 60 min: > 80% and < 20%, respectively, Figure S10). This finding was corroborated by assays using transient expression in *N. benthamiana*, where **27** was almost fully converted to **21** after 3h, while no methylation activity on 6′-hydroxycinchoninone (**16a**/**b**) could be detected, even after 2d. Together, these data suggest that **27** is the native substrate for *Cp*OMT1. We further tested *in planta* the catalytic activity of *Cp*OMT1 on two other hydroxy-indole substrates, 5-hydroxytryptophan and *N*-acetyl-serotonin. We also tested the phenolic substrates quercetin-3-β-glucoside, quercitrin and caffeic acid, since *C. pubescens* also produces a wide range of phenolic natural products (Figure S11). Only *N*-acetyl-serotonin was methylated by *Cp*OMT1 to yield melatonin (Figure S12a), though since this compound was not detected in *C. pubescens*, this enzyme activity may not be physiologically relevant. Moreover, in substrate competition assays, *Cp*OMT1 preferred **27** over *N*-acetyl-serotonin (65% and 40% of conversion, respectively, Figure S12b). Finally, *Cp*OMT1 appears to be evolutionarily related to aromatic *O*-methyltransferases from other monoterpenoid indole alkaloid pathways (Figure S13). We henceforth reclassify *Cp*OMT1 as a serotonin-*O*-methyltransferase.

We then characterized how *Cp*T5H1/2 and *Cp*OMT1 impact the ratio of methoxylated to non-methoxylated products when reconstituted with previously reported downstream *Cinchona* pathway enzymes *in planta*. Specifically, we intended to determine if downstream enzymes *Cp*DCS and *Cp*DCE, which produce intermediate dihydrocorynantheal (**18**) from strictosidine aglycone,^[8d]^ could produce both methoxy and desmethoxy products. However, while previous feeding studies have shown that many downstream monoterpene indole alkaloid pathway enzymes turn over substituted tryptamine analogs,^[11]^ known strictosidine synthase (STR) enzymes do not turn over tryptamine substrates with substituents at the C-5 position.^[12]^ Rationalizing that the *C. pubescens* strictosidine synthase must be able to turn over both tryptamine and 5-methoxytryptamine, we identified and functionally characterized strictosidine synthase from *C. pubescens*, (*Cp*STR) (Figure S14). We also identified two off-pathway enzymes (*Cp*THAS1 and *Cp*THAS2) that reduce the reactive strictosidine aglycone into tetrahydroalstonine **20** (Figure S15).

We transiently co-expressed *Cp*T5H, *Cp*OMT1, *Cp*STR, *Cp*DCS, *Cp*DCE, *Cp*THAS1, and *Catharanthus roseus* strictosidine glucosidase (*Cr*SGD, previously^[13]^ shown to accept 10-methoxy strictosidine) in *N. benthamiana* leaves along with co-infiltration of tryptamine (**10**) and secologanin (**11**). LC-MS analysis of the transformed leaf extracts showed the presence of 5-methoxytryptamine (**21**) in addition to the downstream products strictosidine (**7**), dihydrocorynantheal (**18**), tetrahydroalstonine (**20**), along with the respective methoxylated analogs (Figures 1c and S16). We also noticed that the aldehydes dihydrocorynantheal (**18**) and methoxy dihydrocorynantheal (**24**) were reduced into the corresponding alcohols **34** and **35** by endogenous tobacco reductases (Figure S17). While these results demonstrated that *Cp*STR and all downstream enzymes could turn over both methoxy and desmethoxy intermediates, we noted that all observed dihydrocorynantheal-type products (**18, 24, 34**, or **35**) were produced in low amounts (Figure 2d). In contrast, 10-methoxystrictosidine (**22**) and strictosidine (**12**) accumulated to higher levels (Figure 1d). We hypothesized that 10-methoxystrictosidine (**22**) and strictosidine (**12**), which are likely produced in the vacuole,^[14]^ are not efficiently exported from the vacuole of *N. benthamiana*, and thus cannot access the downstream cytosolic enzymes. Therefore, we searched for and identified an ortholog of a *Catharanthus roseus* strictosidine transporter^[15]^ in the *C. pubescens* transcriptome (*Cp*STTr), which, when co-expressed in *N. benthamiana* leaves along with other biosynthetic enzymes, improved the metabolic flux of both strictosidine (**12**) and 10-methoxystrictosidine (**22**), as evidenced by the relatively higher levels of downstream products, most notably 10-methoxydihydrocorynantheol (**35**) (Figure 1e).

**Figure 2.**
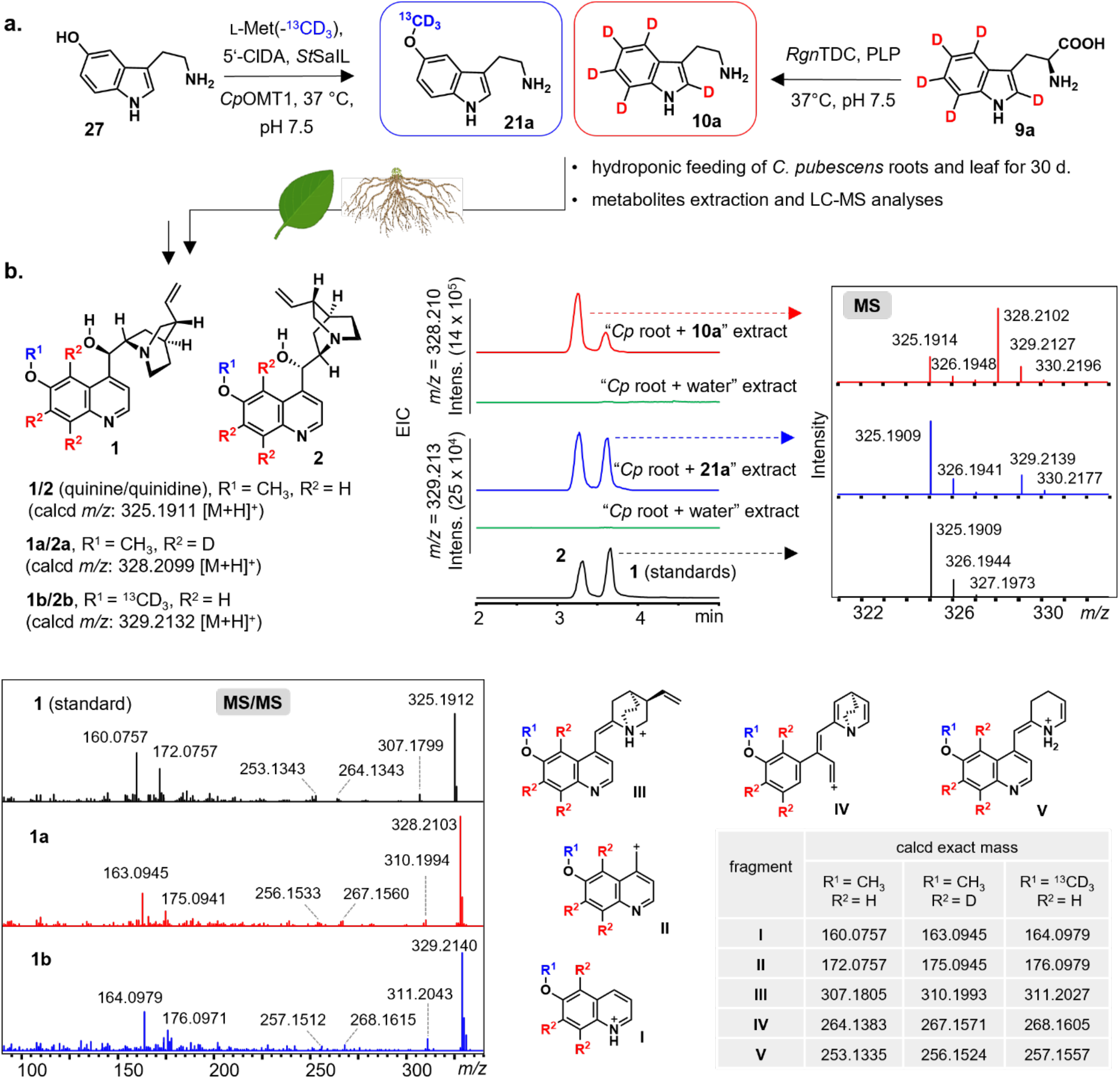
**(a)** synthesis of the labeled precursors tryptamine-(*indole*-*d*5) (**10a**) and 5-methoxytryptamine-(O-*methyl*-^13^C, *d*3) (**21a**), which were separately fed to the roots and leaves of *C. pubescens*; **(b)** key methoxylated Cinchona alkaloids **1** and **2**, and extracted ion chromatograms of the corresponding isotope-labeled analogs, MS isotopic patterns, MS/MS spectrum (20.0-50.0 eV), and selected putative MS/MS fragments **I-V**, all evidencing the incorporation of **21a** and **10a**.

Methoxylated alkaloids appear to be present in larger quantities than desmethoxy alkaloids in *Cinchona* mature plants.^[3a, 16]^ We also observed substantial overproduction of methoxylated products compared to desmethoxy products in *N. benthamiana* reconstitution, though only when the transporter *Cp*STTr was included. The observed ratios of combined production of methoxylated aldehyde **24** and corresponding alcohol **35** versus combined formation of their desmethoxylated counterparts **18** and **34** were 1.23 and 0.23 with and without the transporter, respectively. This indicates that *Cp*STR, *Cp*DCS, and *Cp*DCE are not specific for desmethoxylated substrates. This is particularly remarkable for *Cp*STR, since orthologs from *C. roseus* (*Cr*STR) and *Rauvolfia serpentina* (*Rs*STR) are highly specific for tryptamine.^[17]^ We separately tested *Cp*STR in *N. benthamiana* with tryptamine (**10**), 5-methoxytryptamine (**21**) and 5-hydroxytryptamine (**27**) confirming that this enzyme turns over all of these substrates (Figures S18 and S19). This is consistent with early studies using enzyme activity purified from *Cinchona robusta* suspension culture.^[18]^ Notably, it has been shown that a single point mutation enables *Cr*STR to accept C5-substituted tryptamine analogs,^[12]^ but this mutation is not observed in *Cp*STR. Instead, we noted that *Cp*STR is phylogenetically distant to *Cr*STR and *Rs*STR (Figure S20), which are part of alkaloid pathways where the aromatic methoxylation occurs at a late biosynthetic stage, after formation of strictosidine.

We next wanted to conclusively demonstrate that hydroxylation at this early biosynthetic stage actually occurs in *C. pubescens*, particularly since *Cp*OMT1 can also methylate 6′-hydroxycinchoninone (**16a/16b**), albeit at low efficiency. Therefore, we synthesized isotopically labeled tryptamine-(*indole*-*d*5) (**10a**) and 5-methoxytryptamine-(O-*methyl*-^*13*^*C, d*3) (**21a**) and administrated these two compounds separately to roots and leaf tissues from *in vitro* generated *C. pubescens* plantlets (Figure 2a). Gratifyingly, both labeled substrates were incorporated into quinine (**1**) and quinidine (**2**) (Figure 2b) along with other related alkaloids (Figures S21-S29), as evidenced by LC-MS analysis. In particular, the incorporation of 5-methoxytryptamine (**21**) into **1** and **2**, along with the detection of 5-methoxytryptamine-(*indole*-*d*4) **21b** in the feeding experiments with labeled tryptamine (**10a**) (Figure S25), firmly demonstrated that **1** and related methoxylated metabolites derive from 5-methoxytryptamine (**21**), which, in turn, is formed from tryptamine (**10**) (Schemes 2 and S1).

Since the desmethoxylated alkaloid corynantheal (**13**) had been suggested to be an intermediate of the methoxylated metabolites **1** and **2**,^[8b]^ we also wanted to conclusively determine whether **13** would be incorporated using our established system for feeding studies. Since labeled **13** was not available, we enzymatically synthesized an aromatic ring-*d*4 labeled form of the dihydro analog **18** (i.e., **18a**, Figure S30), to track its incorporation into the downstream dihydro quinine-like compounds present in *Cinchona*. Feeding tissues of *Cinchona* plantlets with **18a** followed by LC-MS analyses of the resulting extracts revealed that **18a** was incorporated into dihydrocinchonidine and dihydrocinchonine (**7** and **8**) (Figure S30). Importantly however, there was no observable conversion of **18a** into the dihydro methoxylated analogs **5** and **6**. Moreover, the labeled isotopic pattern of **18a** was observed in a metabolite tentatively assigned as dihydrocinchonamine **32** (a mid-pathway biosynthetic shunt alkaloid, Figure S31). Hence, this experiment corroborated the finding that methoxylation occurs prior to the formation of **18** ((dihydro)corynantheal) and **32** ((dihydro)cinchonaminal)). In addition, these feeding studies also suggest that the biogenic relationship between **1-4** and the respective dihydro analogs **5-8** diverge early in the pathway, most likely by equilibration of the cyclized forms of strictosidine aglycone as previously^[8d]^ proposed.

In conclusion, metabolomics, transcriptomics and feeding experiments using stable isotope-labeled precursors demonstrate that the biosynthesis of quinine (**1**) and related methoxylated Cinchona alkaloids are initiated by the hydroxylation of tryptamine, followed by *O*-methylation to produce 5-methoxytryptamine. The co-occurrence of both tryptamine and 5-methoxytryptamine *in planta* and the promiscuity of the pathway enzymes drive concomitant parallel formation of methoxylated and desmethoxylated alkaloids in *C. pubescens* by the same pathway enzymes. Along with the discovery of improved product formation gained by inclusion of a vacuolar transporter in reconstitution experiments, this work sets the stage for production of valuable Cinchona alkaloids via synthetic biology.

## Supporting Information

The Supporting Information contains detailed methods and materials. Likewise contained are supplementary scheme, figures and data including chemical structures, LC-MS results, gene nucleotide sequences, phylogenetic analysis, and characterization data of synthesized labeled compounds. The authors have cited additional references within the Supporting Information.^[19-35]^

## Acknowledgements

We are grateful to Dr. Maritta Kunert and Sarah Heinicke for assistance with mass spectrometry, to Jens Wurlitzer for help with some genes cloning, and to Katrin Luck for RNA isolation from roots. Dr. Allwin McDonald is acknowledged for providing *E. coli* mutant harboring *Rgn*TDC. Eva Rothe and the greenhouse team are thanked for taking care of plants. This work was supported by the European Research Council (Grant 788301) and the Max Planck Society. The plant and tissues arts in Figures 1 and 2 were created with BioRender.com. The heatmap in Figure 1 was generated with Morpheus (https://software.broadinstitute.org/morpheus).

