## Supplemental data for "Biosynthetic Origin of the Methoxy Group in Quinine and Related Cinchona Alkaloids"

This file includes:

Material and Methods S2-S12

Supplementary Scheme and Figures S13-S50

MS and MS/MS Spectra of Synthesized Isotope Labeled Compounds S51-S54

Supplementary Tables S55-S62

Supplementary References S63

### Materials and Methods

#### Plant materials

*Cinchona pubescens* plants were germinated and grown on a standard soil mix in a greenhouse using seeds obtained from Edinburgh Botanical Garden, as previously reported.[1] Cultivation temperature fluctuated between 24 and 27 °C, under a 12 h/12 h light/dark cycle. Relative humidity was kept between 70% and 80%. Plants used for the metabolomic and transcriptomic analyses were 1 year old. *Nicotiana benthamiana* plants were cultivated on a standard soil mix in the greenhouse at 22 °C, with a 16 h light and 8 h dark photoperiod and 60% relative humidity. Tobacco plants used in this work were 3-4 weeks old prior to *Agrobacterium tumefaciens* infiltration.

#### Chemicals

All solvents used for extractions, chemical synthesis, and semi-preparative HPLC were of HPLC grade, whilst solvents for UPLC/MS analysis were of MS grade, all purchased from Fisher Scientific. Standards of quinine, quinidine, dihydroquinine, dihydroquinidine, cinchonine, cinchonidine, and tetrahydroalstonine were obtained from Sigma-Aldrich; dihydrocinchonine and dihydrocinchonidine were bought from TCI, and cinchonamine was acquired from ChemSpace. Notably, commercially available cinchonamine was contaminated with its dihydro analog. Cinchona intermediates dihydrocorynantheal, cinchonidinone and its 6-hydroxy- and 6-methoxy analogs were obtained from semi-synthesis as previously described.[1] Standards of strictosidine and its methoxylated congener were readily obtained from enzymatic coupling of secologanin with either tryptamine or 5-methoxytryptamine.[2] The latter three compounds, along with serotonin, and *N*-acetylserotonin were purchased from Sigma-Aldrich. Stable-isotope labeled compounds were synthesized as described below in the section *In vitro* Preparation of Labelled Compounds.

RNA extraction reagents: hexadecyltrimethylammonium bromide (CTAB), polyvinylpyrrolidone (PVP40), chloroform – isoamyl alcohol mixture (24:1), *β*-mercaptoethanol, lithium chloride (LiCl), spermidine trihydrochloride, sodium dodecyl sulfate (SDS), diethylpyrocarbonate, and RNaseZap were all purchased from Sigma-Aldrich. Molecular biology grade DMSO and ethanol absolute were purchased from Fisher Scientific. Carbenicillin and gentamycin were purchased from Formedium, while rifampicin was obtained from Sigma Aldrich and spectinomycin from Fisher Scientific.

#### RNA extraction, purification and sequencing

Total RNA from leaves of three different developmental stages {*i.e.*, the youngest leaf pair at the branch tip (denoted as "1st Leaf" in Figure 1b), the following second leaf pair ("2nd Leaf"), and the third leaf pair ("3rd Leaf")} and from stem of *C. pubescens* (3 biological replicates each) were extracted using the RNeasy Plant Mini Kit (Qiagen), following the manufacturer's instructions. RNA from *C. pubescens* roots (biological triplicates) was extracted following the CTAB protocol, as previously reported.[1] The quality and quantity of obtained RNA were first assessed using an Implen NanoPhotometer® N60. Wherever necessary, RNA was further cleaned up and/or concentrated using the RNA Clean & Concentrator-5 Kit (Zymo Research) according to the manufacturer's instructions. Ultimately, RNA quantitation and integrity were assessed on a 2100 Bioanalyzer system (Agilent) and high-quality samples were submitted to BGI for sequencing, assembly, and annotation.

#### cDNA library preparation and genes amplification

DNAse I-treated RNA from root, stem and youngest leaves of *C. pubescens* were reverse-transcribed using the Superscript IV VILO Master Mix kit (Thermo Fisher). The generated cDNA libraries were aliquoted in Eppendorf tubes and stored at – 20 °C and used to amplify all genes and fragments reported in this work (Table S1). Primers, including overhangs compatible with the appropriate vector on the 5' and 3' end (Table S2), were synthesized by Sigma-Aldrich and gene amplifications were performed with Platinum SuperFi PCR Master Mix polymerase (Thermo Fisher) using MiniAmp Plus Thermal Cycler (Thermo Fisher). Following PCR amplification, DNA products were mixed with TriTrack DNA loading dye (Thermo Fischer), resolved on 1% agarose gel electrophoresis, and visualized on an iBright 1500 Imaging system (Thermo Fischer). DNA fragments of the expected correct size were excised and purified using Zymoclean Gel DNA Recovery Kit (Zymo Research). All kits were used according to manufacturer's instructions.

#### Cloning of gene candidates

The full-length sequences of genes designed for transient expression in *N. benthamiana* were amplified and purified as above described and were ligated into BsaI-linearized (Thermo Fischer) 3Ω1 vector[3] by In-Fusion cloning (Clontech Takara). The In-Fusion reaction was chemically transferred into competent *E. coli* TOP10 cells (Thermo Fischer), which were selected on LB agar supplemented with spectinomycin (200 µg/mL) and incubated at 37 °C overnight. Positive *E. coli* transformants were identified by colony PCR using 3Ω1-specific sequencing primers (Table S4) and were cultivated overnight in LB medium with spectinomycin (37 °C, 225 rpm). Plasmids from positive transformant biomasses were isolated using Wizard Plus SV Minipreps DNA Purification Kit (Promega) and the identities of the inserted sequences were verified by Sanger sequencing (Azenta Life Sciences). The constructs were then used to transform electrocompetent cells of *Agrobacterium tumefaciens* strain GV3101 by electroporation using a MicroPulser™ (BioRad). Recombinant colonies were selected on LB agar containing rifampicin (50 µg/mL), gentamycin (50 µg/mL) and spectinomycin (200 µg/mL) and plates were kept at 28 °C for 2-3 days. Single *A. tumefaciens* colonies were grown in 7-10 mL of LB (50 µg/mL rifampicin, 50 µg/mL gentamycin, and 250 µg/mL spectinomycin) at 28 °C and 250 rpm for up to 1 day. Glycerol stocks of the culture (50%) were thereafter prepared, snap frozen in liquid nitrogen, and stored at −80 °C for further use.

For gene expression in *Escherichia coli*, purified PCR amplicons of genes of interest were inserted downstream of a $His_6$-coding sequence of pOPINF vector[4] linearized with HindIII and KpnI (New England Biolabs) using In-fusion kit. In-fusion assemblies were transformed into competent *E. coli* TOP10 and recombinant colonies were selected on LB agar plates supplemented with carbenicillin (100 µg/mL). Plasmids from positive clones, as identified by colony PCR, were verified by Sanger sequencing. Sequence verified constructs were then used to transform *E. coli* BL21 (DE3) (ThermoFisher) expression cells by heat shock. Positive transformants were confirmed by colony PCR and were then used to inoculate 5-10 mL liquid LB medium (containing 100 µg/mL carbenicillin), which were grown overnight at 37 °C, 200 rpm. These cultures were used to make 50% glycerol stocks, stored at -80 °C.

#### Transient expression of candidate genes in *N. benthamiana*

Transient expression of gene candidates in *N. benthamiana* was performed following the Hawes *et al* protocol,[5] with small modifications. Briefly, A*grobacterium* strains harboring gene constructs of interest were grown from glycerol stocks in 10 mL liquid LB medium (20 µg/mL rifampicin, 30 µg/mL gentamicin, 200 µg/mL spectinomycin) overnight at 28 °C, 250 rpm. Afterwards, cells were harvested by centrifugation at 2,000 *g* for 20 min and the depleted media was discarded. The cell pellets were gently resuspended in 10 mL of infiltration buffer (50 mM MES, 2 mM $Na_3PO_4$, 27.8 mM glucose, 10 mM $MgCl_2$, and 200 µM acetosyringone) and washed to remove residual culture media and antibiotics. Washed cells were recovered by centrifugation at 2,000 *g* for 20 min and were resuspended in 10 mL of infiltration buffer. These preparations were then diluted to an $OD_{600}$ = 0.4-0.5 when a single gene construct was to be tested. When multiple gene constructs were tested in combinations, strains were pooled in equal cell density so that each strain had an $OD_{600}$ of 0.3. The resulting diluted suspensions were incubated in the dark at room temperature with gentle rocking for 1 h and then infiltrated into the abaxial side of *N. benthamiana* leaves covering the entire leaf using a 1-mL needleless syringe. For each *N. benthmiana* plant, only the second pair of fully expanded leaves (counting from the apical meristem) were used for the infection with *Agrobacterium*. After 3 days, 100-200 µM of substrates (cinchonidinone, cinchodinine, cinchonine, strictosidine, cinchonamine, dihydrocorynantheal, and tryptamine) in water with 1 % DMSO were infiltrated into the underside side of previously *Agrobacterium*-infiltrated leaves. The substrate-infiltrated leaf area was marked. 48 h post substrate infiltration, five 10-mm leaf disks were cut from the previously marked parts of the assay leaves and were snap-frozen in liquid nitrogen and stored at -80°C. Each individual infiltration experiment was tested at least 2 times, with biological replicates consisting of at least two leaves from two different tobacco plants.

#### Extraction of metabolites from harvested *N. benthamiana* leaf material

Harvested, snap-frozen *N. benthamiana* leaf disks (100 mg) were ground to a fine powder on a TissueLyser II (Qiagen) using 2 x 2 mm-diameter tungsten beads while shaking vigorously at 22 Hz for 2 min. MeOH (200 µL) was added to each sample and the suspensions were vigorously vortexed for 30 s. After a short centrifugation (10000 *g*, 1 min), the samples were sonicated at room temperature for 15 min. After sonication, samples were centrifuged at full speed (>13000 *g*, 5 min) and filtered through 0.22 µm PTFE syringe filters.

#### Expression and purification of proteins in *E. coli*

For production of recombinant enzymes in *E. coli*, aliquots from glycerol stocks prepared as described in section **Cloning methods** were inoculated in 10 mL LB medium (100 µg/mL carbenicillin) and grown at 37 °C, 200 rpm overnight. The overnight cultures (10 mL) were next used to inoculate 250-500 mL of 2xYT medium (100 µg/mL carbenicillin) to an $OD_{600}$ of about 0.1. The resulting large cultures were further grown at 37 °C, 200 rpm and when they reached $OD_{600}$ of 0.7, they were chilled on ice for 15 min. Thereafter, 0.3 mM isopropyl β-D-1-thiogalactopyranoside (IPTG) were added to induce gene expression. The induced cultures were incubated at 18 °C, 200 rpm. After 16-18 h of incubation, microbial cells were harvested by centrifugation at 4,000 rpm, 4 °C, for 20 min resuspended in 15-30 mL of pre-cooled buffer A (50 mM Tris-HCl pH 7.4, 50 mM glycine, 500 mM NaCl, 5% glycerol, 20 mM imidazole) supplemented with 10 mg lysozyme and 1 tablet of Complete EDTA-free protease inhibitors (Roche Diagnostics), and incubated on ice for 30 min. The resuspended cells were disrupted by sonication using an ultrasonic liquid processor (vibra cell™, Sonics®; 40 % amplitude; 2s on/3s off; total 'on'-time: 3 min). Cell debris were pelleted by centrifugation at 35,000 *g* for 30 minutes. The supernatant was filtered through syringe glass-filter (Sartorius) and 200-300 µL of His60 Ni Superflow Resin (Taka Bio) were added. Prior to their addition, the resin beads were washed three times with 2 mL of ice-cooled A1 buffer. The suspensions were incubated for 30 min under a gentle shaking in the cold room. After incubation, resin beads were pelleted and washed three times with buffer A1. Elution of the proteins was performed twice by resuspending the beads in 600 µL Buffer B (50 mM Tris-HCl pH 8; 50 mM glycine; 500 mM NaCl; 5% glycerol; 500 mM imidazole). Dialysis and buffer exchange was performed using Buffer A4 (20 mM HEPES pH 7.5; 150 mM NaCl) in centrifugal concentrators (Amicon® Ultra Centrifugal Filter, Merck Millipor). Protein concentrations were measured on a nano-photometer (Implen), using the absorbance at 280 nm and theoretical extinction coefficients. Proteins were aliquoted in 100 µL, snap-frozen and stored at -80 ºC.

All recombinant proteins used in this work were produced in this way. However, the expression and purification of *Ruminococcus gnavus* L-tryptophan decarboxylase (*Rgn*TDC) were performed with the following small modifications. The *Rgn*TDC *E. coli* transformant was grown and induced at $OD_{600}$ > 1.0 with 1 mM IPTG and 0.5 mM indole. And the lysis solution (buffer A) contained in addition 150 µM pyridoxal 5′-phosphate (PLP), a colored cofactor useful for monitoring protein elution.

#### Enzymatic assays for *O*-methyltransferase activity

Recombinant OMT candidates were assessed using serotonin as substrate. Reaction mixtures (100 µL, total volume) were composed of 2 µM of the respective OMT candidate, 100 µM SAM, 50 mM HEPES (pH 7.5), and 60 µM serotonin. The reactions were incubated at 30 °C for 1 h and were quenched with 200 µL of methanol. Negative controls consisted of boiled OMT in the reaction mixture. The quenched reaction mixtures were centrifuged at 15000 *g* for 2 min and filtered through 0.22 µm PTFE syringe filters and analyzed by LC/MS-QTOF using Method 2 when serotonin was a substrate or Method 1 when 6'-hydroxycinchoninone was used. OMT candidate genes were also assayed in *N. benthamiana*.

#### Assays of pathway reconstruction in *N. benthamiana*

*Agrobacterium* strains harboring gene constructs of *Cp*T5H-1, *Cp*OMT1, *Cp*STR, *Cp*TAS1, *Cp*DCS, *Cp*DCE, *Cp*STTr, *Cr*SGD and an empty vector were separately cultivated and prepared as described above. Two combinations were made from these cultures. The first combination contained *Cp*T5H-1, *Cp*OMT1, *Cp*STR, *Cp*TAS1, *Cp*DCS, *Cp*DCE, *Cr*SGD and the empty vector strain, each pooled to have a final $OD_{600}$ of 0.3. In the second pool, the empty vector strain was substituted by the strain harboring *Cp*STTr. The two bacterial combinations were used to infiltrated leaves of *N. benthamiana* plants (biological triplicates) from the same batch. After 3 days, a solution of tryptamine and secologanin (each at final concentration of 500 µM) was infiltrated into the underside side of previously *Agrobacterium*-infiltrated leaves and the substrate-infiltrated leaf areas were marked. 48 h post substrate infiltration, the marked parts were harvested, snap-frozen in liquid nitrogen, and homogenized using a TissueLyser II (Qiagen). Aliquots of 200 µL of MeOH were added to each 100 mg of ground tissue. The mixture was then vortexed thoroughly, sonicated in a sonic bath for 15 min, spun down at 13 000 *g* for 2 min and filtered through 0.22 µm PTFE syringe filters. Filtered samples were directly analyzed by LC/MS-QTOF (Method1).

#### Assays of enzyme specificity activity in *N. benthamiana*

For enzyme specificity activity tests using *N. benthamiana* expression system, *Agrobacterium* strains harboring the gene constructs of interest (*Cp*OMT1 or *Cp*STR) were prepared and infiltrated into leaves of *N. benthamiana* plants (biological triplicates) as described above. After 3 days, test substrates were co-infiltrated into the underside side of previously *Agrobacterium*-infected leaves and the substrate-infiltrated leaf areas were marked. In case of *Cp*OMT1, a solution of serotonin and *N*-acetylserotonin (each at final concentration of 60 µM) was used, and for *Cp*STR a solution containing secologanin (120 µM), tryptamine (100 µM) and 5-methoxytryptamine (100 µM) was utilized. 24 h post substrates infiltration, the marked parts were harvested and methanolic extracts were prepared as described above and analyzed by LC-MS (Method 1).

#### LC-MS analysis of assay samples

LC-MS analyses were routinely performed on an UltiMate 3000 ultra-high performance liquid chromatography system (UHPLC; Thermo Fischer) connected to an Impact II UHR-Q-ToF (Ultra-High Resolution Quadrupole-Time-of-Flight) mass spectrometer (Bruker). For all data herein reported from this analytical system, the following conditions were maintained for the mass spectrometer. Ionization was performed in positive mode via pneumatic-assisted electrospray ionization (ESI+) with a capillary voltage of 3500 V, an end plate offset of 500 V, and a nebulizer pressure of 2.5 bar. Nitrogen at 250°C with a flow of 11 L/min was used as the drying gas. Data acquisition was recorded at 12 Hz in a mass range from 80 to 1000 *m/z*, using data-dependent MS/MS, an active exclusion window of 0.2 min and a reconsideration threshold of 1.8-fold change. Fragmentation was triggered on an absolute threshold of 400 and restricted to a total cycle time range of 0.5 s. For collision energy, the stepping option model from 20 to 50 eV was used. Sodium formate solution in isopropanol was used as a calibrant at the beginning and the end of each run. Automatic injection of this calibration solution was operated by an external syringe pump at 0.18 mL/min using a 5 mL syringe with an ID of 10.3 mm. To prevent the spectrometer from salt contamination and avoid injection peak, the initial 1 min of the active chromatographic gradient of each run was directed into waste, after which the sample was directed into the mass spectrometer. Chromatographic resolution conditions were varied according to analyte properties. The following methods were used.

UHPLC method 1

The resolution was performed on a Phenomenex Kinetex XB-C18 (100 x 2.1mm, 2.6 µm; 100 Å) column at 40 °C using a solvent system consisting of MilliQ water with 0.1% formic acid (A) and acetonitrile (B) under the following conditions: 10%B for 1 min, linear gradient from 10% to 30% B in 6 min, 90% B for 1.5 min, and 10% B for 2.5 min. A flow rate of 0.6 mL/min and an injection volume of 2 µL were used. This was the standard method used for analysis of samples from all biochemical assays, unless otherwise specified.

UHPLC method 2

Here samples were run on a Phenomenex Synergi Hydro-RP (150 x 2 mm, 4 µm; 80 Å) column at 25 °C using 5 mM ammonium and 0.01% formic acid (A) and methanol (B) under the following conditions: 1%B for 0.5 min, linear gradient from 1% to 10% B in 1.5 min and from 10% to 50% B in 4 min, 90% B for 1.5 min, and 1% B for 2.5 min. The flow rate was set to 0.6 mL/min and 2 µL of sample were injected. This method was used for the analysis and detection of serotonin **27**.

UHPLC method 3

The column used here was a Waters Acquity UPLC BEH-C18 column (2.1 x 50 mm; 1.7 µm; 130 Å; column temperature 40°C). Samples were run at a flow rate of 0.6 mL/min in an ammonia 0.025%- acetonitrile gradient linearly increasing from 5% to 65% acetonitrile in 17 minutes. 2 µL of samples were injected. This method was used to resolve the aldehyde dihydrocorynantheal (**18**) and its corresponding alcohol **34** (Figure S18). It was also used to differentiate tetrahydroalstonine from its stereioisomers and isobaric compounds.

#### Metabolomic analyses

Sample preparation and LC-MS analyses

Plant material (root, stem, and a young leaf at the apical meristem) from 1-year old *C. pubescens* (two biological replicates) were harvested, wrapped into an aluminum foil and snap frozen in liquid nitrogen. The frozen tissues were next ground with mortar and pestle to a fine powder. Aliquots of tissue powder were extracted with MeOH and sonicated at room temperature for 15 min. The volume of MeOH was normalized on fresh tissue weight (100:1 = mg:mL). The methanolic suspensions were centrifuged at 15000 *g* for 4 minutes and the supernatant filtered through a 0.2 µm PTFE filter. The resulting methanolic filtrates were diluted 10x times and submitted to UHPLC-HRMS analysis, performed on a Vanquish (Thermo Fisher Scientific) system coupled to a Q-Exactive Plus Orbitrap (Thermo Fisher Scientific) mass spectrometer. A Waters Acquity UPLC BEH-C18 column (2.1 x 50 mm; 1.7 µm; 130 Å) at a temperature of 40 °C was used for metabolites resolution. The solvents system was composed of MilliQ water supplemented with 0.1% formic acid (A) and acetonitrile (B). The gradient elution started with 5% B and increased linearly to 30% B over 10 min, and from 30% to 100% B in 1 min. The column wash stage was done with 100% B for 1 min and the conditioning stage at 5% B was performed within 2.5 min. The flow rate was maintained at 0.6 mL/min throughout the run. 2 µL of samples were injected.

The Q-Exactive Plus Orbitrap mass spectrometer (Thermo Fisher Scientific) was equipped with a heated electrospray ionization (HESI) source. The operating parameters of HESI were set according to the UHPLC flow rate of 0.6 mL/min leading to the following source auto default: sheath gas flow rate = 55; auxiliary gas flow rate = 15; sweep gas flow rate = 3; spray voltage = 3.50 kV; capillary temperature = 275 °C; auxiliary gas heater temperature = 450 °C; and S-lens RF level = 50. Data acquisition was carried out in full scan MS mode (resolution 70,000) in positive mode over the mass range *m/z* from 100 to 1,000. The full-scan and data-dependent MS/MS mode (full MS/dd-MS2 Top5) was used to simultaneously record the spectra of the precursors as well as their MS2 fragmentation. The parameters for dd-MS2 were set up as follows: resolution 17,500, mass isolation window 4.0 m/z, and normalized collision energy (NCE) 30%. The format used for generated spectrum data was centroid. The Pierce positive and negative ion mass calibration solution (Thermo Fisher Scientific) was used to calibrate the mass spectrometer.

Authentic standards of cinchona alkaloids (80 nM to 1 uM in 7:3 MeOH/$H_2O$) were also analyzed under the same conditions for an unambiguous assignment through comparison of compound retention time, MS spectrum and MS/MS fragmentation. The data as reported in Figures S1-S5 were obtained from the runs on this LC-Orbitrap system. However, the samples were also analyzed on LC-QTOF (method1) described in LC-MS analysis of Assays Samples section, for comparison of the structural prediction and annotation from MS databases described below.

LC-MS data processing and analyzing

Raw data generated from LC-Orbitrap system were directly imported into Mzmine 4.2.0 software for data processing.[6] Features were detected using the workflow setup mzwizard with the following parameters: Minimum feature height: 500.0; *m/z* tolerance (scan-to-scan): 0.002 *m/z* or 10.0 ppm; *m/z* tolerance (intra-sample): 0.0015 *m/z* or 3.0 ppm; *m/z* tolerance (sample-to-sample): 0.0015 *m/z* or 5.0 ppm; retention time tolerance (intra-sample): 0.4 min; and retention time tolerance (sample-to-sample): 0.1 min. The Mzmine processed features were exported to SIRIUS 5 software.[7] Prediction of compound classes was done with CANOPUS[8] and chemical classification was performed using NPclassifier.[9] In parallel, the Mzmine processed data were also exported to the GNPS platform[10], [11] for feature-based molecular networking analysis[12] and spectral library search. The matches between samples spectra and library spectra were kept only if they had a score above 0.7 and at least 6 matched peaks.

LC-QTOF generated data were analyzed using Bruker Compass MetaboScape 2021b software (version 7.0.1). Peak detection and area quantitation were performed using the T-Rex 3D algorithm and the non-targeted metabolomics workflow with an intensity threshold set to 1000. The generated output, containing a list of mass signatures with retention times, along with qualitative peak intensities by automated integration of extracted ion chromatograms with 5 ppm tolerance and MS/MS spectrum, was exported for further analyzed with SIRIUS and GNPS platform as described above.

All LC-MS-related figures herein presented were prepared in OriginPro 2019 (version 9.6.0) from the Mzime or MetaboScape processed data, and were thereafter arranged in Microsoft Powerpoint.

***In vitro* enzymatic preparation of stable-isotope labeled compounds**

Preparation of tryptamine-(*indole-d*5) (**10a**)

Tryptamine-(*indole-d*5) (**10a**) was obtained by decarboxylation of L-tryptophan-(indole-*d*5) (**9a**), purchased from CDN Isotopes, using *Ruminococcus gnavus* tryptophan decarboxylase (*Rgn*TDC).[13] The reaction mix (total volume 1 mL) contained **9a** (430 µM), pyridoxal 5′-phosphate (PLP, 100 µM), and *Rgn*TDC (5 µM) in HEPES (pH 7.5 50 mM). The reactions were incubated at 37 °C for 20 h and were stopped by adding 1 mL MeOH. The reaction mix were then combined and methanol was evaporated on a rotavapor. The aqueous suspension was passed through reverse-phase solid-phase extraction (SPE) cartridges (Discovery DSC-18, 1g, Supelco) columns, washed with 10 mL water. Tryptamine-(*indole-d*5) (**10a**) was then eluted with 4 mL MeOH and dried in vacuum. Product purity was verified by HPLC-UV and LC/MS-QTOF (Method 1) and structural characterization was done by comparing the MS/MS fragmentation and retention time of the obtained isotope labeled product to the spectrum and retention of an unlabeled authentic standard (elution of the labeled compound is about 0.02 min earlier than the unlabeled analog). HRESIMS *m/z* 166.1380 $[M+H]^+$ (calcd *m/z* for $C_{10}H_8D_5N_2^+$, 166.1387). For MS/MS spectra, see Figure S33.

Synthesis of *d*4-strictosidine (**12a**)

*d*4-Strictosidine (**12a**) was prepared from 9 mM tryptamine-(*indole*-*d*5) (**10a**), 8 mM secologanin (**11**) and 5 µM of recombinant STR from *Catharanthus roseus* (*Cr*STR) in a total volume of 20 mL HEPES buffer (50 mM, pH 7.5), incubated with stirring at 30 °C for 18 h. The reaction products were submitted to SPE columns, thereby removing salts and denaturated protein. Elution with 10 mL methanol provided **12a** accompanied with some impurities. The latter were removed on semi-preparative HPLC (see below), yielding 3 mg of *d*4-strictosidine (**12a**). As for **12a**, the product structure was inferred by comparing the MS/MS fragmentation and retention time of the obtained isotope labeled product to the spectrum and retention of an unlabeled authentic standard (elution of the labeled compound is about 0.02 min earlier than the unlabeled analog). HRESIMS *m/z* 535.2580 $[M+H]^+$ (calcd *m/z* for $C_{27}H_{31}D_4N_2O_9^+$, 535.2588). For MS/MS spectra, see Figure S34.

Preparation of *d*4-dihydrocorynantheal (**18a**)

*d*4-Dihydrocorynantheal was prepared by enzymatic conversion of *d*4-strictosidine (**12a**) with recombinant *Cr*SGD, *Cp*DCS, and *Cp*DCE enzymes. The reaction mix (500 µL) consisted of **12a** (200 µM), *Cr*SGD (4 µM), *Cp*DCS (8 µM), *Cp*DCE (6 µM), NADPH (200 µM) and HEPES (pH 7.5, 50 mM). The reaction mix were incubated at 30 ºC for 2 h. After incubation, the samples were stopped by adding 1 volume of MeOH and vigorously vortexing for 30 sec. After removal of MeOH on rotavapor, the reaction products were placed into SPE cartridges and washed with water. Retained compounds were eluted with 10 mL MeOH, concentrated, and resolved by semi-preparative HPLC (see below), yielding *d*4-dihydrocorynantheal (0.8 mg). As for **10a**, the product structure was inferred by comparing the MS/MS fragmentation and retention time of the obtained isotope labeled product to the spectrum and retention of an unlabeled authentic standard (elution of the labeled compound is about 0.02 min earlier than the unlabeled analog). HRESIMS *m/z* 301.2214 $[M+H]^+$ (calcd *m/z* for $C_{19}H_{21}D_4N_2O^+$, 301.2212). For MS/MS spectra, see Figure S35.

Preparation of 5-methoxytryptamine-(O-*methyl*-$^{13}$C, *d*3) (**21a**)

To obtain 5-methoxytryptamine-(O-*methyl*-$^{13}$C, *d*3) (**21a**), 5′-chloro-5′-deoxyadenosine (160 µM) and L-methionine-(*methyl*-$^{13}$C, *d*3) (400 µM) were first incubated with recombinant *Salinispora tropica* SalL enzyme (10 µM ) in HEPES (pH 7.5, 50 mM) for 10 min at 37 ºC to generate *in situ S*-adenosyl-L-methionine-(*methyl*-$^{13}$C, *d*3).[14] After the 10 min incubation, serotonin **27** (final concentration 250 µM) and recombinant *Cp*OMT1 (final concentration 2 µM) were added, making the reaction total volume 1 mL. The reaction mix were incubated for 3 h, after which MeOH (1 mL) was added and the mix was processed through the SPE workup as described above. The recovered fraction was resolved on a semi-preparative HPLC column (see below), yielding 0.9 mg of **21a**. As for **10a**, the product structure was inferred by comparing the MS/MS fragmentation and retention time of the obtained isotope labeled product to the spectrum and retention of an unlabeled authentic standard (elution of the labeled compound is about 0.02 min earlier than the unlabeled analog). HRESIMS *m/z* 195.1396 $[M+H]^+$ (calcd *m/z* for $C_{10}{}^{13}CH_{12}D_3N_2O^+$, 195.1401). For MS/MS spectra, see Figure S36.

**Purification of compounds by semi-preparative HPLC**

Compound purification was performed on an Agilent 1260 Infinity HPLC system (Agilent Technology), equipped with an automatic sample injection system, a binary pump, an oven, a DAD detector, and a fraction collector. All samples were resolved on a Phenomenex Kinetex XB-C18 (4.6 x 10.0 mm; 5 µm; 100 Å) column operating at 40 ºC, using water with 0.1% formic acid (solvent A) and acetonitrile (solvent B) with the following conditions. A linear gradient of 10% – 20% B over 17 min was used for the separation of *d*4-strictosidine (**12a**) (collection time range: 10.8 – 12.8 min). For the resolution of *d*4-dihydrocorynantheal (**18a**), a linear gradient starting with 5% to 17.4% B over 15 min was utilized (collection time range: 11.2 – 12.8 min). 5-methoxytryptamine-(O-*methyl*-$^{13}$C, *d*3) (**21a**) was obtained from a gradient of 5% – 8.6% B over 5 min (collection time range: 4.5 – 4.9 min). For all separations, the flow rate was 0.8 mL/min and compounds were detected by UV absorption at 4 wavelengths: 195 nm, 214 nm, 254 nm; and 290 nm. Collected fractions were assessed on LC/MS-QTOF and fractions containing pure compound of interest were combined and dried using a Genevac EZ-2 Plus evaporation system. The products were then collected in 2 mL methanol, transferred in a glass vial and dried under nitrogen at room temperature.

**Feeding experiments**

Tryptamine-(*indole*-*d*5) (**10a**) was prepared as a 3.3 mM solution in water with 1% DMSO, and 5-methoxytryptamine-(O-*methyl*-$^{13}$C, *d*3) (**21a**) and *d*4-dihydrocorynantheal (**18a**) were prepared in the same way as 1 mM solution. Feeding studies were performed using tissues from *C. pubescens* plantlets grown *in vitro*. Briefly, *C. pubescens* seeds were germinated under sterile conditions on half Murashige and Skoog medium prepared in Gelrite (Duchefa Biochemie). The grown seedlings developed side shoots, which were cut in single shoots or groups of shoots and placed on McCown Woody plant medium (sucrose: 30 g/L; 6-(γ,γ-dimethylallylamino)-purine (2iP) : 2 mg/L; plant agar: 8 g/L at pH 5.8). The plantlets were sub-cultured every 6 weeks and maintained in a growth chamber at 25 °C, with a 16 h/8 h photoperiod under LED light and 70% humidity. For feeding experiments, young secondary and tertiary roots from a 2-month-old plantlet (two biological replicates) were cut in about 1-cm pieces, which were washed through a short submersion into sterile de-ionized water. 3 to 4 washed root pieces were placed into wells of a 48-well plate containing 100 µL of sterile de-ionized and 20 µL of McCown Woody plant medium. From the same plantlet, the pair of young leaves at the apical meristem were cut. Leaves were first sectioned into apical and base portions. The apices were discarded and the bases were sliced through the midvein. The half leaf bases were placed into separate wells of the abovementioned 48-well plate containing 100 µL of sterile de-ionized water and 20 µL of McCown Woody medium. For each isotope labeled compound, there was one well containing roots and a second well containing a leaf portion. 90 µL of each solution of the isotope labeled compounds were added and the volume in every well was brought to 300 µL with sterile de-ionized water (final concentration for **21a** and **18a**: 300 µM, and for **10a**, 1 mM). Plantlet tissues treated with sterile de-ionized water were used as controls. The 48-well plate with treated and control plant tissues was sealed with paraffin and incubated at 25 °C, with shaking at 80 rpm. After 4 weeks, tissues were collected and ground on a TissueLyser II (Qiagen). Methanolic extracts (100 mg/mL) were prepared and analyzed by LC–MS, as described above.

**Phylogenetic analysis**

Unless specified, phylogenetic trees were generated in MEGA11.[15] The Maximum Likelihood method and Poisson correction model[16] were used to infer the evolutionary history. The percentage of trees in which the associated taxa clustered together is shown next to the branches. Initial tree(s) for the heuristic search were obtained automatically by applying Neighbor-Join and BioNJ algorithms to a matrix of pairwise distances estimated using the Poisson model, and then selecting the topology with superior log likelihood value. The tree is drawn to scale, with branch lengths measured in the number of substitutions per site.

#### Supplementary Scheme and Figures

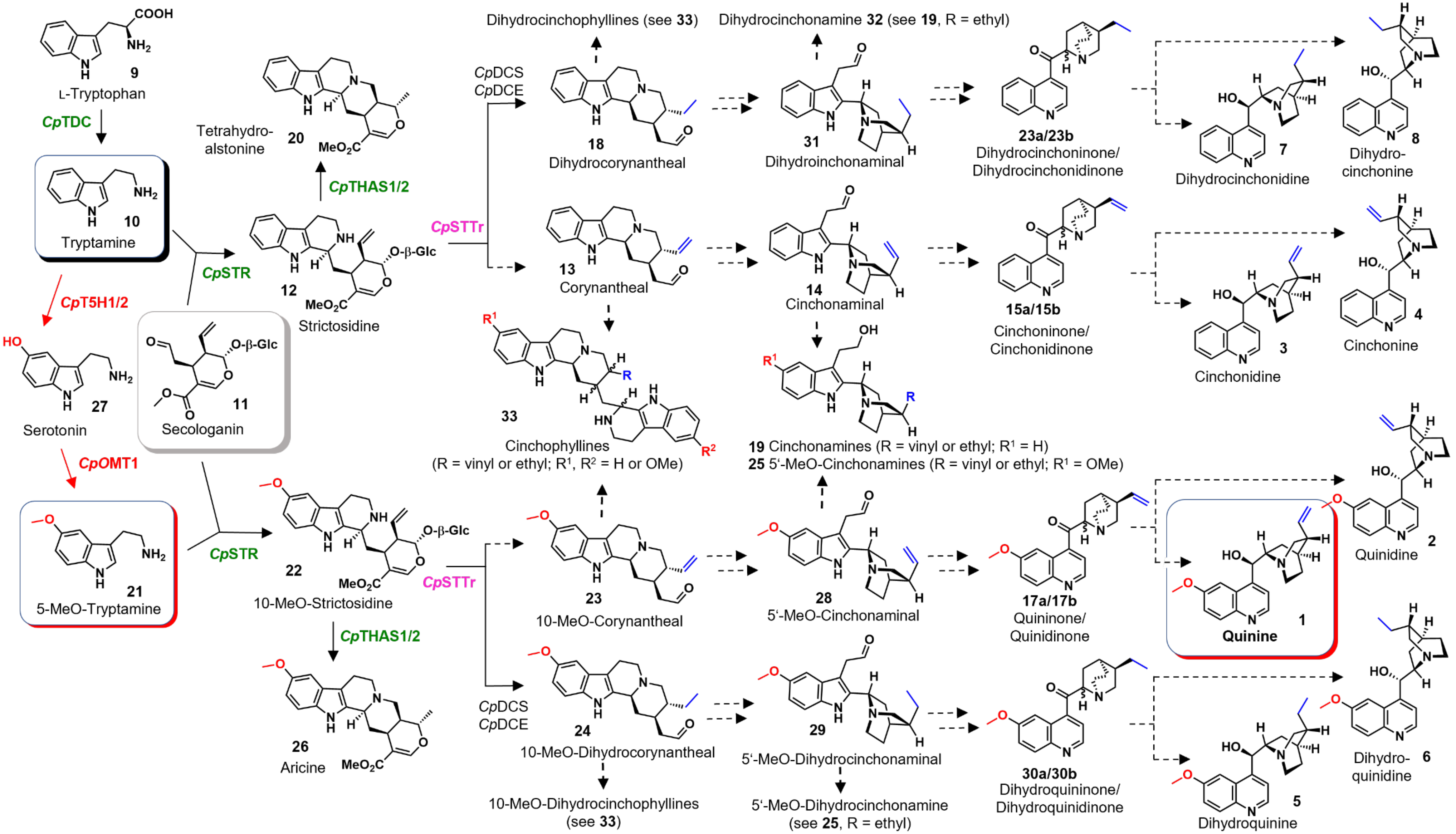


**Scheme S1. Revised proposed pathways to quinine (1) and related Cinchona alkaloids**. The key steps and involved enzymes that introduce the aromatic methoxy group and leading to parallel routes as elucidated in this work are highlighted in red. Other enzymes discovered in this work are highlighted in green (for biosynthetic proteins) and in pink (for substrate transporter protein). Dashed arrows designate steps for which genes are still unknown.

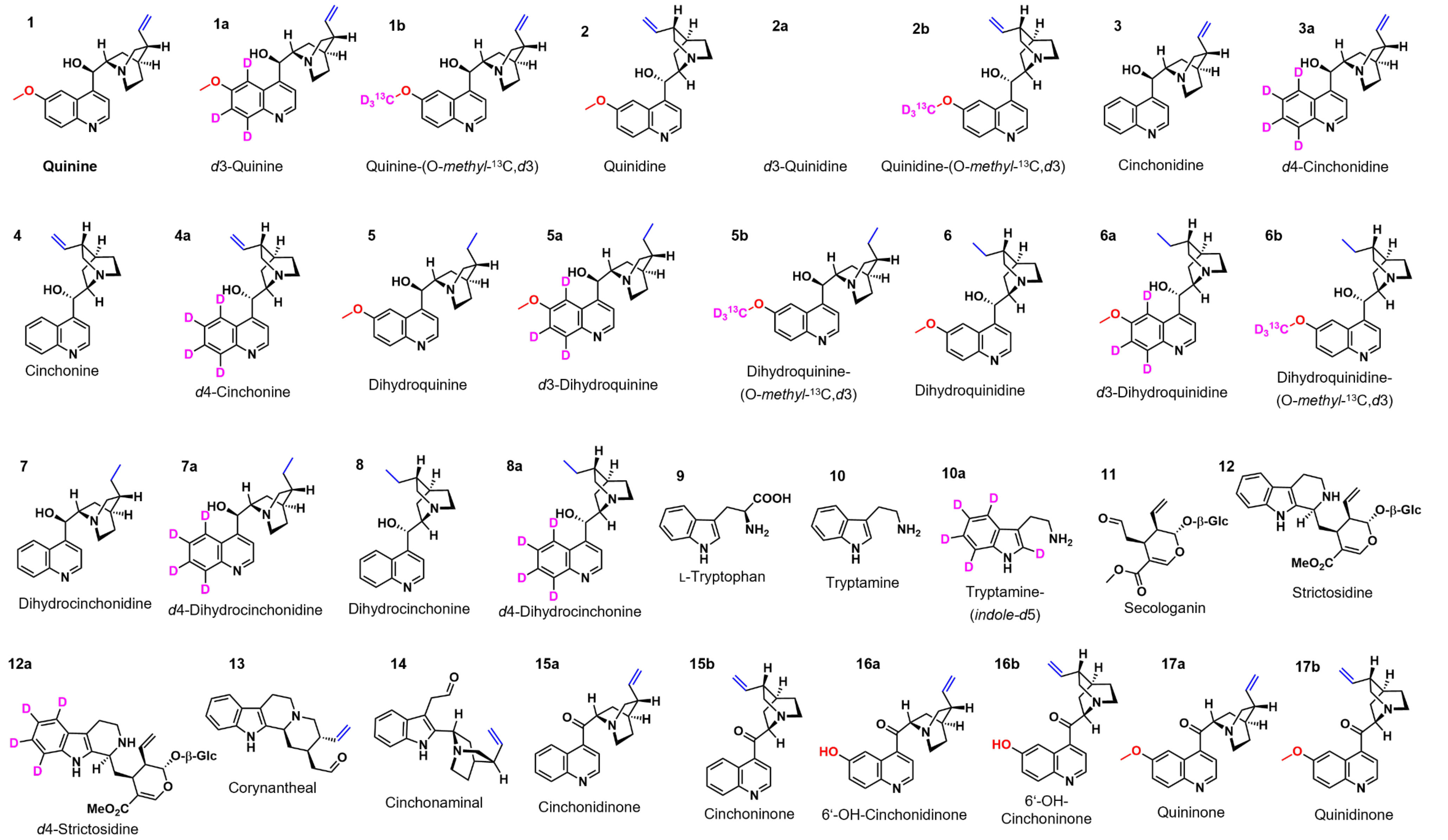


**Figure S1** continues on next page

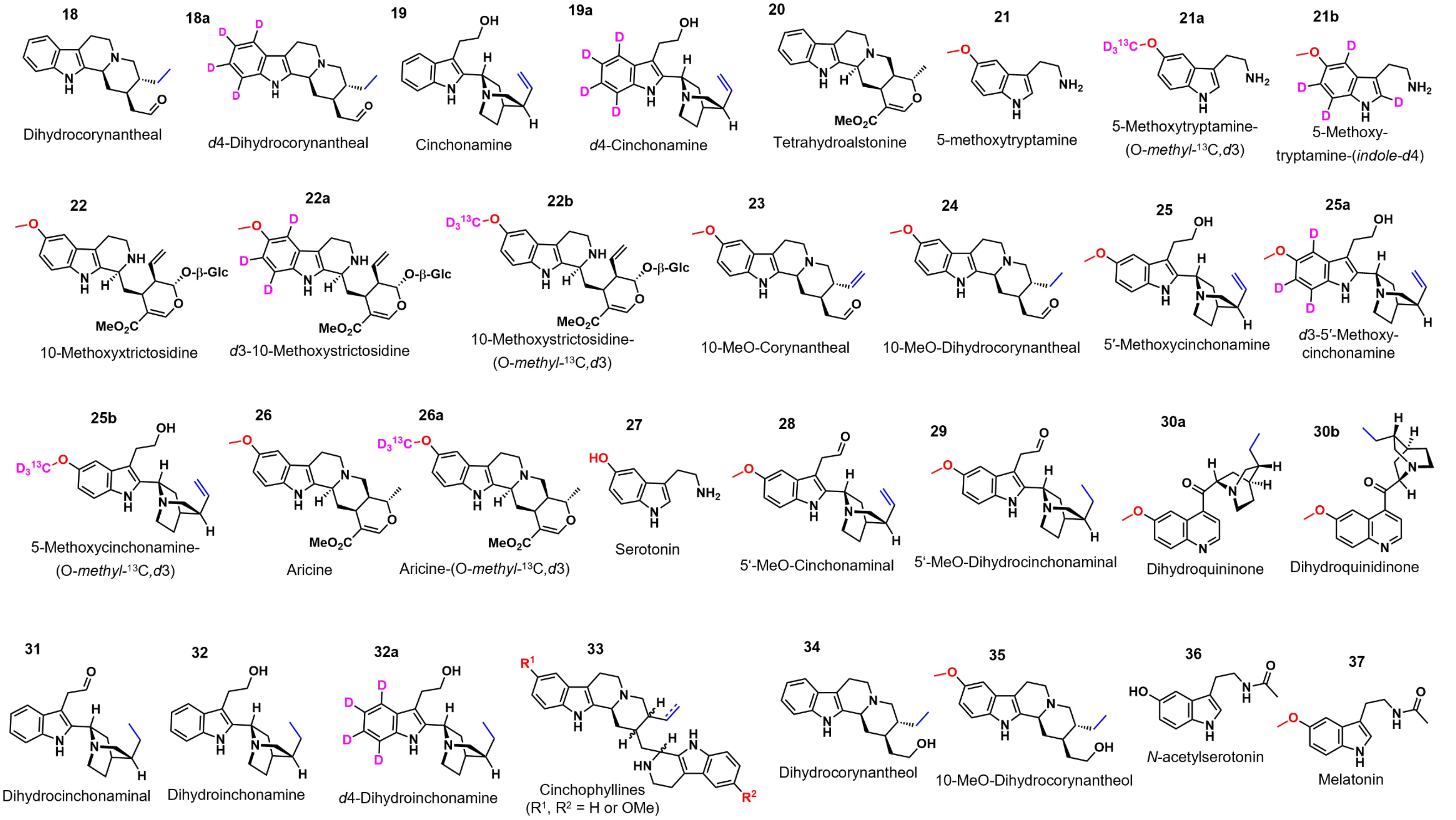


**Figure S1. Compound structures, names, and numbers mentioned in this work.**

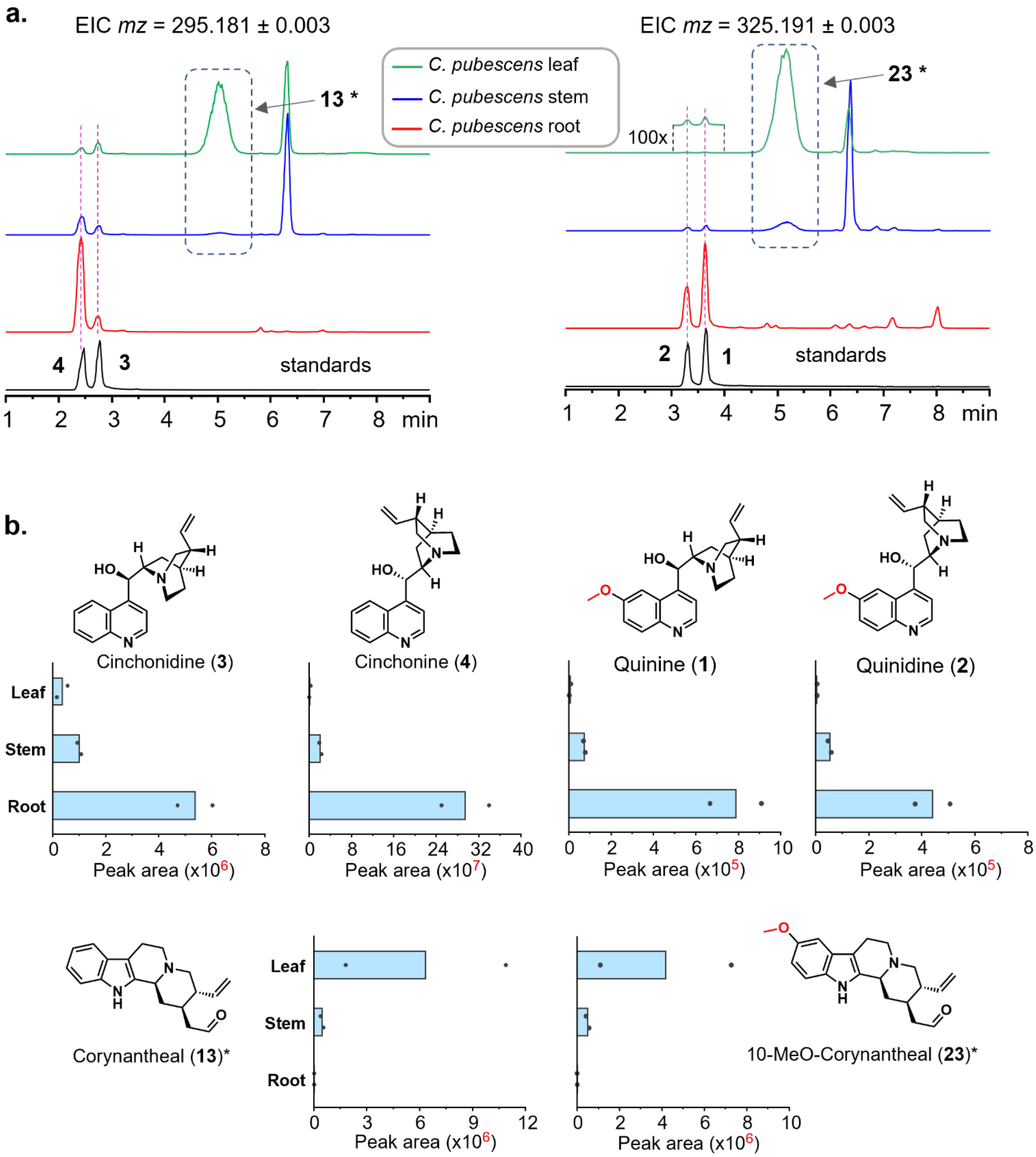


**Figure S2. Alkaloids identified in *Cinchona pubescens* tissues – part I**. **(a)** Extracted ion chromatograms of *m/z* 295.181 (corresponding to cinchonidine **3**, cinchonine **4**, and corynantheal **13**) and *m/z* 325.191 (corresponding to quinine **1**, quinidine **2**, and 10-methoxy-corynantheal **23**) from methanolic extracts of young tissues (leaf, stem, and root) of *C. pubescens* and extracted ion chromatograms of corresponding standards; **(b)** average LC-MS peak area (n = 2 biological replicates) of alkaloids detected in different tissues of *C. pubescens.* The asterisk symbol (*) on **13** and **23** indicates that these compounds were tentatively identified by comparative analyses of MS/MS spectra and chromatographic behavior (peak shape and retention time) with regard to their analog **18** (see Figure S6). The scales of the peak area are colored red to highlight the difference in compound levels.

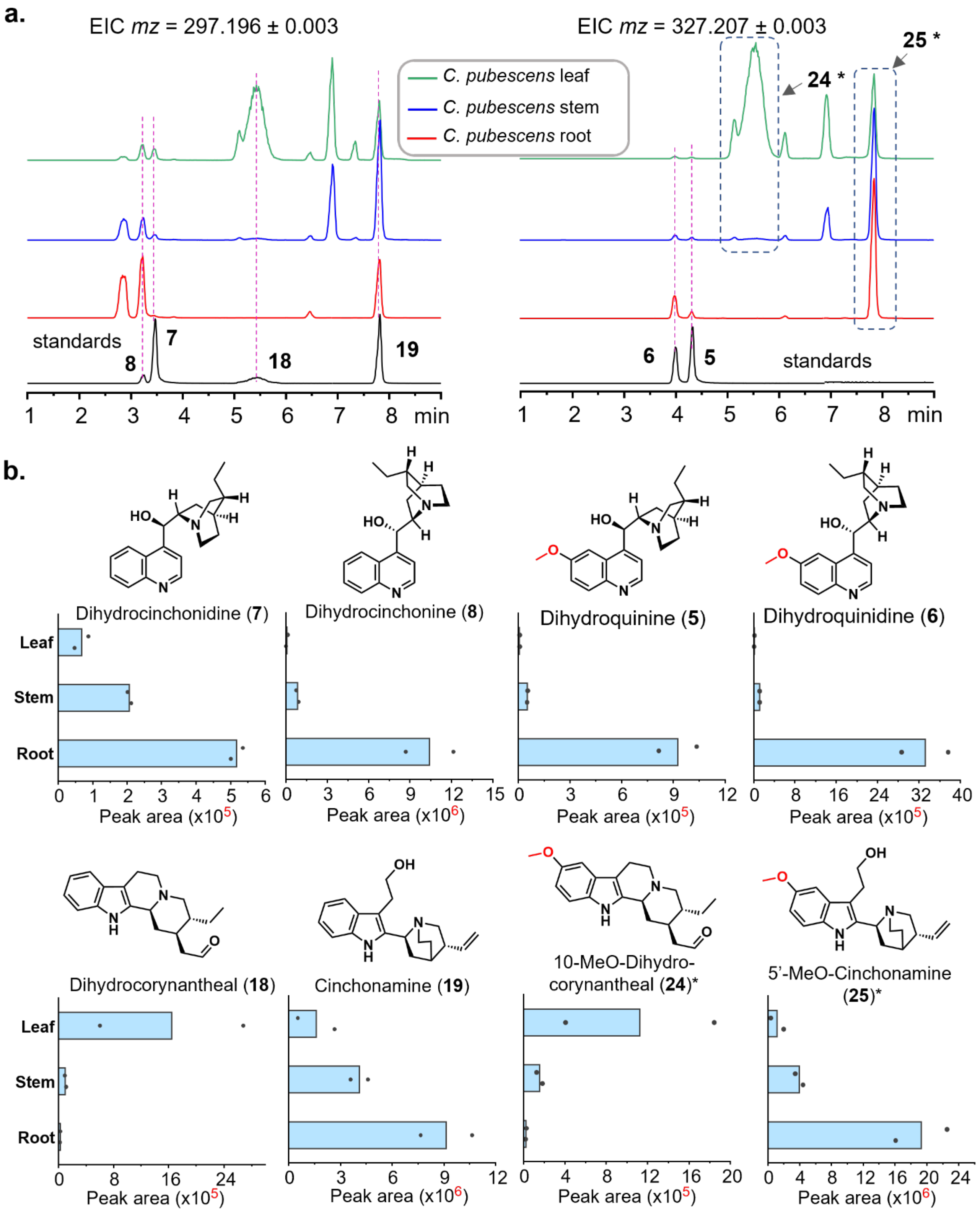


**Figure S3. Alkaloids identified in *C. pubescens* tissues – part II**. **(a)** Extracted ion chromatograms of *m/z* 297.196 (dihydrocinchonidine **7**, dihydrocinchonine **8**, dihydrocorynantheal **18**, and cinchonamine **19**) and *m/z* 327.196 (corresponding to dihydroquinine **5**, dihydroquinidine **6**, 10-methoxy-dihydrocorynantheal **24**, and 5′-methoxycinchonamine **25**); **(b)** average LC-MS peak area (n = 2 biological replicates) of alkaloids detected in different tissues of *C. pubescens.* The asterisk symbol (*) on **24** and **25** denote that these compounds were tentatively identified by comparative analyses of MS/MS spectra and chromatographic behavior (peak shape and retention time) with regard to their respective analog **18** (see Figure S6) and **19** (see Figure S7). The scales of the peak area are colored red to highlight the difference in compound levels.

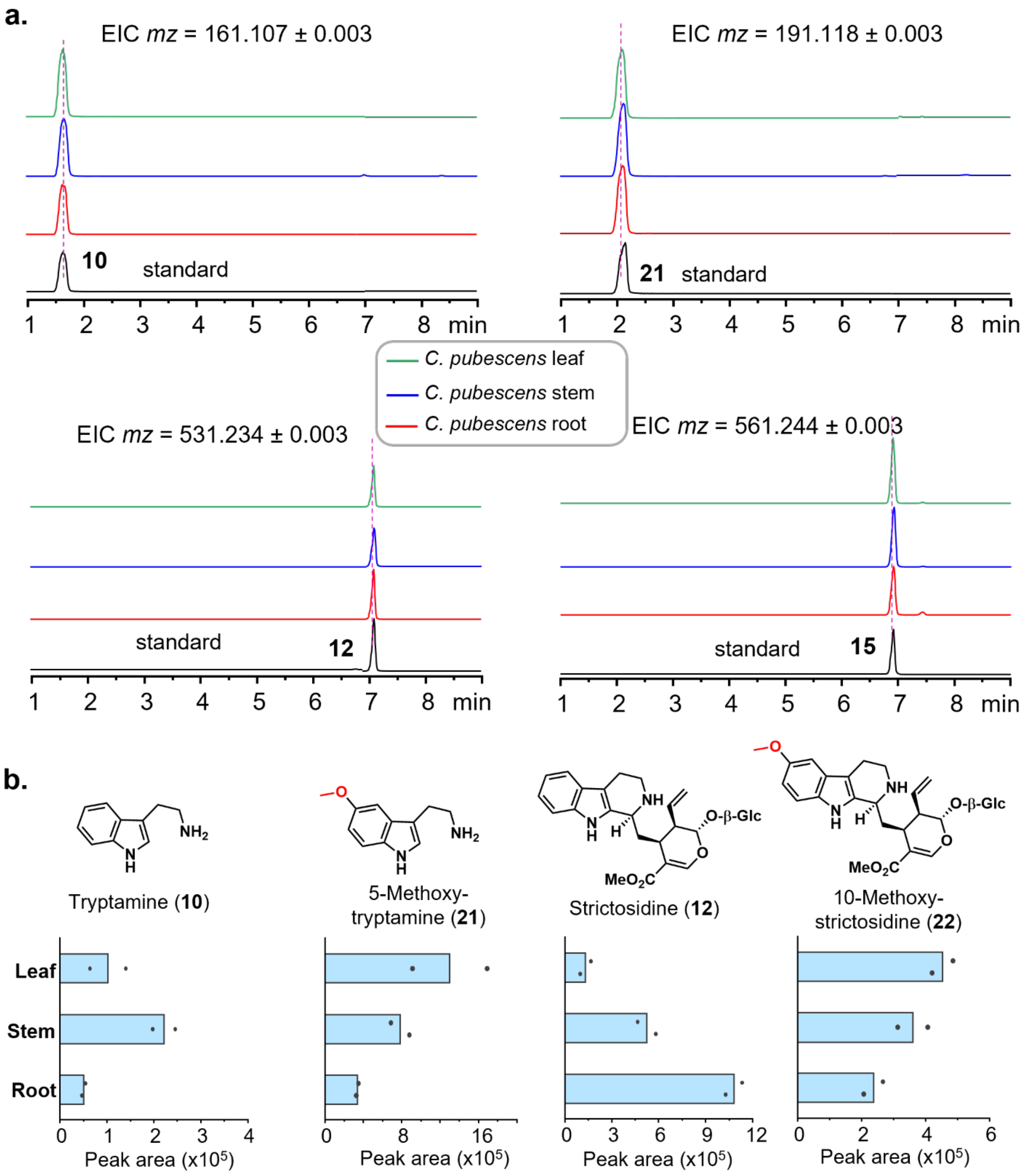


**Figure S4. Alkaloids identified in *Cinchona pubescens* tissues – part III**. **(a)** Extracted ion chromatograms of *m/z* 161.107 (corresponding to tryptamine **10**), *m/z* 191.118 (corresponding to 5-methoxytryptamine **21**), *m/z* 531.234 (corresponding to strictosidine **12**), and *m/z* 561.244 (corresponding to 10-methoxystrictosidine **22**) from methanolic extracts of young tissues (leaf, stem, and root) of *C. pubescens* and extracted ion chromatograms of corresponding standards; **(b)** average LC-MS peak area (n = 2 biological replicates) of alkaloids detected in different tissues of *C. pubescens.*

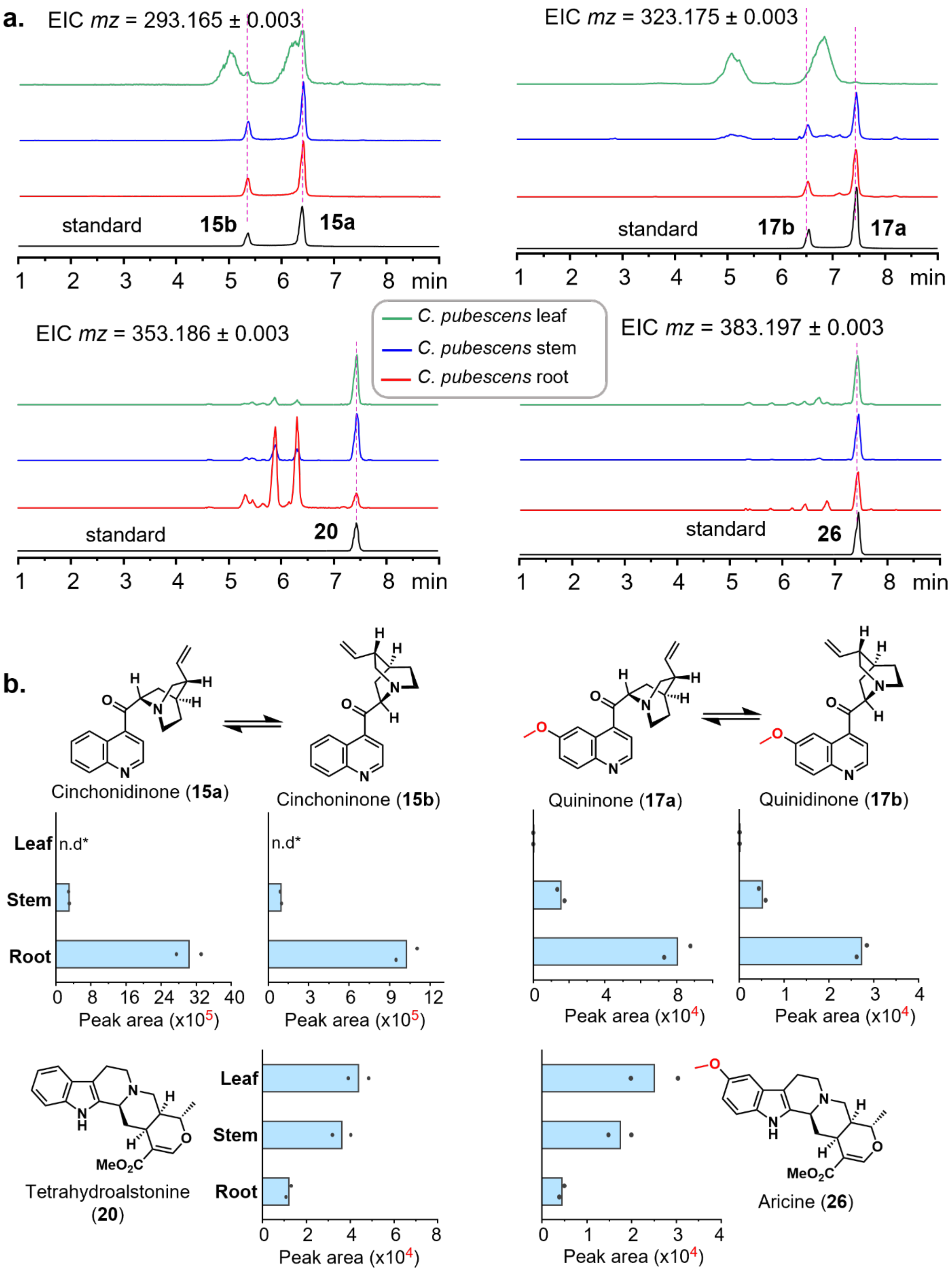


**Figure S5. Alkaloids identified in *Cinchona pubescens* tissues – part IV**. **(a)** Extracted ion chromatograms of *m/z* 293.165 (corresponding to cinchonidinone **15a** and cinchoninone **15b**), *m/z* 323.175 (corresponding to quininone **17a** and quinidinone **17b**), *m/z* 353.186 (corresponding to tetrahydroalstonine **20**), and *m/z* 383.197 (corresponding to aricine or 10-methoxytetrahydroalstonine **26**), from methanolic extracts of young tissues (leaf, stem, and root) of *C. pubescens* and extracted ion chromatograms of corresponding standards; **(b)** average LC-MS peak area (n = 2 biological replicates) of alkaloids detected in different tissues of *C. pubescens.* Note that values for the ketones **15a** and **15b** and of **17a** and **17b**, which are respectively in equilibrium, might be interchanged; here their elution order is depicted by referring to the elution order of their reduced analogs respectively **3** and **4**, and **1** and **2** (see Figure S1). "n.d.*" denotes not determined, due to the overlap of other isobaric compounds.

**a.**

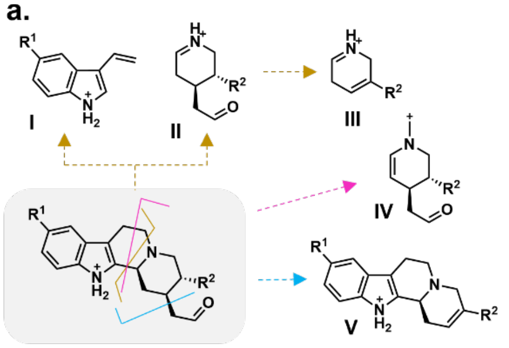


**18** (Dihydrocorynantheal; R¹ : H; R² : $CH_2–CH_3$ )
**24** (10-MeO-Dihydrocorynantheal; R¹ : H; R² : $CH_2–CH_3$ )
**13** (Corynantheal; R¹ : H; R² : $CH_2=CH_2$ )
**23** (10-MeO-Corynantheal; R¹ : H; R² : $CH_2=CH_2$ )

| Fragment | calcd exact mass (chemical formula) | | | |
|---|---|---|---|---|
| | **18**<br>R¹ : H<br>R² : $CH_2–CH_3$ | **24**<br>R¹ : $OCH_3$<br>R² : $CH_2–CH_3$ | **13**<br>R¹ : H<br>R² : $CH_2=CH_2$ | **23**<br>R¹ : $OCH_3$<br>R² : $CH_2=CH_2$ |
| I | 144.0808 ($C_{10}H_{10}N^+$) | 174.0913 ($C_{10}H_{10}N^+$) | 144.0808 ($C_{10}H_{10}N^+$) | 174.0913 ($C_{10}H_{10}N^+$) |
| II | 154.1226 ($C_9H_{16}NO^+$) | 154.1226 ($C_9H_{16}NO^+$) | 152.1070 ($C_9H_{14}NO^+$) | 152.1070 ($C_9H_{14}NO^+$) |
| III | 110.0964 ($C_7H_{12}N^+$) | 110.0964 ($C_7H_{12}N^+$) | 108.0808 ($C_7H_{10}N^+$) | 108.0808 ($C_7H_{10}N^+$) |
| IV | 166.1226 ($C_{10}H_{16}NO^+$) | 166.1226 ($C_{10}H_{16}NO^+$) | 164.1070 ($C_{10}H_{16}NO^+$) | 164.1070 ($C_{10}H_{16}NO^+$) |
| V | 253.1699 ($C_{17}H_{21}N_2^+$) | 283.1805 ($C_{18}H_{23}N_2O^+$) | 251.1543 ($C_{17}H_{19}N_2^+$) | 281.1648 ($C_{18}H_{21}N_2O^+$) |

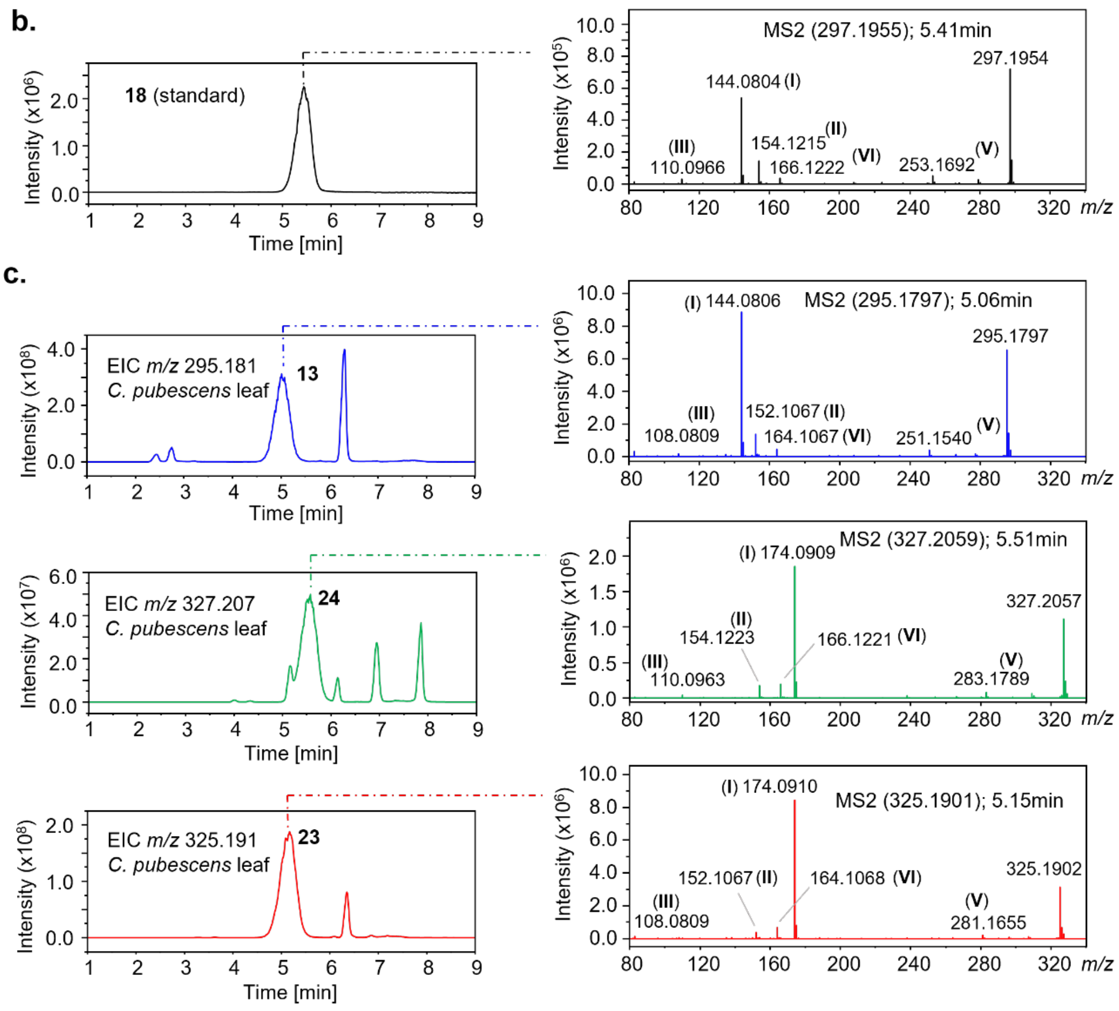


**Figure S6. Identification of corynantheal (13), 10-methoxycorynantheal (23), and 10-methoxydihydrocorynantheal (24) in *C. pubescens*.** **(a)** Putative MS/MS fragments (**I-V**) of **13**, **23**, **24** and of the standard dihydrocorynantheal (**18**) and corresponding calculated exact mass and chemical formula; **(b)** extracted ion chromatogram of the standard **18** and its MS/MS spectrum; **(c)** extracted ion chomatograms corresponding to *mz* of **13**, **24** and **23** from *C. pubescens* leaf extract and the respective MS/MS spectra.

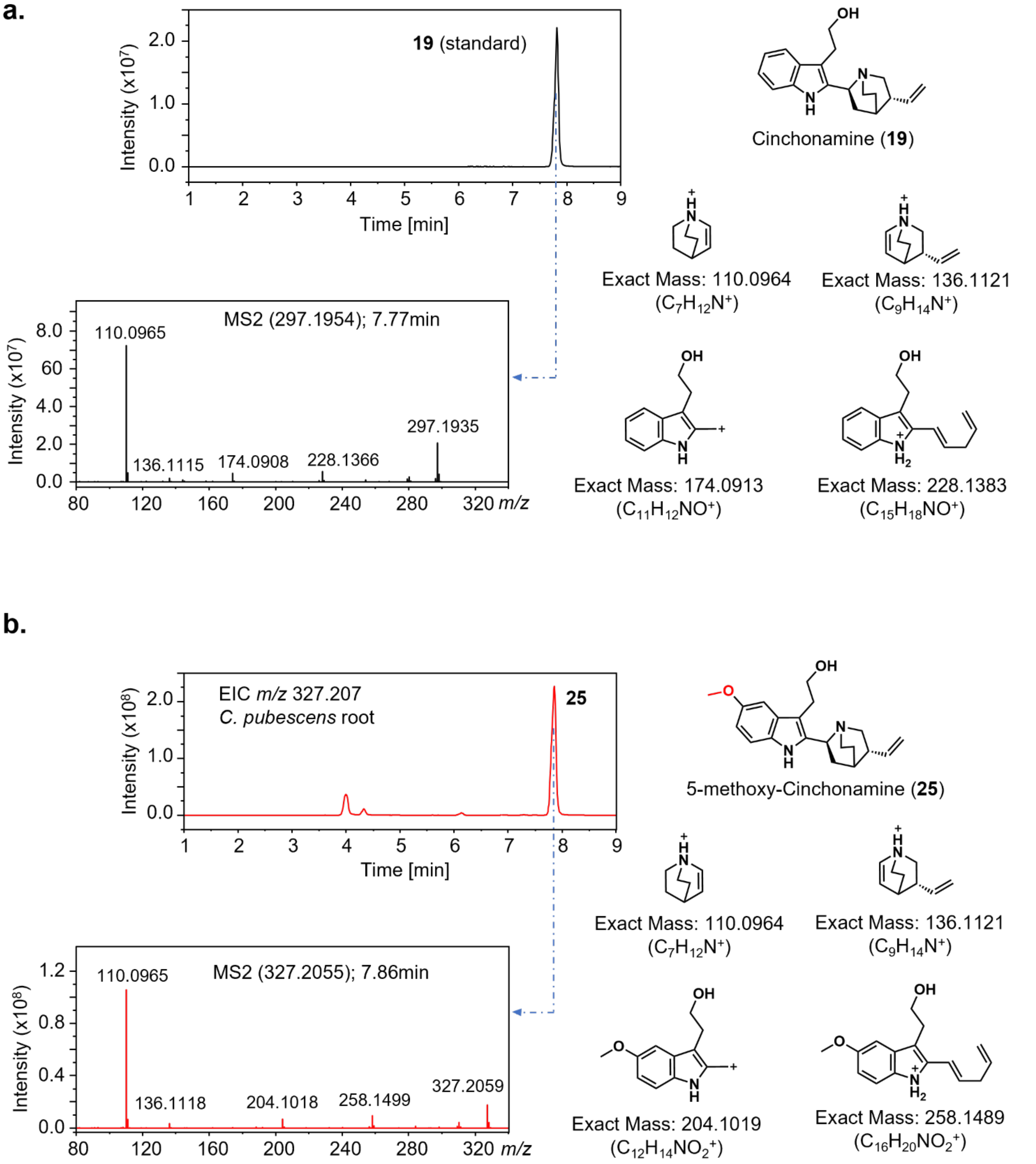


**Figure S7. Identification of 5′-methoxycinchonamine (25) in *Cinchona pubescens*.** **(a)** Extracted ion chromatogram of the standard cinchonamine (**19**), MS/MS spectrum and putative major MS/MS fragments; **(b)** extracted ion chromatogram corresponding to *mz* of methoxylated cinchonamine (**25**), MS/MS spectrum and putative major MS/MS fragments.

a.

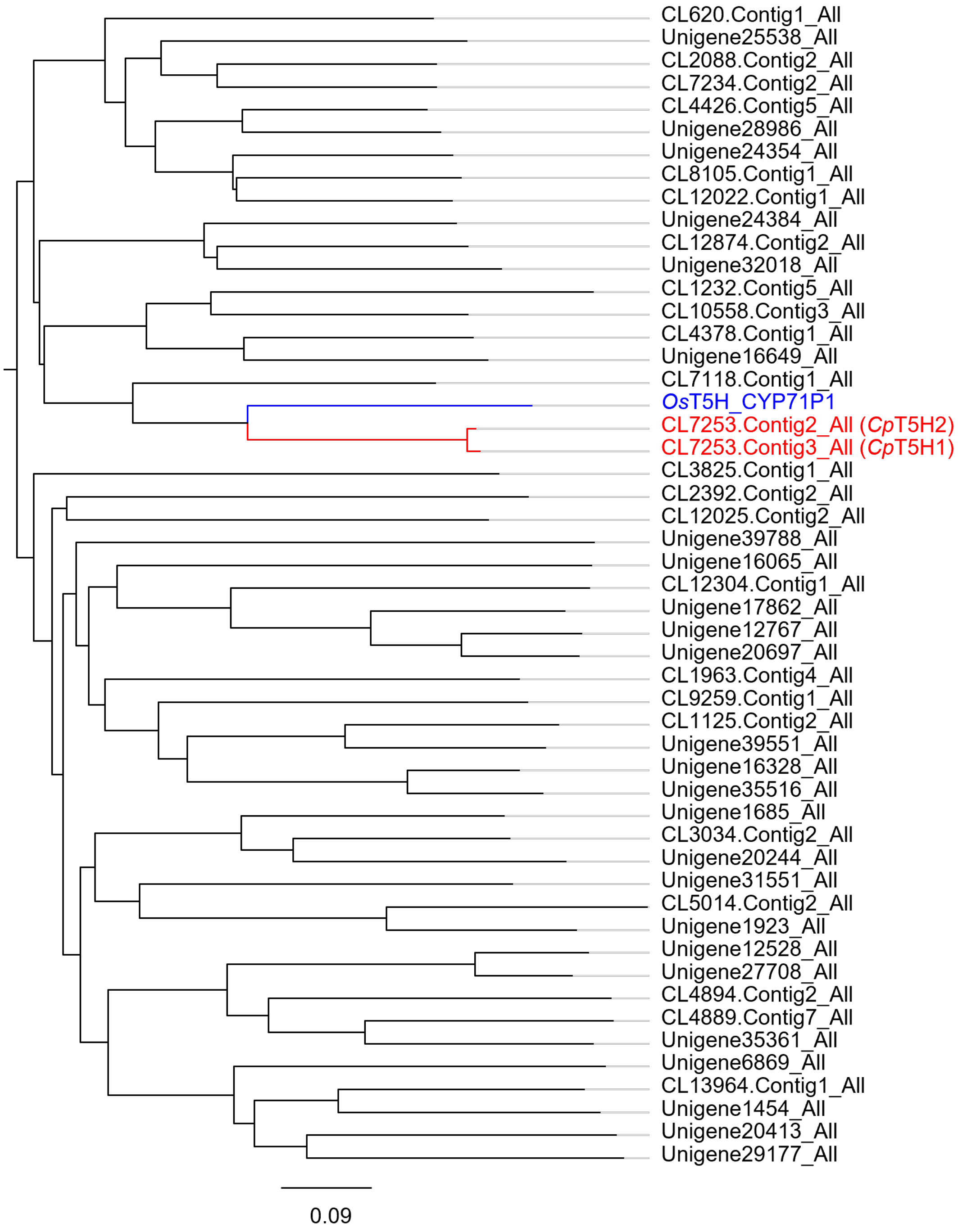


b.

| **Enzymes** | ***Os*T5H** | ***Cp*T5H1** | ***Cp*T5H2** |
|---|---|---|---|
| ***Os*T5H** | | 51.9 % | 52.1 % |
| ***Cp*T5H1** | 51.9 % | | 97.9 % |
| ***Cp*T5H2** | 52.1 % | 97.9 % | |

**Figure S8** continues on the next page

c.

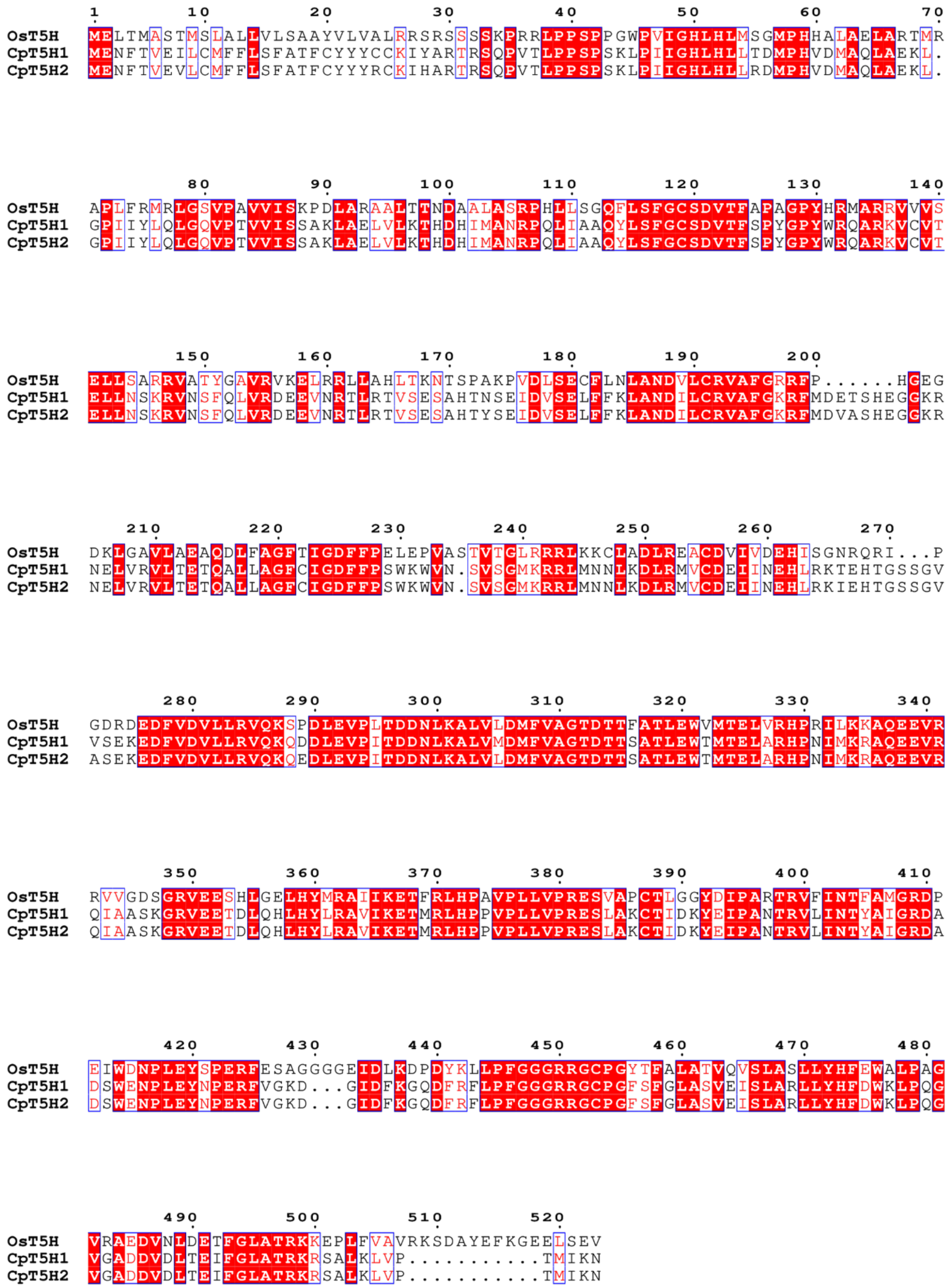


**Figure S8. Identification of *Cinchona pubescens* tryptamine hydroxylases *Cp*T5H1 and *Cp*T5H2. (a)** Phylogenetic relationship of the known[17] *Oryza sativa* tryptamine hydroxylase *Os*T5H with selected putative hydroxylases (CYP450s) from the *Cinchona pubescens* transcriptome; **(b)** amino acid sequence identity matrix of the two hydroxylases functionally identified in this work along with the bait protein, *Os*T5H; **(c)** protein sequence alignment of *Cp*T5H1, *Cp*T5H2 and *Os*T5H. The tree was generated Geneious2024.0.5, Muscle 5.1[18] was used for the calculation of sequence identities and alignment, and ESPript V3 was used for alignment plotting.[19]

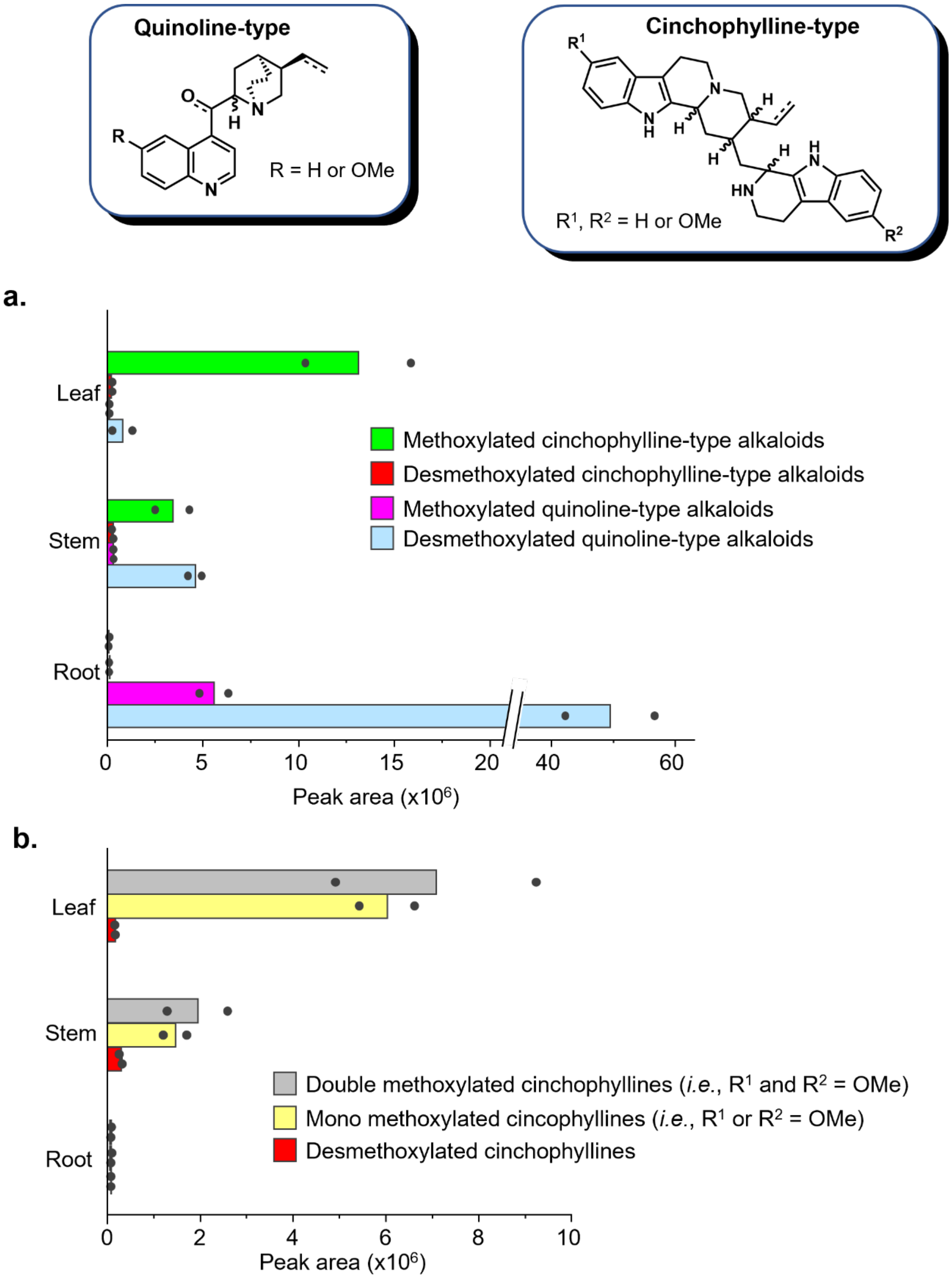


**Figure S9. Distribution of major alkaloids across *Cinchona pubescens* young tissues** (2 biological replicates). **(a)** Combined peak areas of quinoline-type and of cinchophylline-type alkaloids; and **(b)** more detailed distribution among cinchophylline-type metabolites. Quinoline-type metabolites include quinine (**1**), quinidine (**2**), cinchonidine (**3**), cinchonine (**4**) and the dihydro analogs (**5-8**) and ketones (**15a**, **15b**, **17a**, and **17b**) detected based on authentic standards, as shown in Figures S2-S5. Representatives of desmethoxylated and methoxylated cinchophyllines were detected in SIRIUS analyses and, using the MS and MS/MS fragmentations of these hits, closely related analogs were pooled as cinchophyllines. *m/z* for pooled desmethoxylated cinchophyllines were 437.269 ± 0.001 and 439.285 ± 0.001; *m/z* for pooled mono-methoxylated cinchophyllines: 467.279 ± 0.001 and 469.295 ± 0.001; and *m/z* 497.290 ± 0.001 and 499.306 ± 0.001 were used for double-methoxylated cinchophyllines.

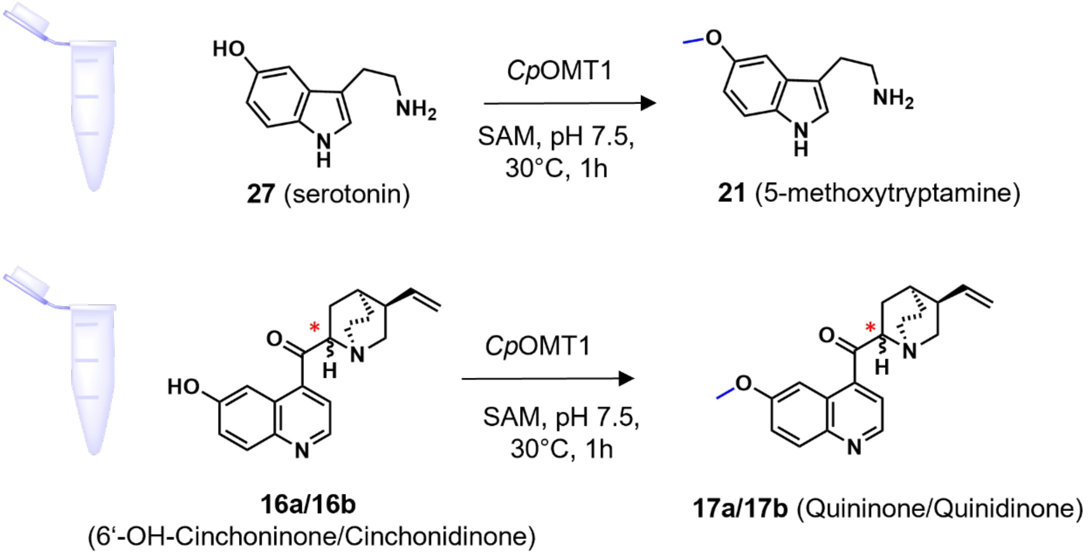


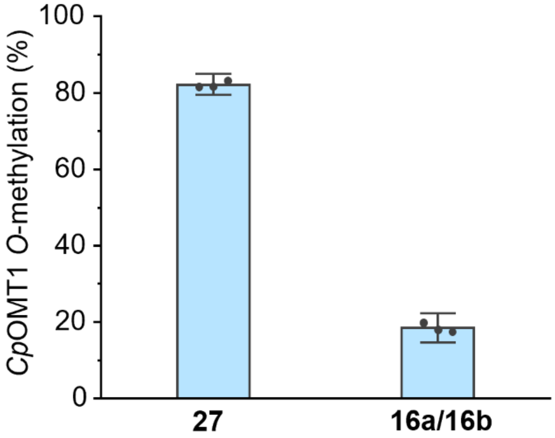


**Figure S10. Comparison of the catalytic activity of *Cinchona pubescens O*-methyltrasferase 1 (*Cp*OMT1) *in vitro*.** *Cp*OMT1 more efficiently converts serotonin (**27**) than the ketones **16a/16b** to the corresponding methylated products. Reaction mix (100 µL) contained 0.5 µM *Cp*OMT1, 50 mM HEPES pH 7.5, 100 µM SAM and 100 µM of substrate (**27** or **16**a/**16**b).

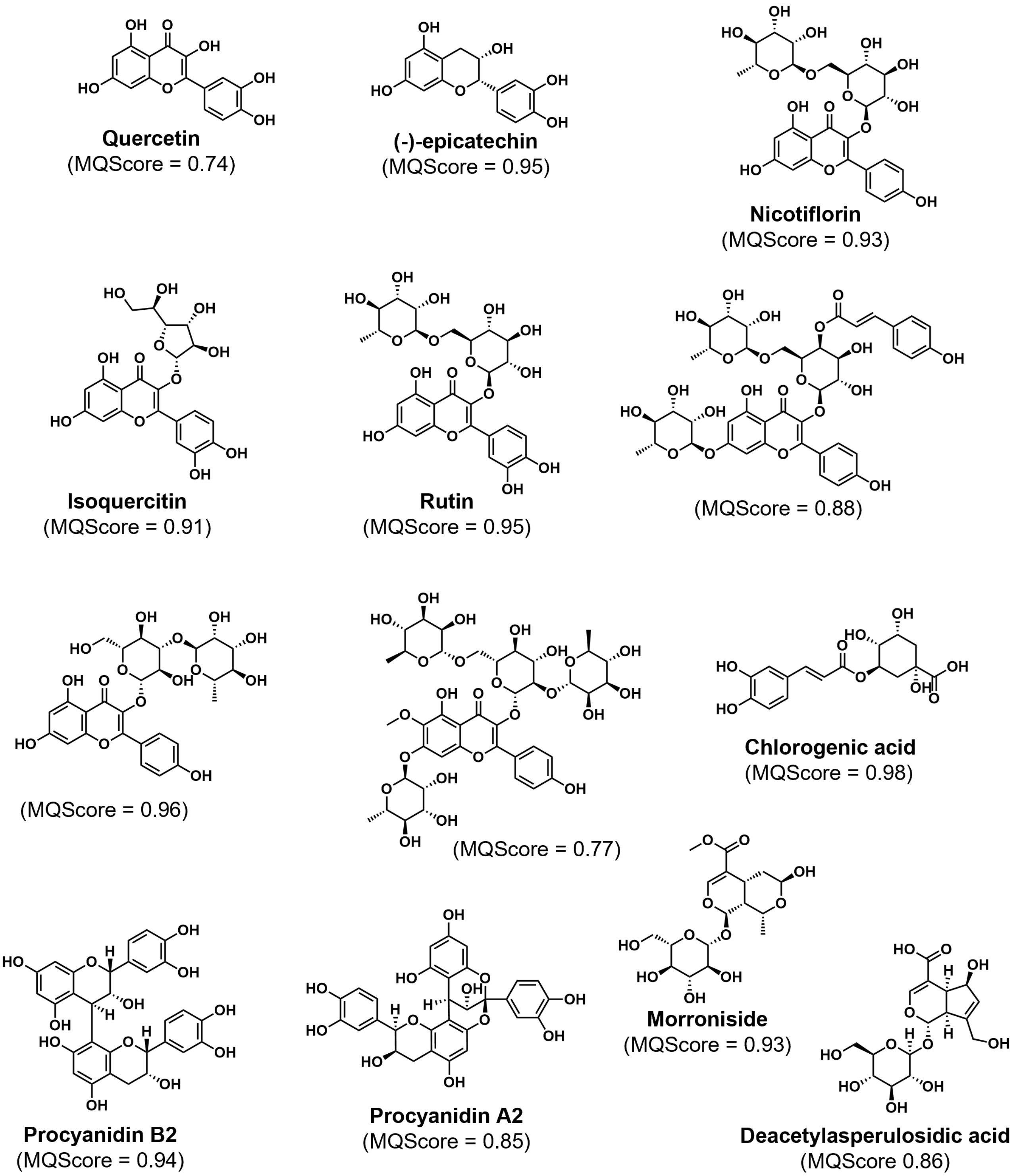


**Figure S11. Selected non-alkaloidal metabolites from *C. pubescens*** (phenolics and iridoids; MQ score > 0.70). These compounds were identified via GNPS feature-based molecular networking and SIRIUS analyses.

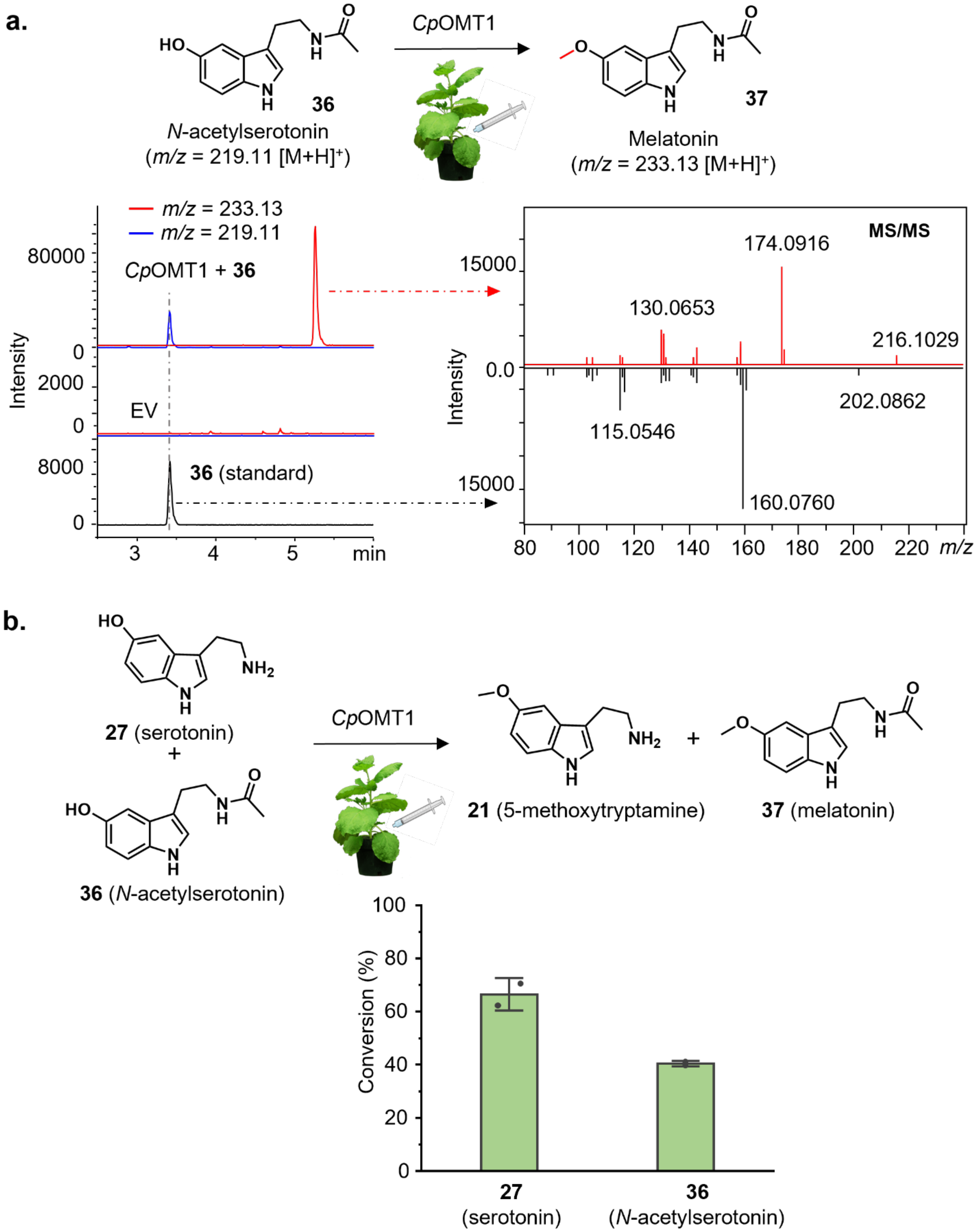


**Figure S12. Substrate specificity of *Cp*OMT1 using *Nicotiana benthamiana* expression system. (a)** Schema illustrating *in planta* assays using *N*-acetylserotonin (**36**) as substrate and *Cp*OMT1 transiently expressed in *N. benthamiana* leaves, and extracted ion chromatograms and MS/MS spectra evidencing that *Cp*OMT1 also *O*-methylates **36**; **(b)** substrate competition assays of *Cp*OMT1, showing that the enzyme seems to preferentially act on serotonin **27** compared to **36**. In addition, neither melatonin **37** nor *N*-acetylserotonin **36** was detected in *C. pubenscens*.

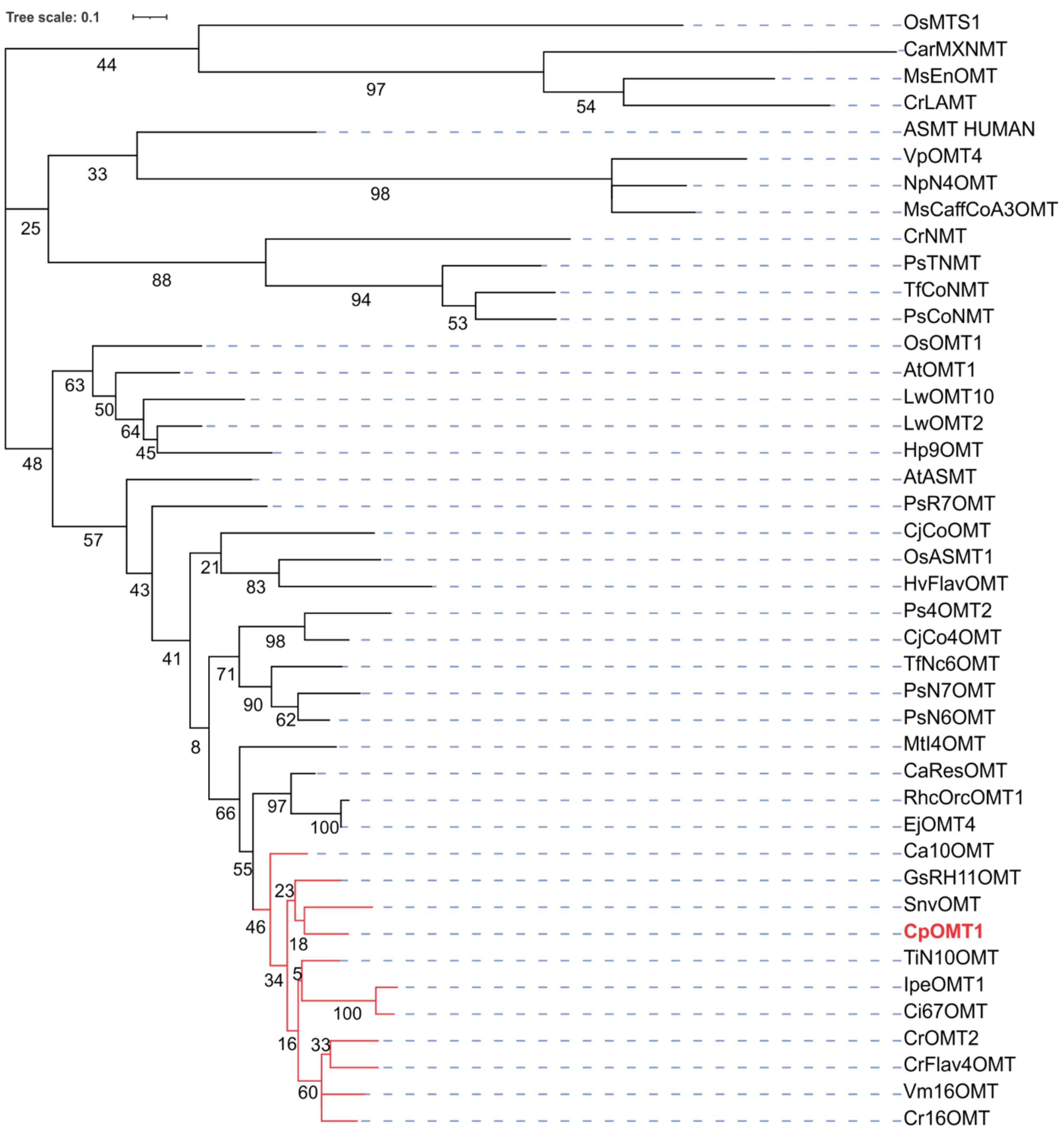


**Figure S13. Phylogenetic tree of *Cp*OMT1 with previously characterized methyltransferases and other hydroxy-indole methyltransferases from monoterpenoid indole alkaloid (MIA)-producing plants.** The clade of aromatic MIA *O*-methyltransferases is highlighted in red. The enzymes full names, plant species, and Genbank/Uniprot accession numbers are included in the Table S3.

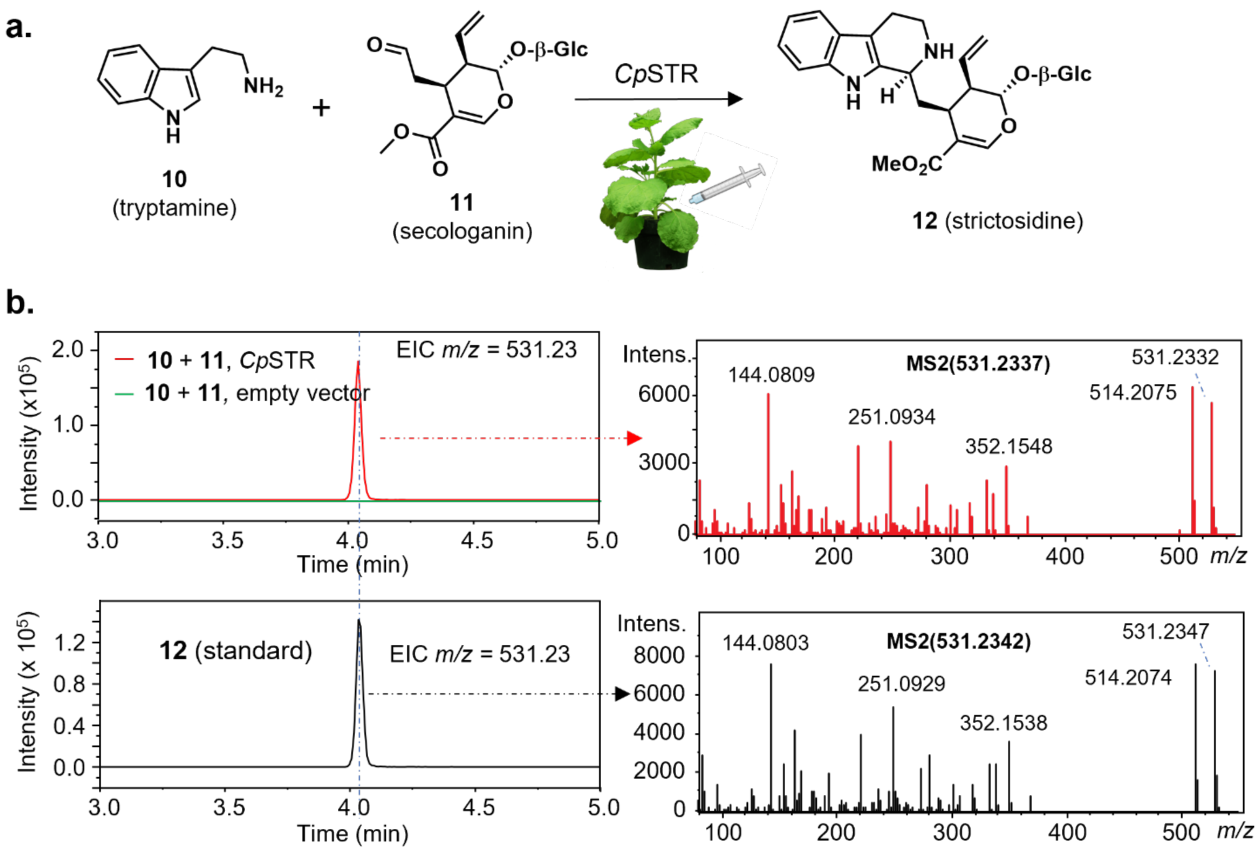


**Figure S14. Biochemical characterization of *Cinchona pubescens* strictosidine synthase (*Cp*STR) using expression in *Nicotiana benthamiana*. (a)** Scheme illustrating *in planta* assays using secologanin (**11**) and tryptamine (**10**) as substrates and *Cp*STR transiently expressed in *N. benthamiana* leaves; **(b)** extracted ion chromatograms consistent with *m/z* of strictosidine (531.23) and corresponding MS/MS spectra, evidencing formation of strictosidine **12**.

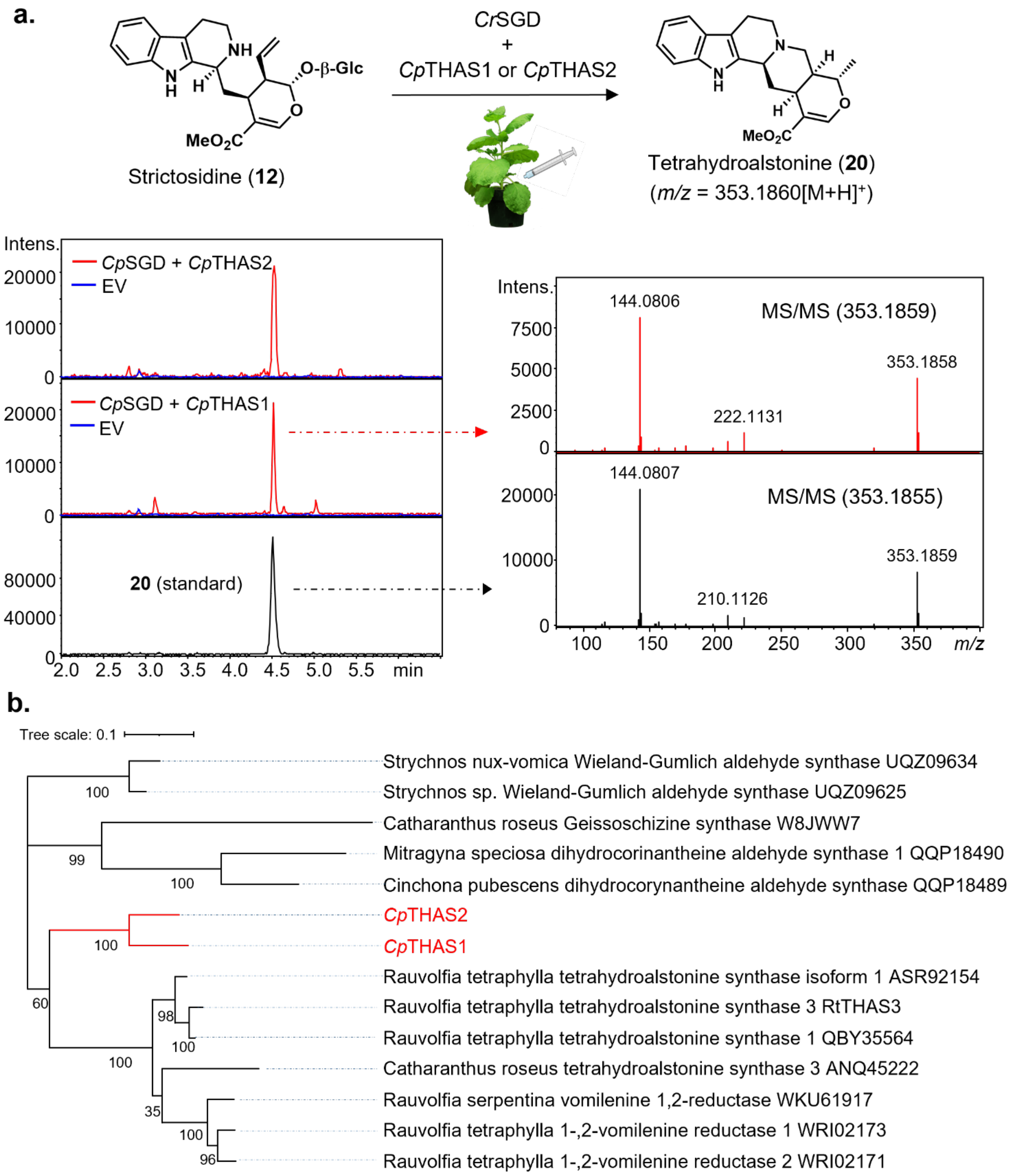


**Figure S15. Identification of Biochemical characterization of *Cinchona pubescens* tetrahydroalstonine synthases (*Cp*THAS1-2). (a)** Biochemical characterization of *Cp*THAS1-2 via expression in *Nicotiana benthamiana.* The schema illustrates the *in planta* assays using strictosidine (**12**) as substrate and *Cr*SGD transiently co-expressed in *N. benthamiana* leaves with either *Cp*THAS-1 or *Cp*THAS-2. Extracted ion chromatograms and MS/MS spectra evidencing the formation of **20** are also shown; **(b)** phylogentic relationship of *Cp*THASs with related characterized MIAs reductases.

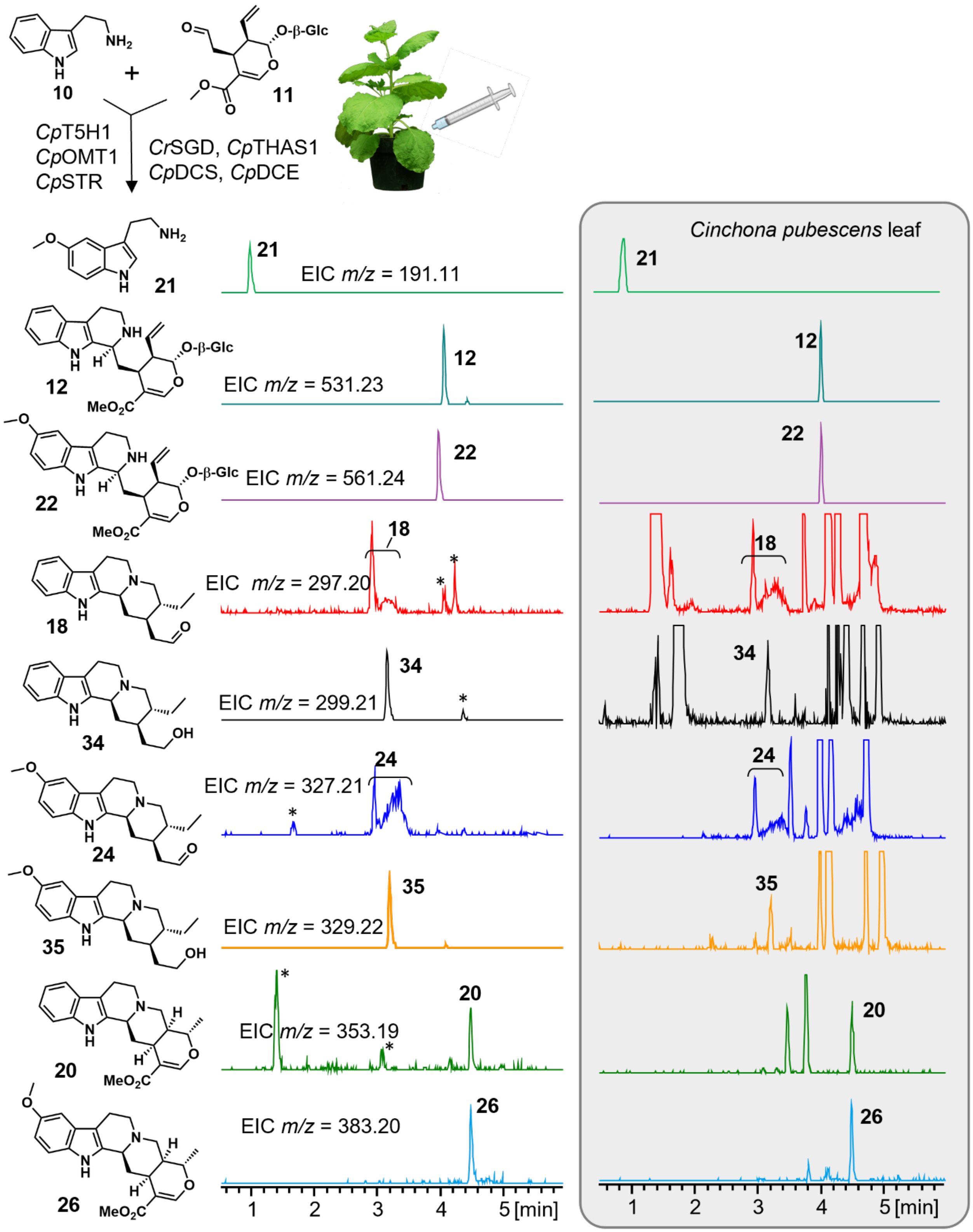


**Figure S16**. **Heterologous production of early pathway Cinchona alkaloids in *N. benthamiana***. Extracted ion chromatograms of *m/z* corresponding to *Cinchona* metabolites from a methanolic extract of an *N. benthamiana* leaf expressing the shown *Cinchona* biosynthetic genes. The occurrence of the corresponding targeted metabolites in a *Cinchona* leaf extract is shown in the box. (*) indicates tobacco endogenous metabolites.

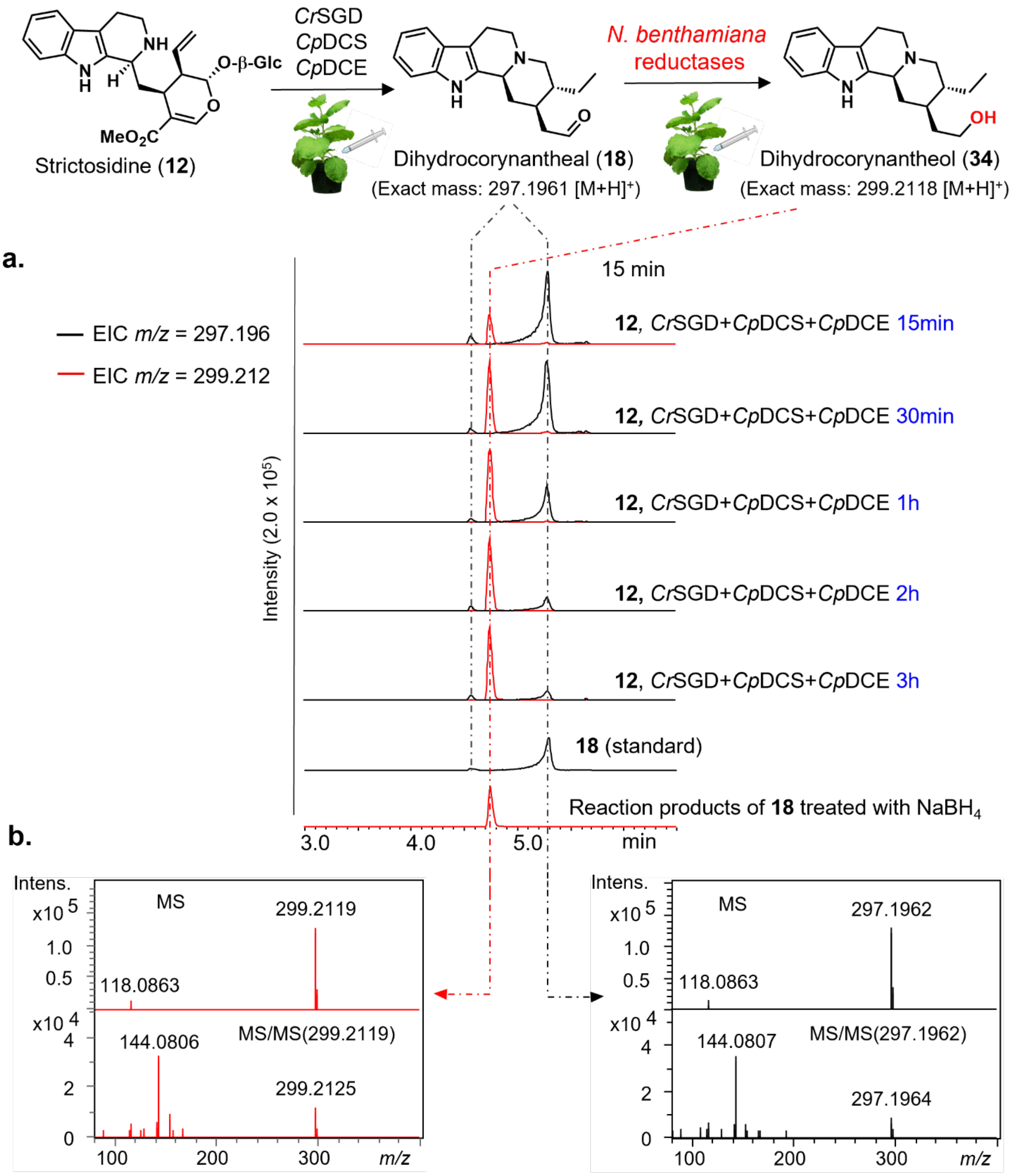


**Figure S17**. **Reduction of dihydrocorynantheal (18) into dihydrocorynantheol (34) by *N. benthamiana* endogenous enzyme activity**. **(a)** Extracted ion chromatograms showing *m/z* corresponding to **18** and the corresponding alcohol **34** from an *N. benthamiana* leaf at different times post-infiltration of strictosidine **12**; and **(b)** MS/MS spectra of the standard and *in-situ* produced **18** and of a standard and *in-planta* generated **34**. **18** is observed as two peaks in the chromatogram due to the chemical equilibrium of the aldehyde and enol forms.

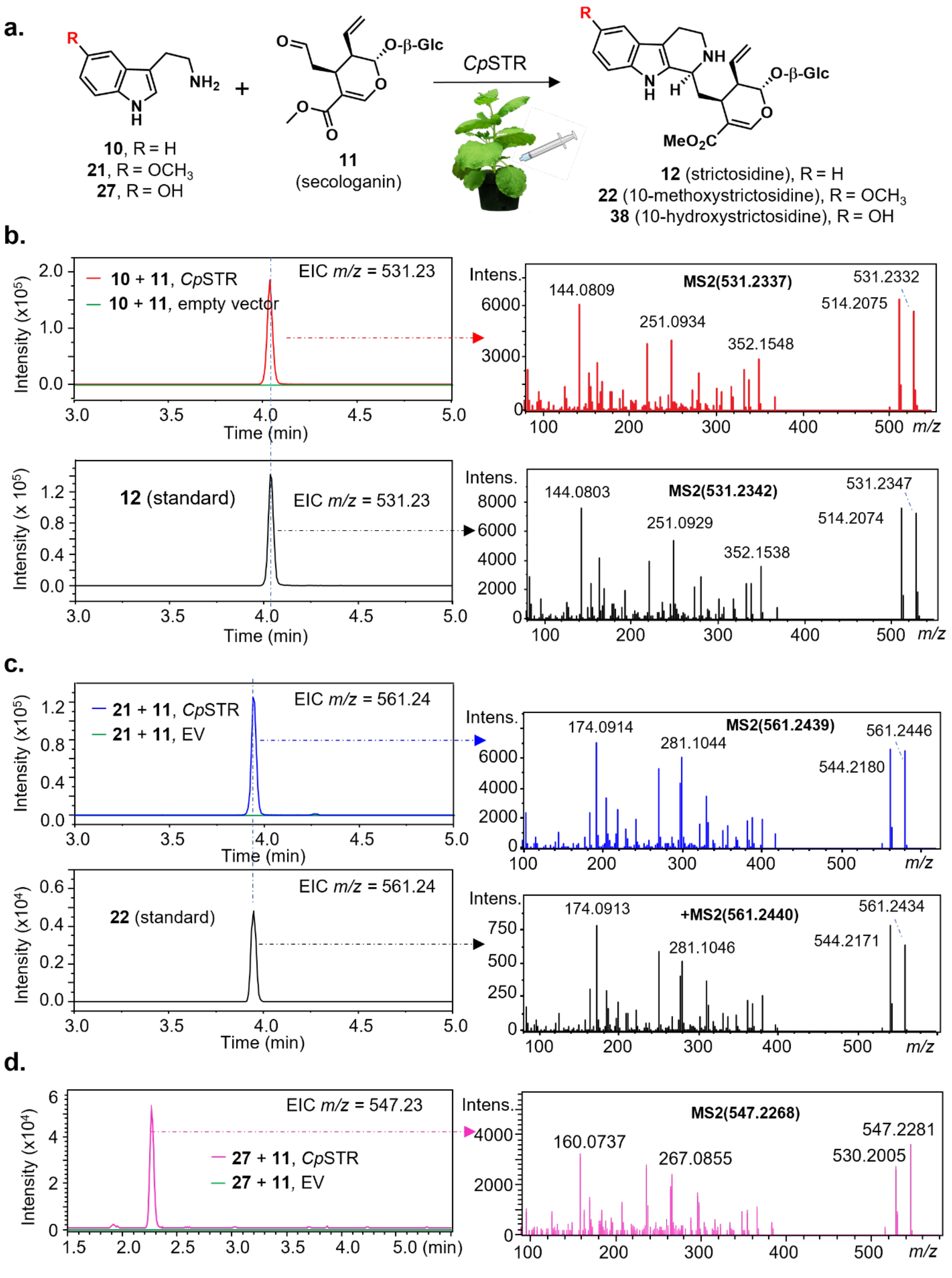


**Figure S18. Substrate scope of *Cinchona pubescens* strictosidine synthase (*Cp*STR) using expression in *Nicotiana benthamiana*. (a)** Scheme illustrating *in planta* assays using secologanin (**11**) with either tryptamine

(**10**), 5-methoxytryptamine (**21**), or serotonin (**27**) as substrates and *Cp*STR transiently expressed in *N. benthamiana* leaves; **(b)** extracted ion chromatograms consistent with *m/z* of strictosidine (531.23) and corresponding MS/MS spectra, evidencing formation of strictosidine **12**; **(c)** extracted ion chromatograms consistent with *m/z* of 10-methoxystrictosidine (561.24) and corresponding MS/MS spectra, establishing biosynthesis of **22**; **(d) e**xtracted ion chromatograms consistent with *m/z* of 10-methoxystrictosidine (547.23) and corresponding MS/MS spectra, showing the formation of 10-hydroxystrictosidine **38**.

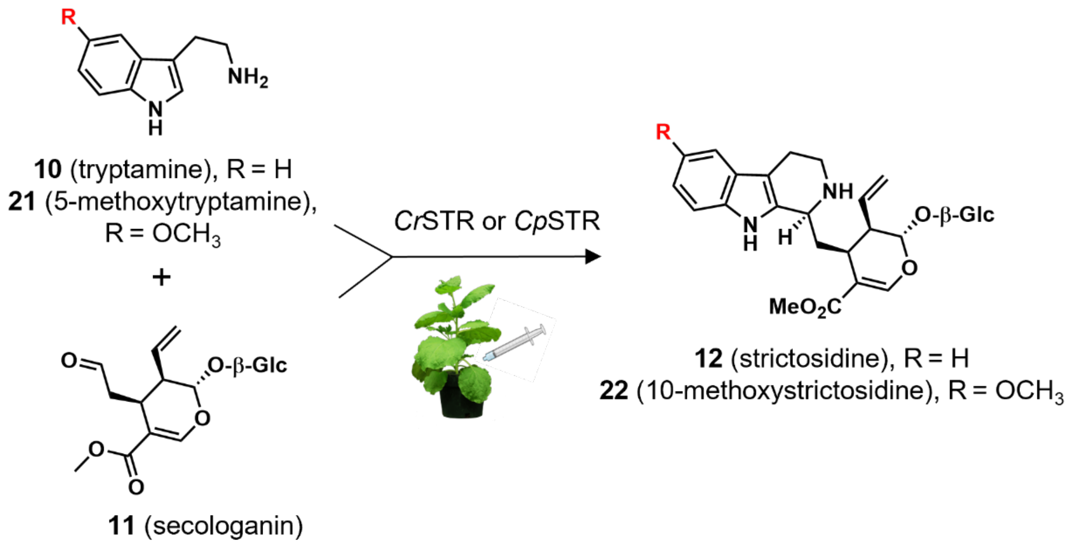


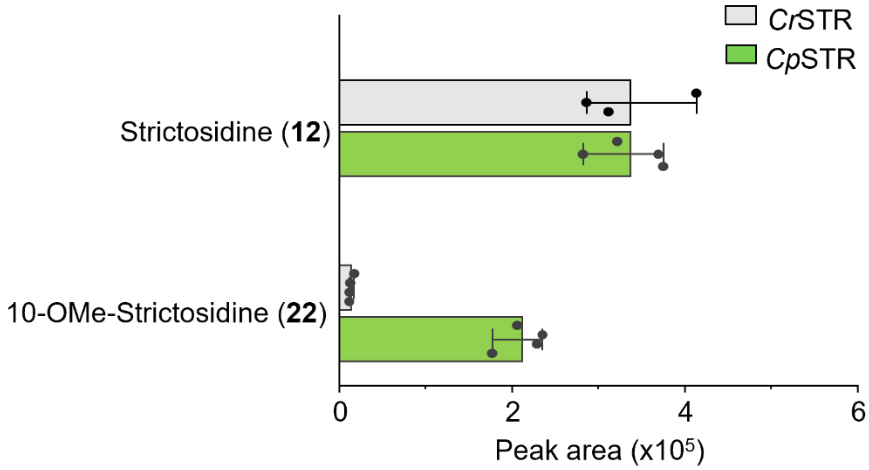


**Figure S19. Comparison of the substrate scope of strictosidine synthases from *Cinchona pubescens* (*Cp*STR) and *Catharanthus roseus* (*Cr*STR) in *Nicotiana benthamiana*.** *Cp*STR accepts both tryptamine (**5**) and methoxylated analog (**13**) while *Cr*STR shows specificity towards **5**. These *in planta* results are consistent with early *in vitro* studies on STR purified from *Cinchona robusta* suspension cultures.[20]

**a.**

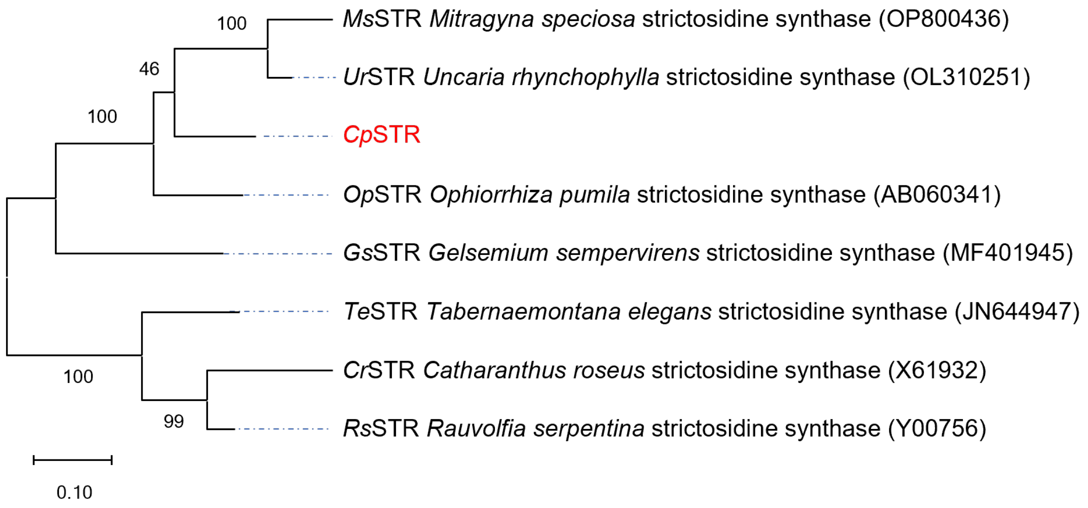


**b.**

| Enzymes | Amino acids identity (%) | | | | | | |
|---|---|---|---|---|---|---|---|
| | *Cr*STR | *Gs*STR | *Ms*STR | *Op*STR | *Rs*STR | *Te*STR | *Ur*STR |
| *Cp*STR | 51.5 | 63 | 71.7 | 76.2 | 54.7 | 55.9 | 74.9 |

**Figure S20. Evolutionary analysis of *Cinchona pubescens* strictosidine synthase (*Cp*STR) identified in this work. (a)** Phylogenetic tree with previously characterized strictosidine synthases; **(b)** amino-acid similarity of *Cp*STR with these related enzymes orthologs.

.

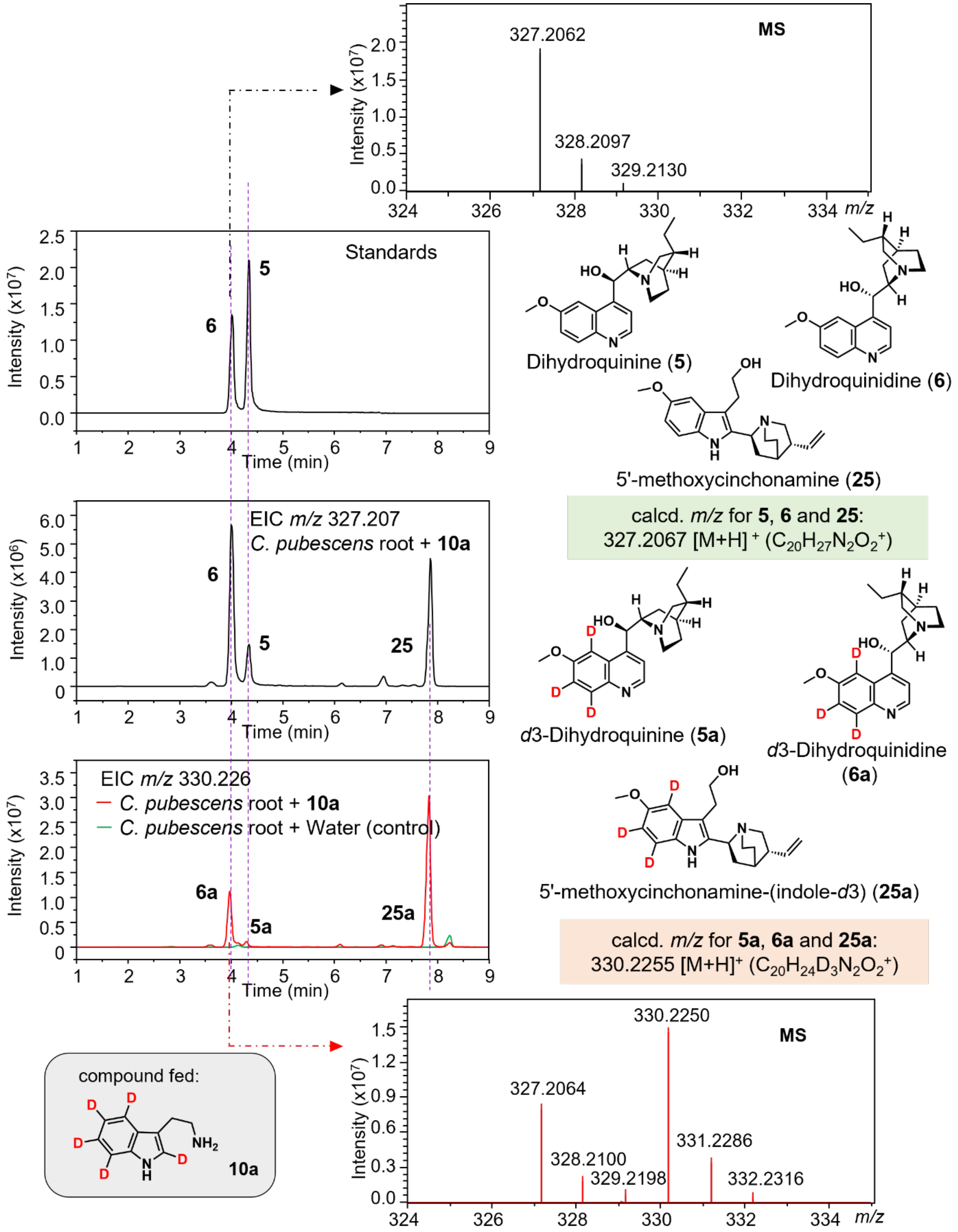


**Figure S21. Incorporation of tryptamine-(*indole-d*5) (10a) into dihydroquinine (5), dihydroquinidine (6), and 5′-methoxycinchonamine (25).** Extracted ion chromatograms and isotope patterns, evidencing the formation of **5a**, **6a** and **25a** in roots of *Cinchona pubescens* fed with **10a**. For MS/MS spectra, see Figure S22 for **6a** (and, thus, **5a**) and Figure S23 for **25a**.

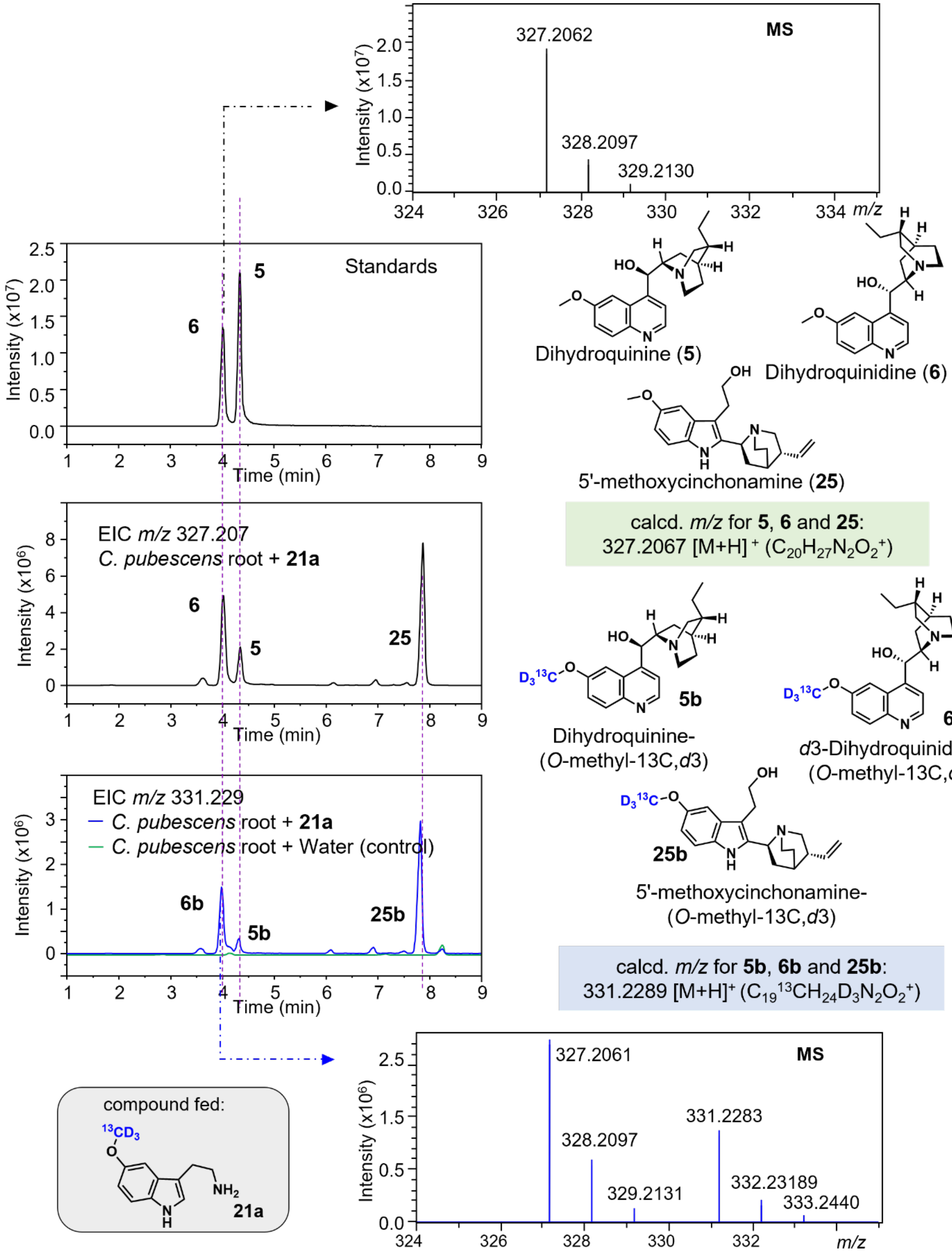


**Figure S22. Incorporation of 5-methoxytryptamine-(O-*methyl*-$^{13}$C, *d*3) (21a) into dihydroquinine (5), dihydroquinidine (6), and 5′-methoxycinchonamine (25).** Extracted ion chromatograms and isotope patterns, evidencing the formation of **5b**, **6b** and **25a** in roots of *Cinchona pubescens* fed with **21a**. For MS/MS spectra, see Figure S22 for **5b** and **6a** and Figure S23 for **25a**.

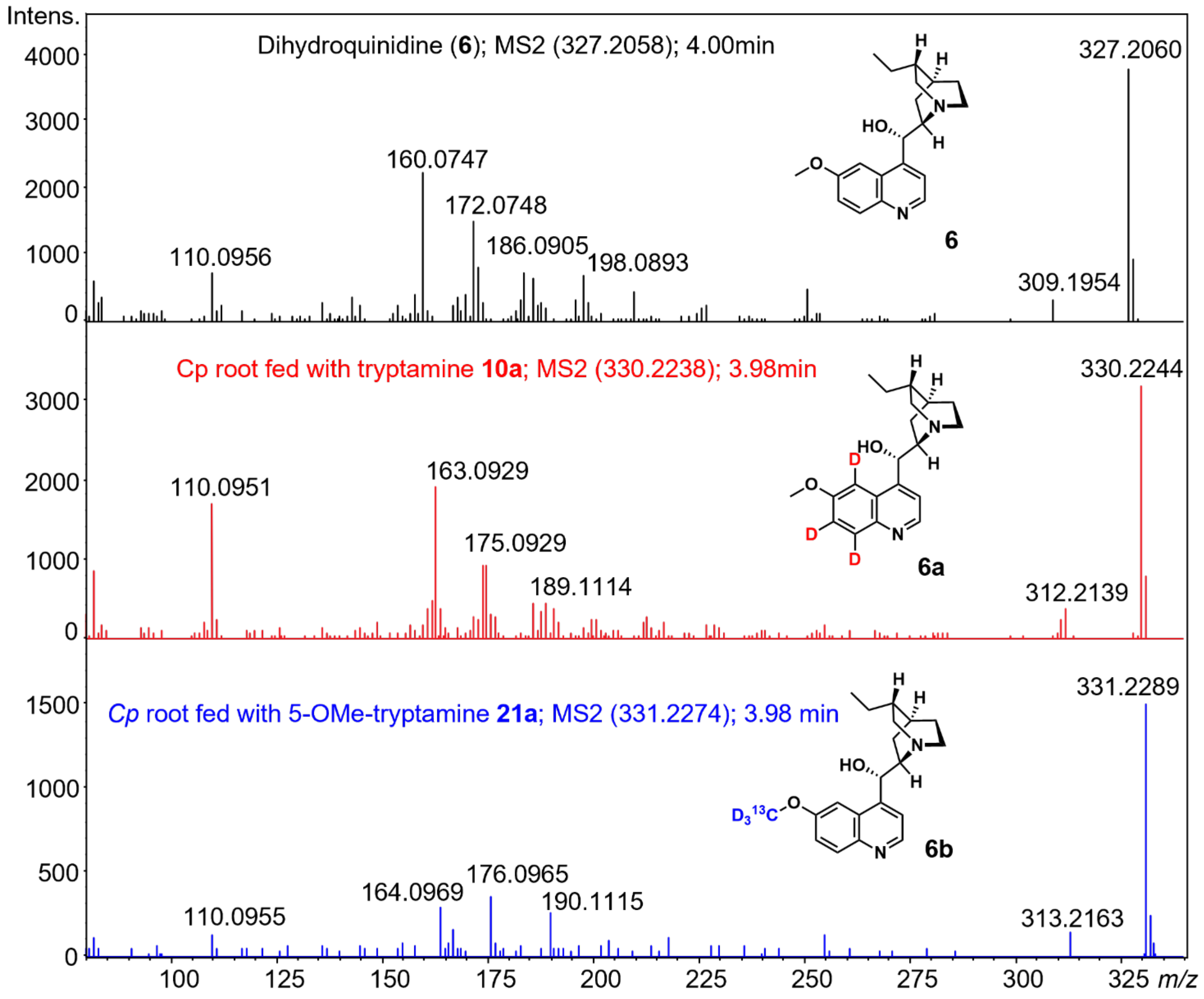


**Figure S23. MS/MS spectra of the standard dihydroquinidine (6) and of isotope-labeled analogs 6a and 6b.** The latter were detected in roots of *Cinchona pubescens* after feeding plantlet roots with tryptamine-(indole-*d*5) **10a** and 5-methoxytryptamine-(O-*methyl*-$^{13}$C, *d*3) **21a**, respectively.

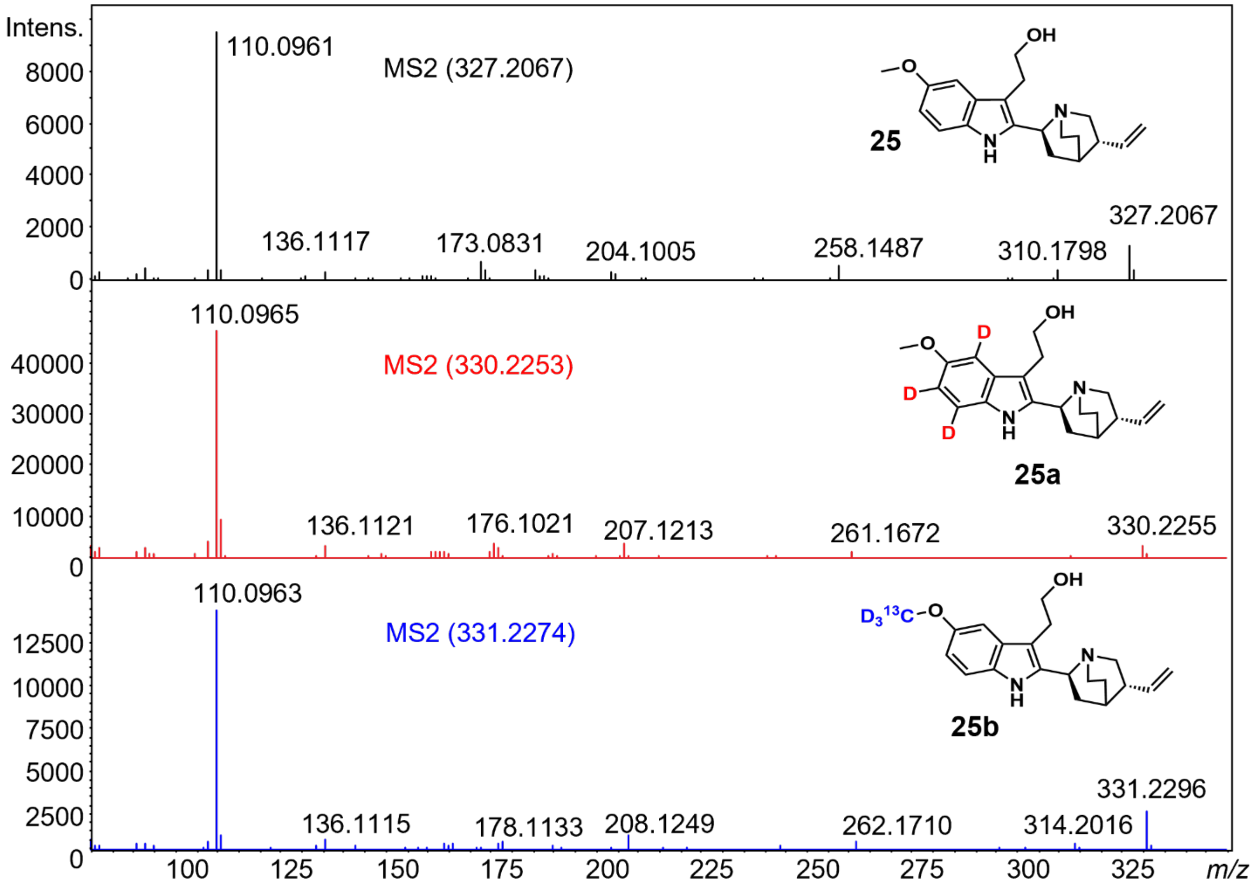


**Figure S24. MS/MS spectra of 5′-methoxycinchonamine and of isotope-labeled analogs 25a and 25b.** The latter were detected in *Cinchona pubescens* roots after supplementation of tryptamine-(indole-*d*5) **10a** and 5-methoxytryptamine-(O-*methyl*-$^{13}$C, *d*3) **21a**, respectively.

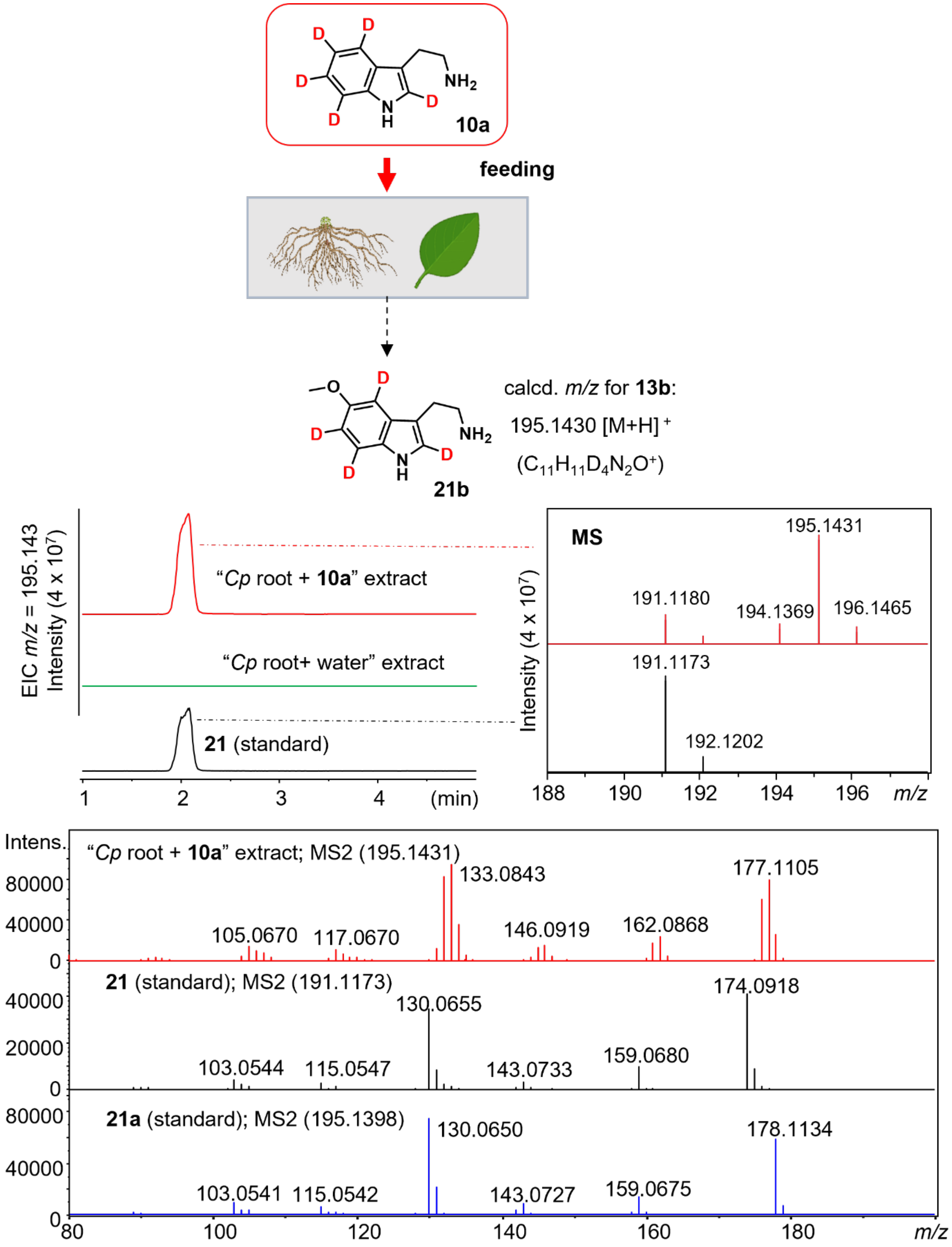


**Figure S25. Conversion of tryptamine-(indole-*d*5) (10a) into 5-methoxytryptamine-(*indole-d*4) (21b).** Extracted ion chromatograms, isotope patterns, and MS/MS spectra evidencing the formation of **21b** in roots of *Cinchona pubescens* fed with **10a**. The MS/MS spectrum of the standard **21a**, which is almost isobaric to **21b** ($\Delta m/z$ = 0.003), is here also shown for comparison.

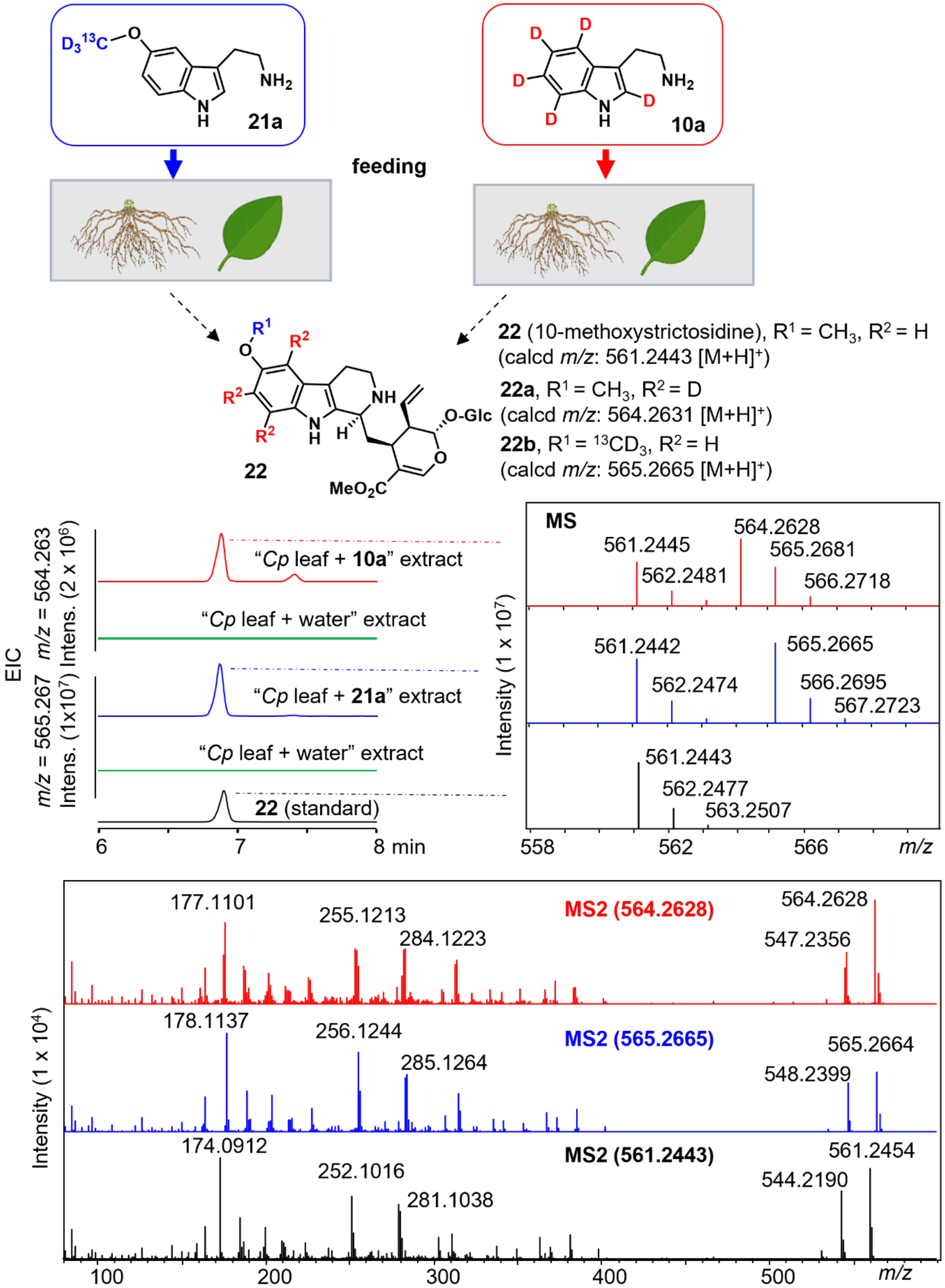


**Figure S26. Incorporation of tryptamine-(indole-*d*5) (10a) and 5-methoxytryptamine-(O-*methyl*-$^{13}$C, *d*3) (21a) into 10-methoxystrictosidine (22).** Extracted ion chromatograms, isotope patterns, and MS/MS spectra evidencing the formation of **22a** and **22b** in roots of *Cinchona pubescens* tissues fed with **10a** and **21a**, respectively.

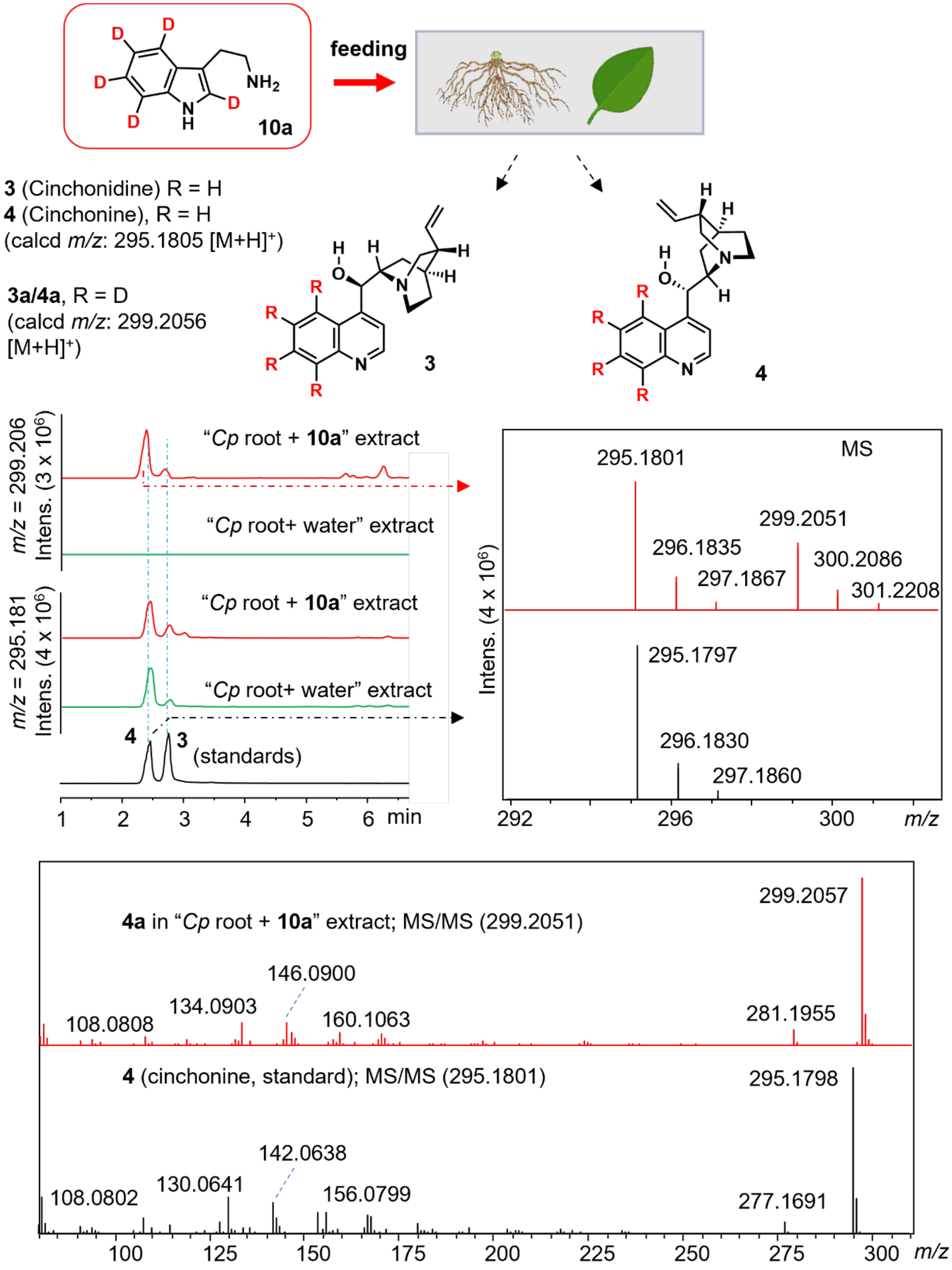


**Figure S27. Incorporation of tryptamine-(indole-*d*5) (10a) into cinchonidine (3) and cinchonine (4).** Extracted ion chromatograms, isotope patterns, and MS/MS spectra evidencing the formation of **3a** and **4a** in roots of *Cinchona pubescens* tissues fed with **10a**.

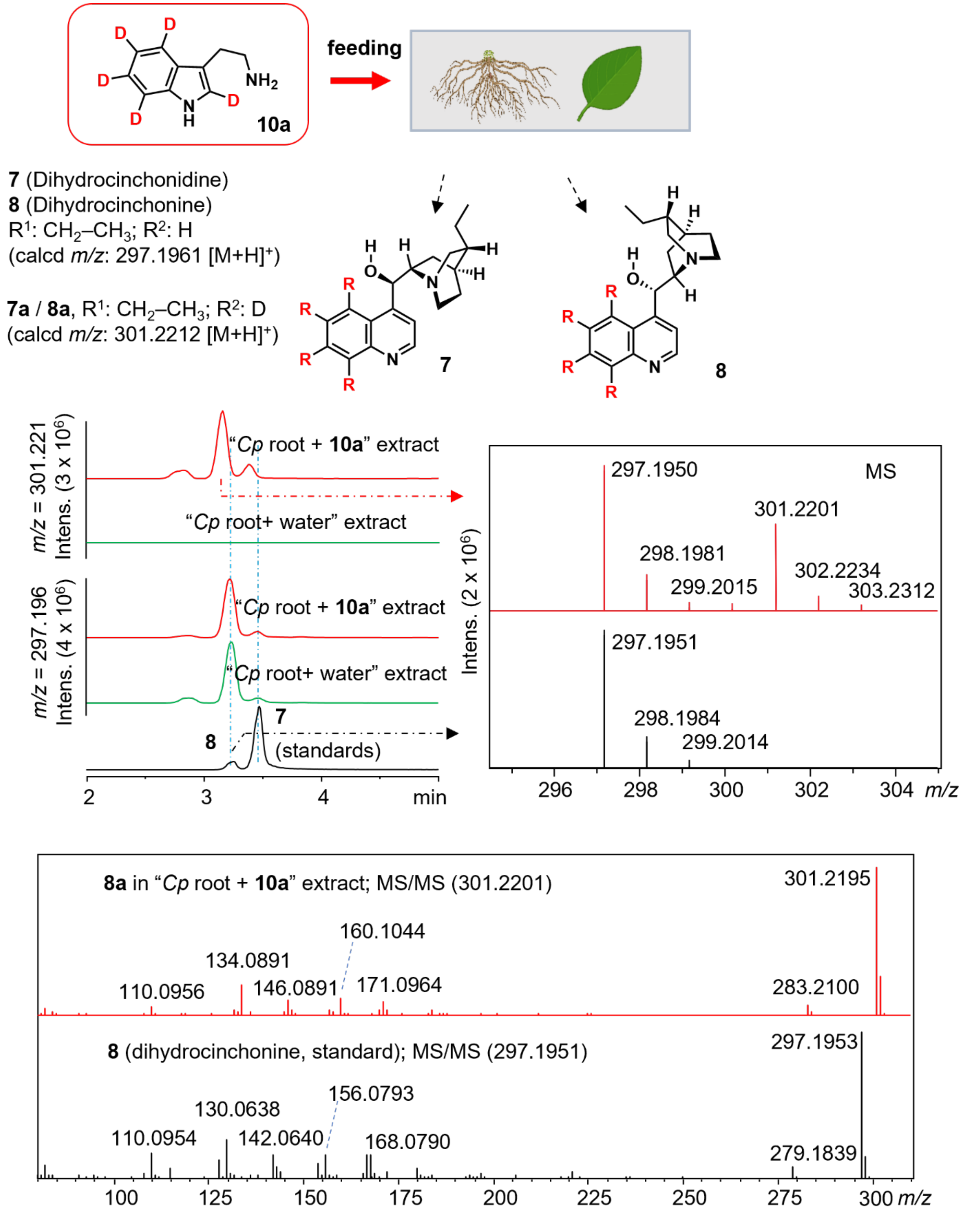


**Figure S28. Incorporation of tryptamine-(indole-*d*5) (10a) into dihydrocinchonidine (7) and dihydrocinchonine (8).** Extracted ion chromatograms, isotope patterns, and MS/MS spectra evidencing the formation of **7a** and **8a** in roots of *Cinchona pubescens* tissues fed with **10a**.

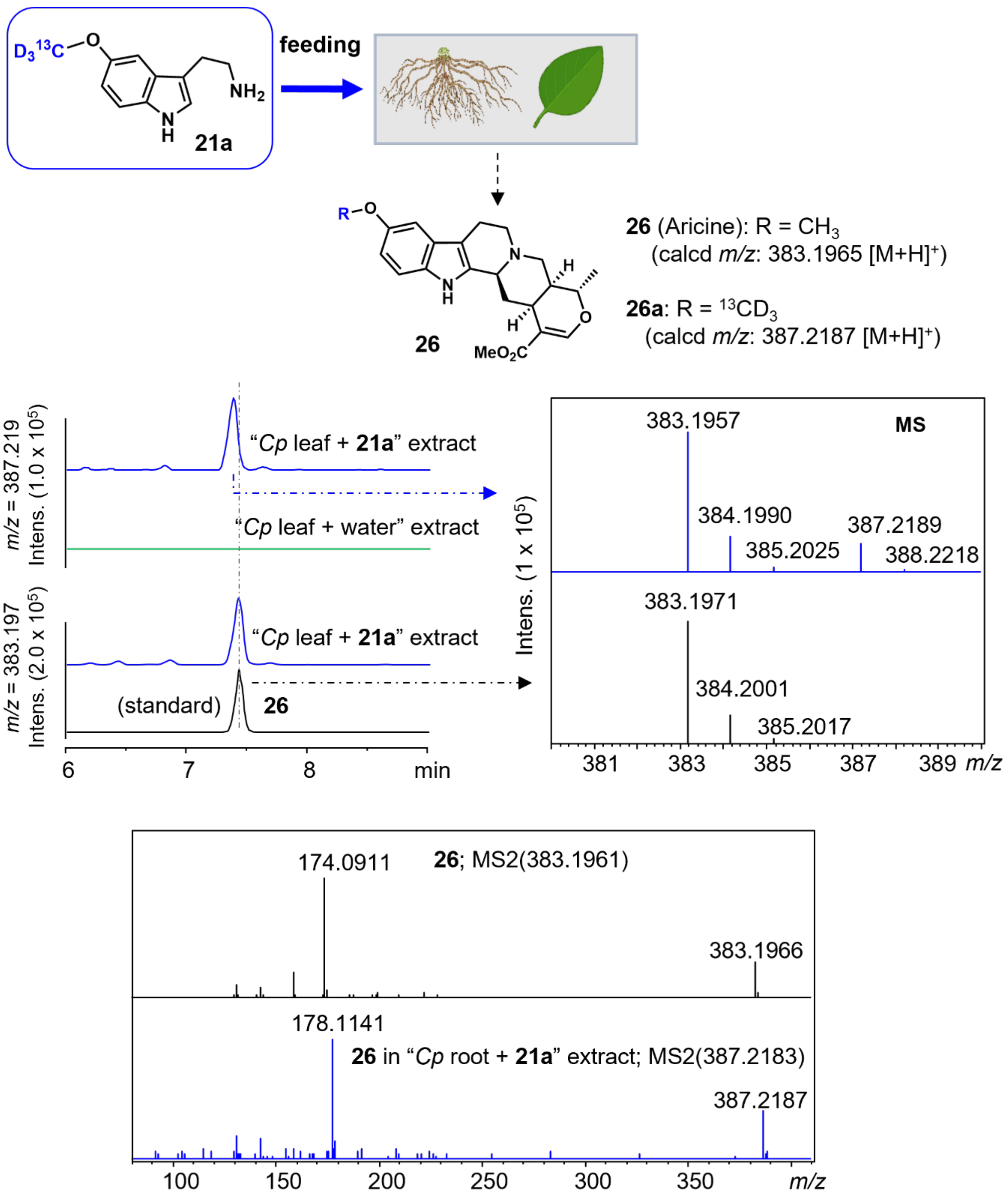


**Figure S29. Incorporation of 5-methoxytryptamine-(O-*methyl*-$^{13}$C, *d*3) (21a) into aricine (26).** Extracted ion chromatograms, isotope patterns, and MS/MS spectra evidencing the formation of **26a** in roots of *Cinchona pubescens* tissues fed with **21a**.

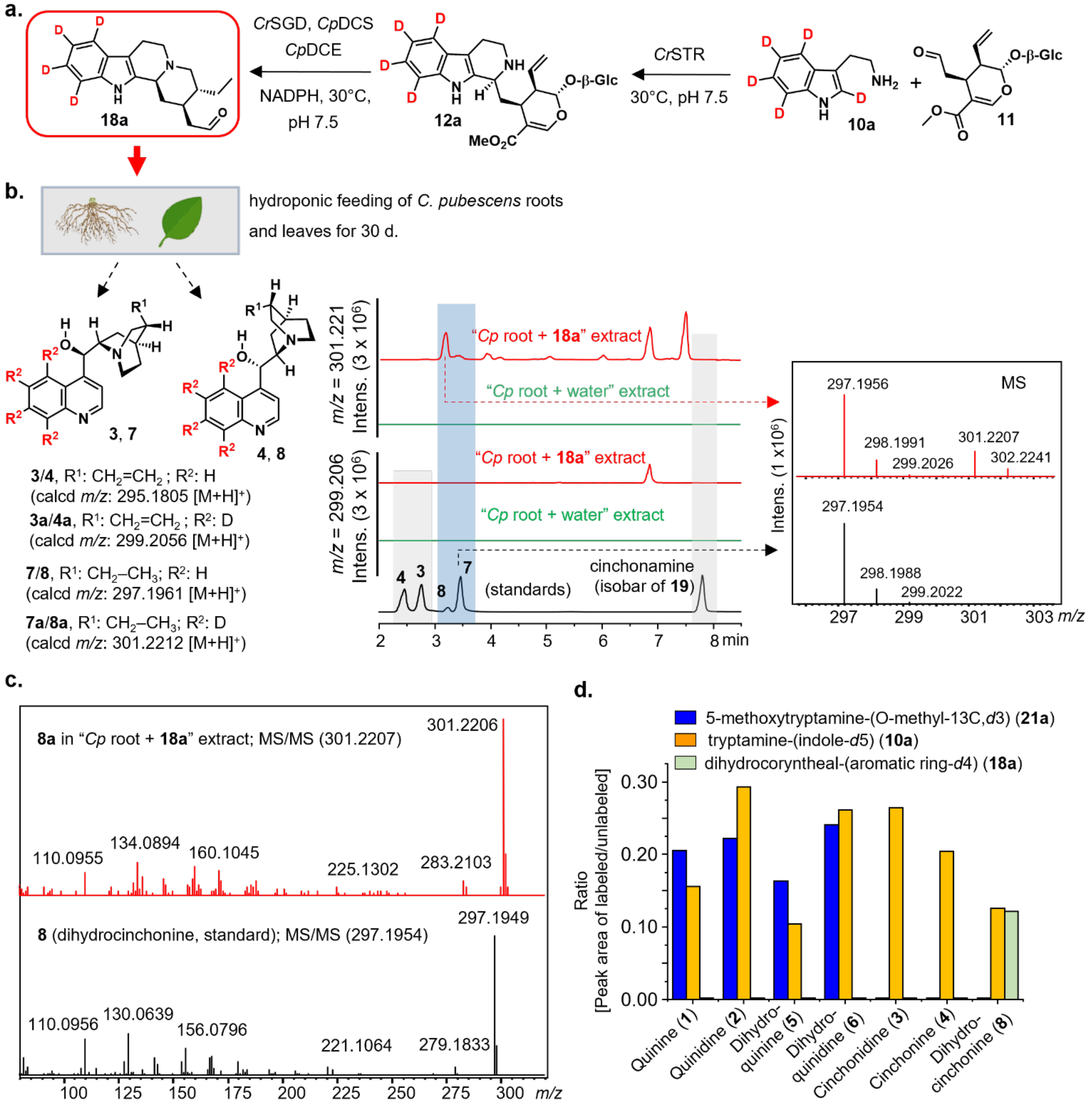


**Figure S30.** (**a**) synthesis of the labeled precursors *d*4-dihydrocorynantheal (**18a**), which was fed to the roots and leaves of *C. pubescens*; (**b**) key desmethoxylated cinchona alkaloids cinchonidine, cinchonine and dihydro analogs (**3**, **4**, **7** and **8**), along with extracted ion chromatograms of *m/z* corresponding to isotope-labeled analogs (**3a**, **4a**, **7a** and **8a**), and MS isotopic patterns, showing the incorporation of **18a** into **7** and **8**, but not into **3** and **4**; **(c)** MS/MS spectrum (20.0-50.0 eV) corroborating the formation of labeled **8a**; and **(d)** incorporation ratio of the three labeled precursors into main Cinchona quinoline-type alkaloids.

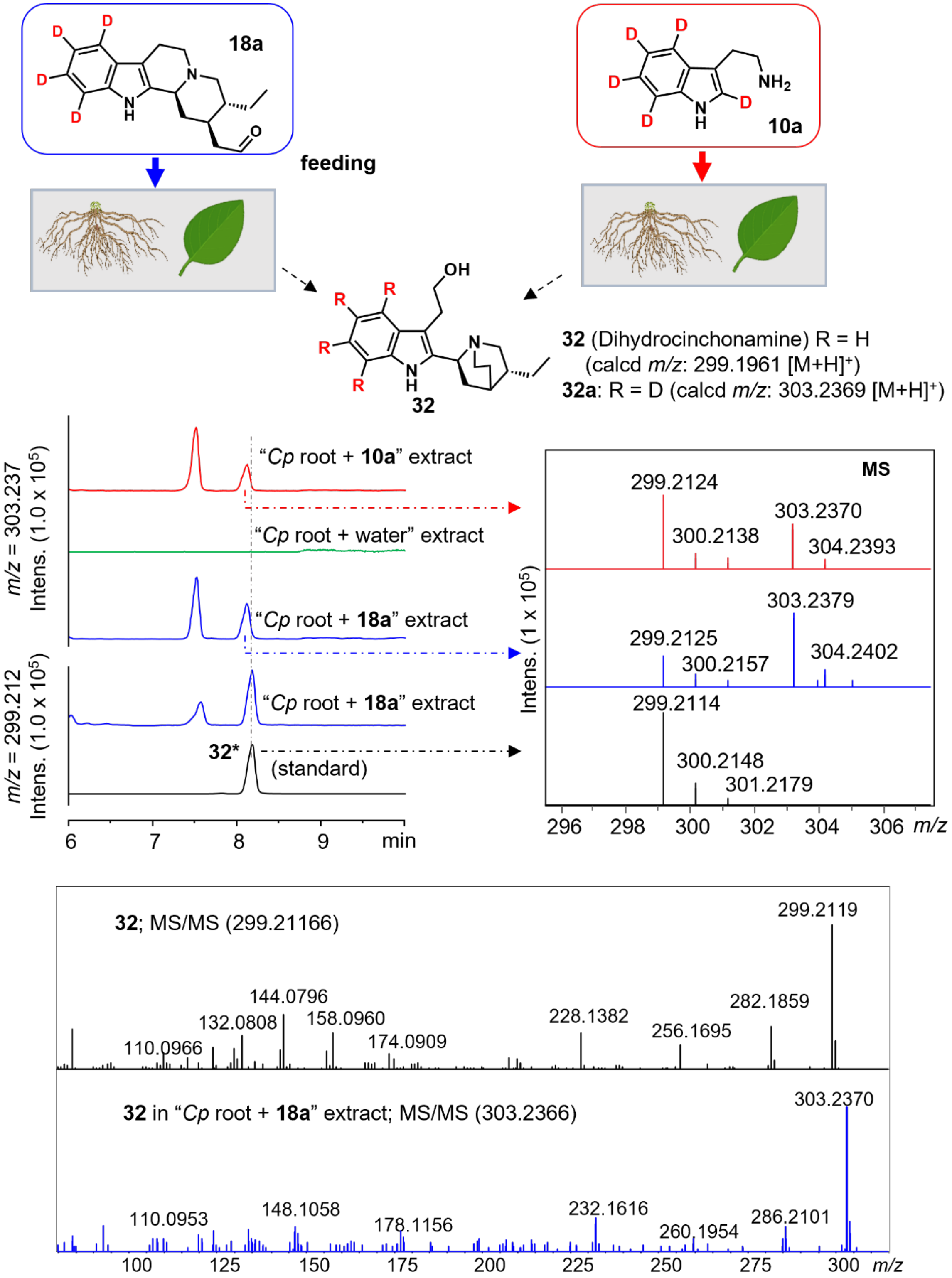


**Figure S31. Incorporation of tryptamine-(indole-*d*5) (10a) and of *d*4-dihydrocorynantheal (18a) into dihydrocinchonamine (32).** Extracted ion chromatograms, isotope patterns, and MS/MS spectra evidencing the formation of **32a** in roots of *Cinchona pubescens* tissues separately fed with **10a** and **18a**, respectively. The asterisk symbol (*) indicates that this reference compound is present in the purchased cinchonamine **19**, as already emphasized in the section Methods and Materials, but its structure has not been firmly established.

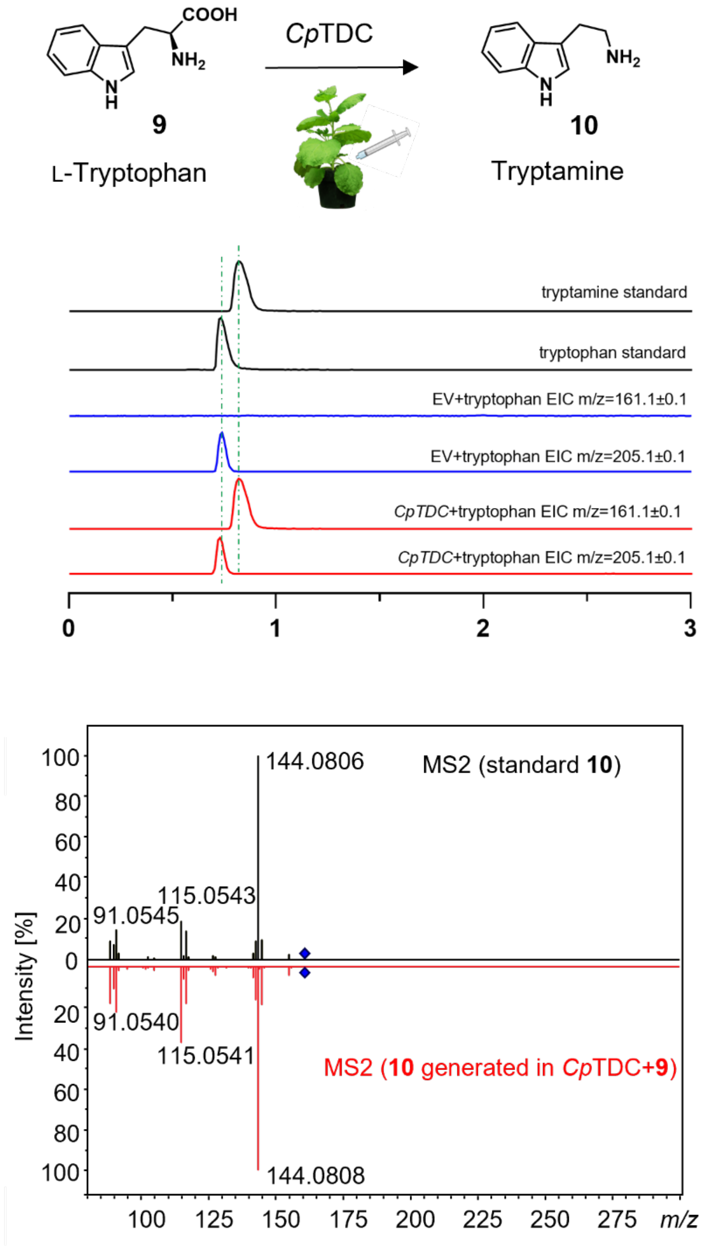


**Figure S32. Biochemical characterization of *Cinchona pubescens* tryptophan decarboxylase (*Cp*TDC) in *N. benthamiana* expression system.**

#### MS and MS/MS spectra of synthesized isotope labeled compounds

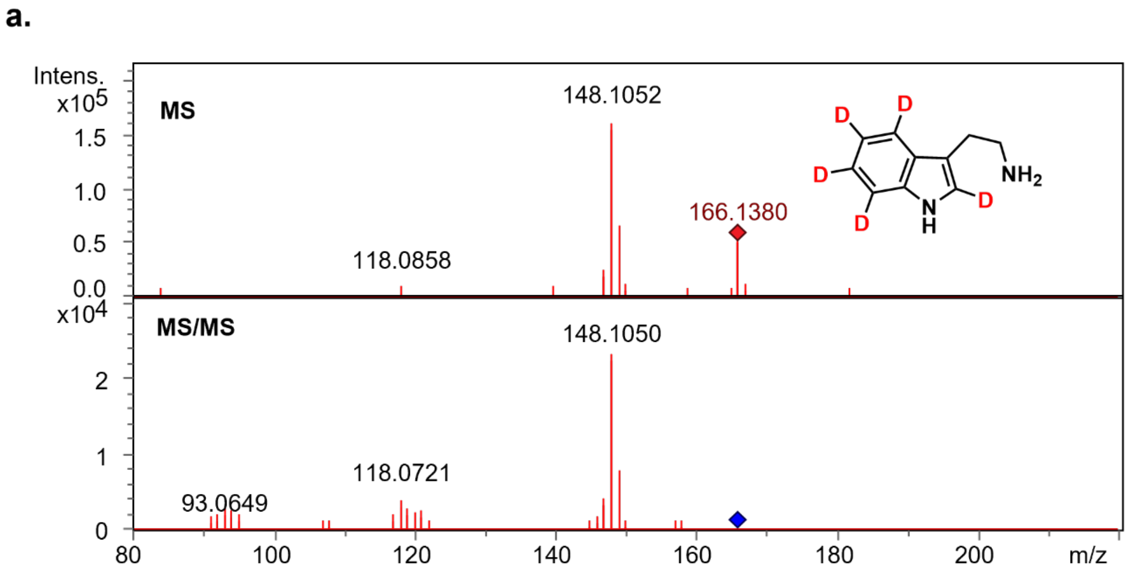


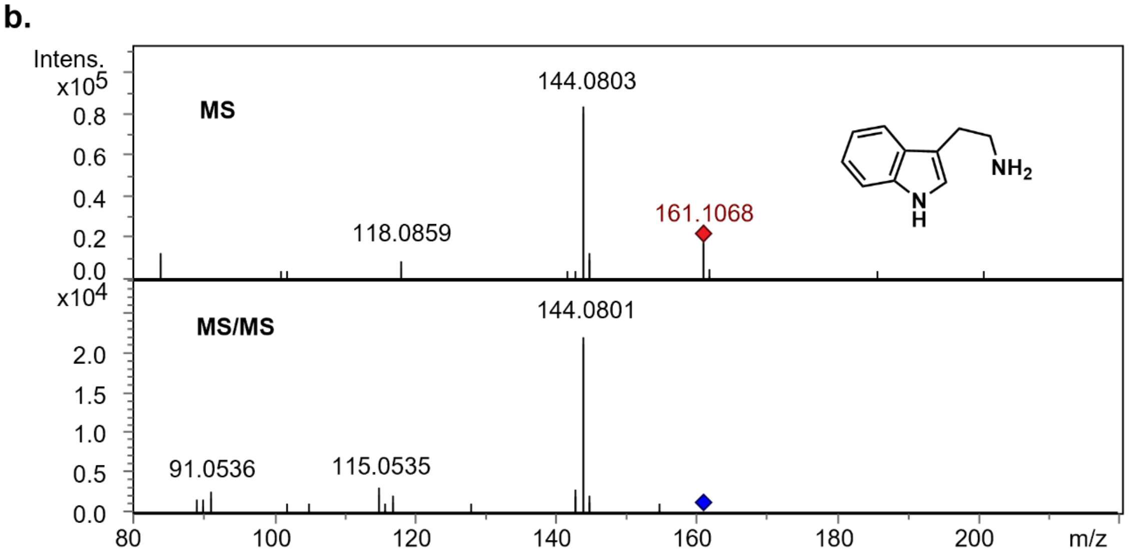


**Figure 33. Comparison of the HRESI-MS data of the synthesized isotopically labeled and of the unlabeled 5-tryptamine**. **(a)** MS and MS/MS (20.0-50.0eV) spectra of tryptamine-(*indole*-d5) (**10a**); and, for comparison, **(b)** MS and MS/MS (20.0-50.0eV) spectra of an authentic standard tryptamine (**10**).

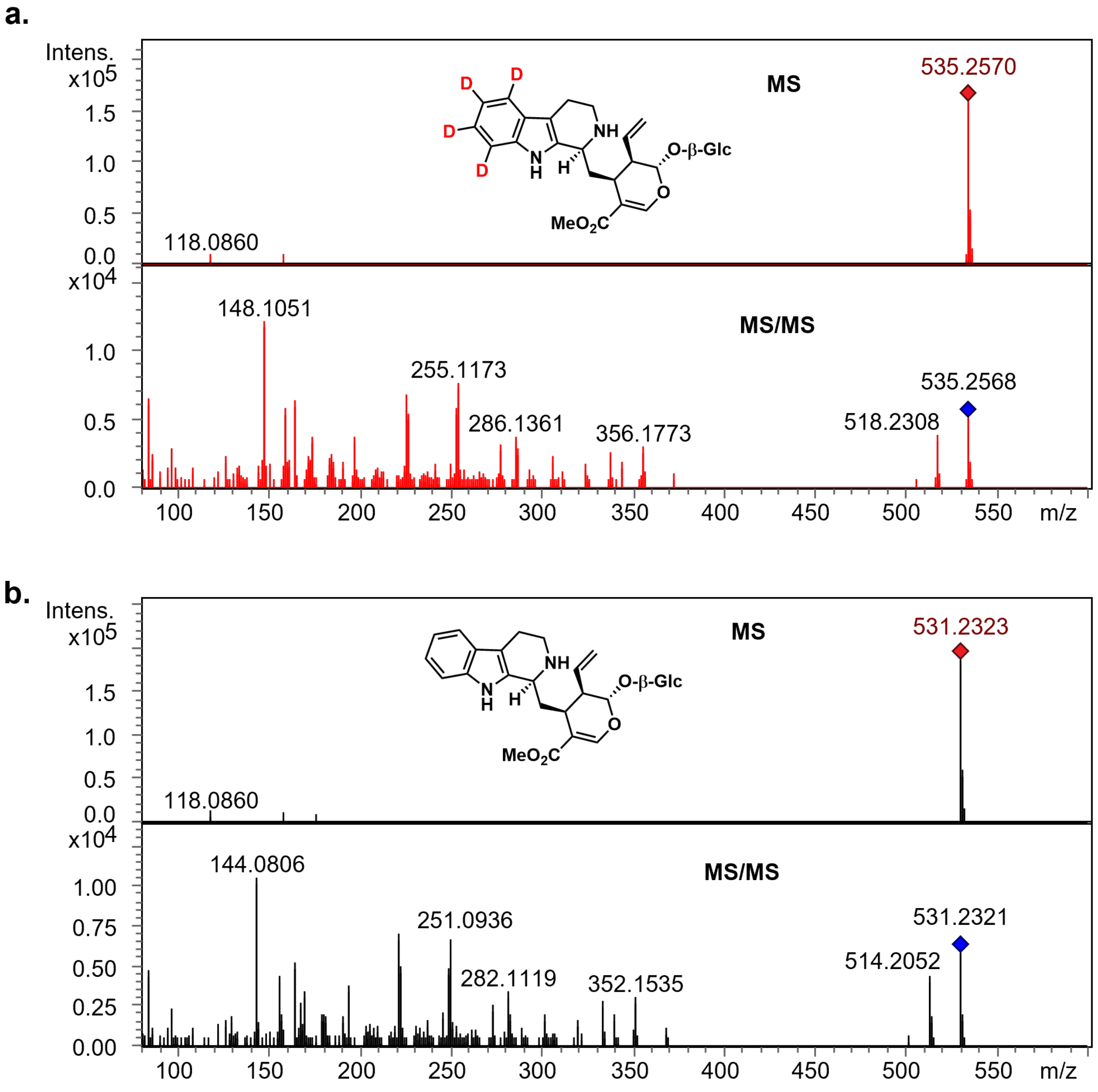


**Figure 34. Comparison of the HRESI-MS data of the enzymatically prepared isotopically labeled strictosidine and of the unlabeled standard**. **(a)** MS and MS/MS (20.0-50.0eV) spectra of *d*4-strictosidine (**12a**); and, for comparison, **(b)** MS and MS/MS (20.0-50.0eV) spectra of an authentic standard strictosidine (**12**).

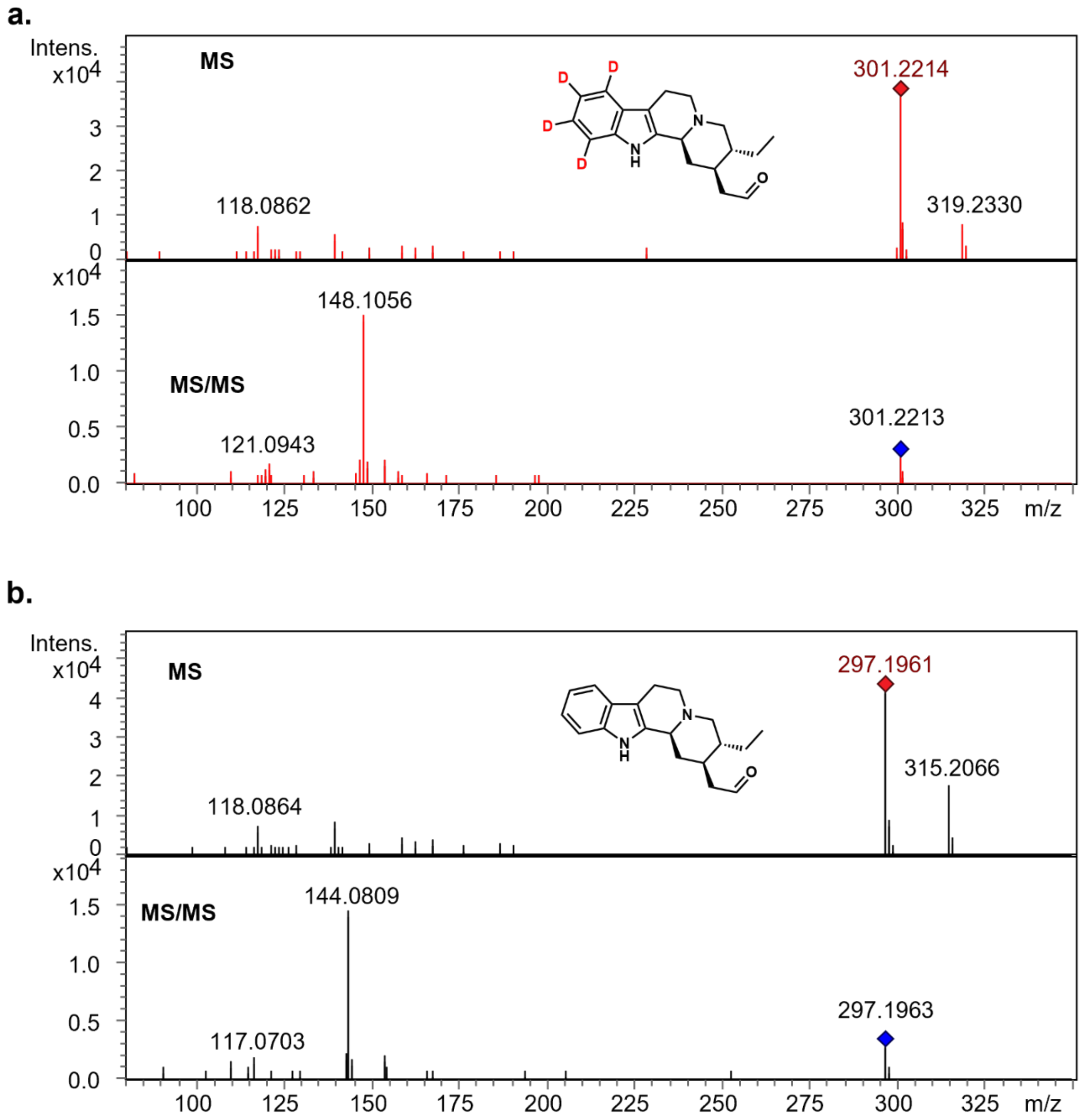


**Figure 35. Comparison of the HRESI-MS data of the enzymatically prepared isotopically labeled and of the unlabeled dihydrocorynantheal**. **(a)** MS and MS/MS (20.0-50.0 eV) spectra of *d*4-dihydrocorynantheal (**18a**); and, for comparison, **(b)** MS and MS/MS (20.0-50.0eV) spectra of an authentic standard dihydrocorynantheal (**18**).

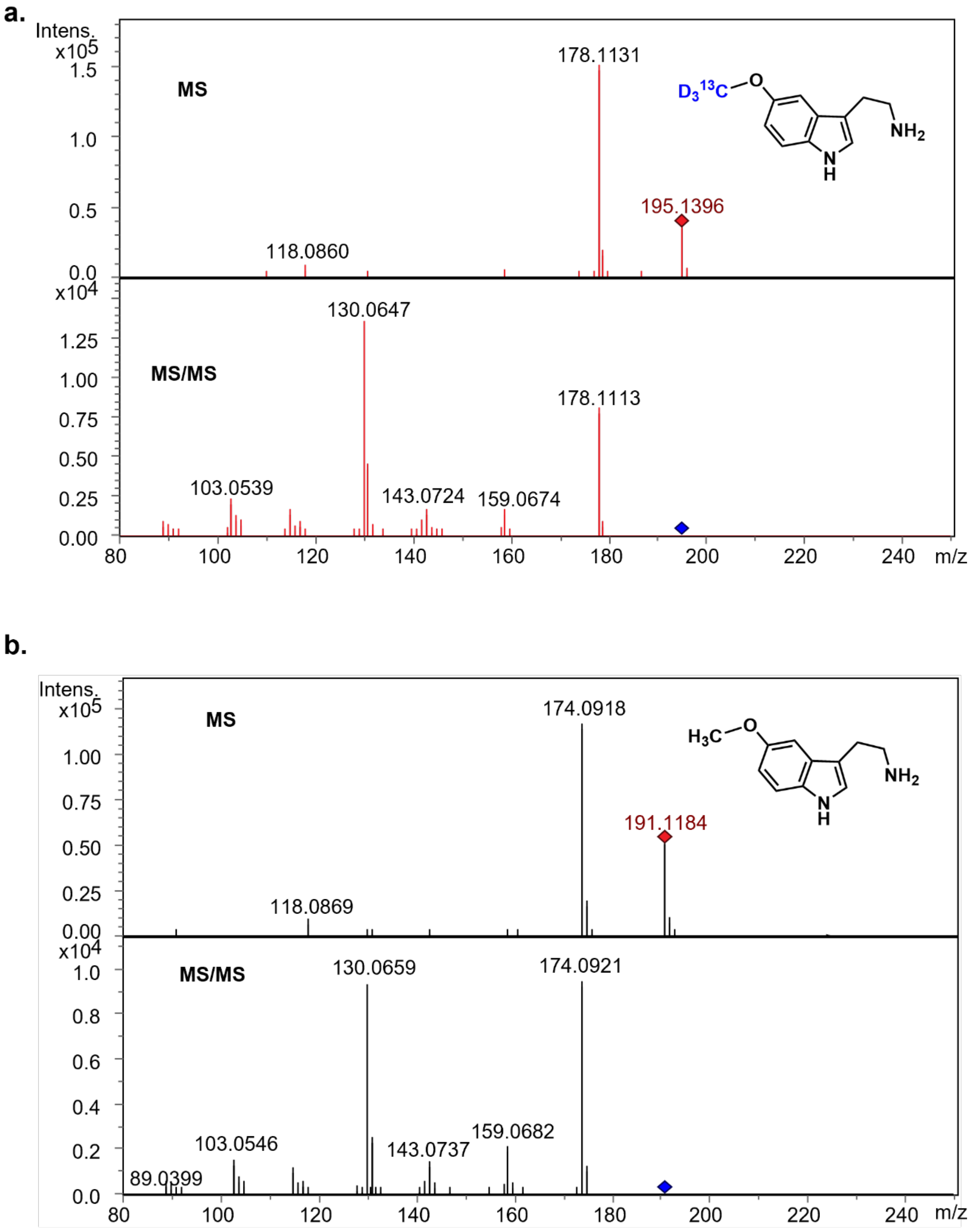


**Figure 36. Comparison of the HRESI-MS data of the synthesized isotopically labeled and of the unlabeled 5-methoxytryptamine**. **(a)** MS and MS/MS spectra (20.0-50.0eV) of 5-methoxytryptamine-(O-*methyl*-$^{13}C$, *d*3) (**21a**); and **(b)** MS and MS/MS spectra (20.0-50.0eV) of an authentic standard 5-methoxytryptamine (**21**).

### Supplementary Tables

**Table S1.** Nucleotide sequences for genes described and used in this study.

| Gene Name | Nucleotide sequence |
|---|---|
| *Cp*T5H1 | ATGGAGAATTTTACAGTTGAAATCCTCTGCATGTTTTTTCTCTCTTTTGCCA<br>CCTTTTGTTACTACTATTGTTGCAAAATTTATGCACGGACTCGGTCACAGCC<br>TGTGACTCTACCACCGTCACCCTCAAAACTCCCCATCATCGGTCACCTACA<br>CCTCCTCACCGACATGCCCCACGTTGACATGGCTCAACTCGCTGAAAAACT<br>CGGCCCAATAATCTACCTCCAACTCGGTCAAGTCCCCACTGTGGTTATCTC<br>GTCGGCTAAACTCGCCGAACTCGTTCTCAAAACTCACGATCACATAATGGC<br>GAATCGGCCCCAACTCATTGCCGCTCAGTACCTCTCATTCGGCTGCTCCGA<br>CGTCACTTTCTCCCCTTACGGCCCTTACTGGCGGCAAGCGAGAAAGGTCTG<br>CGTCACCGAGTTACTGAATTCGAAACGAGTCAACTCGTTCCAACTCGTTCG<br>AGATGAGGAAGTGAATCGCACGTTACGCACCGTGTCCGAGTCAGCTCATA<br>CAAACTCTGAAATCGACGTGAGCGAGTTGTTCTTCAAACTCGCCAACGACA<br>TCCTCTGCCGCGTAGCATTCGGGAAGAGGTTTATGGACGAGACGAGTCATG<br>AAGGAGGGAAGAGGAACGAGTTGGTCCGAGTCTTGACGGAAACGCAGGCT<br>TTGCTAGCTGGGTTTTGCATCGGAGACTTTTTTCCGAGTTGGAAGTGGGTTA<br>ATTCAGTGAGTGGGATGAAGAGAAGGTTGATGAATAATTTGAAGGATTTG<br>AGAATGGTTTGTGATGAAATAATTAATGAGCATTTGAGAAAGACAGAGCA<br>TACAGGTTCAAGCGGCGTCGTTTCGGAGAAAGAAGATTTTGTGGATGTTTT<br>GCTCAGAGTTCAAAAGCAAGATGATCTTGAAGTTCCTATTACTGATGATAA<br>TCTCAAAGCTCTTGTTATGGACATGTTTGTAGCTGGGACAGATACAACATC<br>AGCAACATTAGAGTGGACAATGACAGAACTGGCAAGGCATCCTAATATAA<br>TGAAAAGAGCACAAGAAGAAGTAAGGCAAATTGCAGCAAGCAAAGGAAG<br>AGTAGAAGAAACTGACCTTCAGCATCTTCATTACTTGAGAGCTGTGATCAA<br>AGAAACCATGAGACTTCATCCACCAGTCCCTCTTCTCGTTCCACGCGAATC<br>TCTTGCCAAATGTACGATTGATAAGTATGAAATACCAGCGAATACTCGGGT<br>TTTGATCAACACTTACGCTATTGGGAGGGATGCTGATTCATGGGAGAACCC<br>TTTGGAATACAATCCTGAAAGGTTTGTAGGGAAAGATGGTATTGATTTTAA<br>GGGTCAAGATTTCAGGTTTTTGCCATTTGGAGGTGGAAGAAGAGGTTGCCC<br>TGGTTTCTCCTTTGGGTTGGCAAGTGTTGAGATTTCGCTAGCCCGTTTACTG<br>TATCATTTTGACTGGAAATTGCCTCAAGGGGTTGGAGCAGATGATGTTGAT<br>CTTACAGAGATTTTCGGACTTGCTACTAGGAAAAGATCAGCACTAAAATTG<br>GTTCCTACAATGATCAAGAATTAA |
| *Cp*T5H2 | ATGGAGAATTTTACAGTTGAAGTCCTCTGCATGTTTTTTCTCTCTTTTGCTA<br>CCTTTTGTTACTACTATCGTTGCAAAATTCATGCACGGACTCGATCACAGCC<br>TGTGACTCTACCACCGTCACCCTCAAAACTCCCCATCATCGGTCACCTACA<br>CCTCCTGAGAGACATGCCCCACGTTGACATGGCTCAACTCGCAGAAAAACT<br>CGGCCCAATCATCTACCTCCAACTCGGTCAAGTCCCCACCGTGGTTATCTC<br>GTCGGCTAAACTCGCCGAACTCGTTCTCAAAACTCACGATCACATAATGGC<br>GAATCGGCCCCAACTCATTGCCGCTCAGTACCTCTCATTCGGCTGCTCCGA<br>CGTCACTTTCTCCCCTTACGGCCCTTACTGGCGGCAAGCGAGAAAGGTCTG<br>CGTCACCGAGTTACTGAATTCGAAACGAGTCAACTCGTTCCAACTCGTTCG<br>AGATGAGGAAGTGAATCGCACGTTACGCACCGTGTCCGAGTCAGCTCATA<br>CATACTCGGAAATCGACGTGAGCGAGTTGTTCTTCAAACTCGCGAACGACA<br>TCCTCTGCCGCGTGGCATTCGGGAAGAGGTTTATGGACGTGGCGAGTCATG<br>AAGGAGGGAAGAGGAACGAGTTGGTCCGAGTCTTGACGGAAACGCAGGCT<br>TTGCTAGCTGGGTTTTGCATCGGAGACTTTTTTCCGAGTTGGAAGTGGGTTA<br>ACTCAGTGAGTGGGATGAAGAGAAGGTTGATGAATAATTTGAAGGATTTG |

| | |
|---|---|
| | AGAATGGTTTGTGATGAAATAATTAATGAGCATTTGAGAAAGATAGAGCA<br>TACAGGTTCAAGCGGCGTCGCTTCGGAGAAAGAAGATTTTGTGGATGTTTT<br>GCTCAGAGTTCAAAAGCAAGAAGATCTTGAAGTTCCTATTACTGATGATAA<br>TCTCAAAGCTCTTGTTCTGGACATGTTTGTAGCTGGGACAGATACAACATC<br>AGCAACATTAGAGTGGACAATGACAGAACTGGCAAGGCATCCTAATATAA<br>TGAAAAGAGCACAAGAAGAAGTAAGGCAAATTGCAGCAAGCAAAGGCAG<br>AGTAGAAGAAACTGACCTTCAGCATCTTCATTACTTGAGAGCTGTGATCAA<br>AGAAACCATGAGACTTCATCCACCAGTCCCTCTTCTCGTTCCACGCGAATC<br>TCTTGCCAAATGTACGATTGATAAGTATGAAATACCAGCGAATACTCGGGT<br>TTTGATCAACACTTACGCTATTGGGAGGGATGCTGATTCATGGGAGAACCC<br>TTTGGAATACAATCCTGAAAGGTTTGTAGGGAAAGATGGTATTGATTTTAA<br>GGGTCAAGATTTCAGGTTTTTGCCATTTGGAGGTGGAAGAAGAGGTTGCCC<br>TGGTTTCTCCTTTGGGTTGGCAAGTGTTGAGATTTCGCTAGCCCGTTTACTG<br>TATCATTTTGACTGGAAATTGCCTCAAGGGGTTGGAGCAGATGATGTTGAT<br>CTTACAGAGATTTTCGGACTTGCTACTAGGAAAAGATCAGCACTAAAATTG<br>GTTCCTACAATGATCAAGAATTAA |
| *Cp*OMT1<br>(GenBank accession:<br>MW456557.1) | ATGGCAACATCTGAGAATTCTGCTGAGCTTATGAGAGCTCACAATCTTATT<br>TGGAACCAAACATTCAACATCAAGAATTCAGCCTGTCTAAAATGTGCAATT<br>CAACTAGGCATACCAGACGTCATCCACAAGCATGGAAAGCCCATCGCTCTC<br>GCTGACCTGACTTCTGCCCTTCCAATTAACCCTTCTAAAGCTCCGTACATCA<br>AACGCTTAATGCAAATTCTAAAAGATGCTGGGTTTTTTGCTCAAGAAAAAG<br>AGGGTTATTATGCTCTTACTTTTGCGGGCCGCCTTCTTGTCGAAAATGAGCC<br>AATGAATGGAAGAGAGTTCGTTCTTATGAATCTTGATGCTGCTATGATGAA<br>GCCTTGGATTGTGTTGAGTGAGTGGTTTCAGAATGATGATCATGCTCCATTT<br>GACACTGCTCATGGGAAGAATTTTTGGGATCATAATGCCGATGAACCAAA<br>AGTTGGCAAACTATTTAATGAAGCTATGGCTAGTGATTCTCAGATGGTTAC<br>GACGGTGCTGACTAAAGAATGTGGATATGTGTTTGAAGGGTTGACATCTTT<br>GGTGGATGTTGGTGGTGGCACAGGCACAGTTGCTAGGTCCATTGCTAAAAT<br>GTTCCCGAATTTAAAATGCGCTGTGTTTGATCTCCCACATGTGATTGCCAAT<br>CAAGAAGGAACTCAGAACTTGGATTTTGTTGCAGGAGATATGTTCGAGAA<br>GGTGCCCCCAGCTAATGCAATCTTACTTAAGTGGATTCTCCATGACTGGGG<br>TGATGAGGAATGCATAAAGATTCTCAAGAACTGCAAAAAGGCAATTCCAG<br>GAAGAGACGAAGGGGGAAAATTGATCATCATAGAGATGGTTATGGAAAGC<br>CAGATAGAGGATGAGGATTCAGTTGAAGAGCAAATTTGCTCGGACTTGCA<br>AATGTTTGTTTTATTCCGTACTAAAGAAAGAACAGAGAAAGAATGGGCAA<br>CACTTTTCTGGAATGCTGGATTCAGCGACTATAAAGTATTTCCAGTATTGG<br>GTGCAAGGAGTATCATTGAGGTTTATCCTTAA |
| *Cp*STR | ATGCACATTTCTGAAAATATGTTCGTCGTCACCATTTCCTTCATCCTTTTCTT<br>GTCATCTCCTTCACTCGTTCTCTCTTCTCCATATTTCCAATTTATTCAAGCAC<br>CATCCTACGGCCCCAACGCCTATGCTTTTGATTCAGCTGGTGGACTCTATGC<br>TGTCGTAGAGGATGGTAGAATTGTGAAGTATGAAGGATCAAGCAATGCAT<br>TCTTGGACCATGCTGTTGCCTCTCCATTCTGGACTAAAAAACTGTGTGAGA<br>ACAACACTAAACCTCAGCTAAAACCCCTGTGTGGGAGGGCATATGACCTC<br>GGATTCCACTATGAAACTCAGCAATTATACATTGCTGATTGCTATTTTGGTC<br>TTGGTGTTGTTGGACCTGAAGGAGGGCTTGCAAAAAAGCTTGCCAAAAGT<br>GGAGATGGTGTGGAATTCAAGTGGCTTTATGCCTTGGTTGTGGACCAGCAA<br>ACTGGCTTTGTTTACGTCACCGATGTTAGCACAAAATATGATGACAGGGGT<br>GTTCAAGATATCCTAAGGACAAATGATACAACAGGAAGATTAATCAAATA<br>TGATCCCACAACCAGAGAAGTTACAGTTTTGATGAAAGGCCTAAATGTACC<br>AGGTGGTGCGGAAATTAGCAAAGATGGCTCTTTTATTCTTATAGGTGAATT<br>CTTAAGCAACCAAATTCTCAAGTATTGGCTAAAGGGTCCCAAAGCAAATAC |

| | |
|---|---|
| | TTTAGAATTCTTGTTACATGTTAAGGGTCCAGGTAGTATTAGGCGGACTAAGGCTGGAGATTTTTGGGTGGCTTCAAGTGATAATAATGGAATTACGGTTACTCCTAGAGGAATAAGGTTTGATGAATCTGGCAACATTTTGGAAGTTGTGCCTATTCCTCTACCATACAAAGGTGAACATATTGAACAAGTTCAAGAACACAATGGTGCACTCTACATTGGATCTTTGTTCCATGGTTTCATAGGTATATTGTACAATTACAAGGGTTTATCAGAGGAAAATAATCTAGGTGGGGTCGTTGAATCATTGAAAGGAGAGTCGTTTTCTTTCTGA |
| *Cp*STTr | ATGGAAGCTGCAGTGATGTCTTCTTTGGACGACGACGCTGAAACTCAACTGTTACAACAACCCAGTCCCAAGACTAAGAAAGGCGGTTGGATCACCTTCCCATTTCTTATAGCAACCAGGGCTGGCATGACGGTTGCAGCATTAGGATGGAGCGCCAATCTTATTGTCTACCTCATTGACAAGTACAATATCGAGAGTATTGATGCTGCACAAATCTTCAATGTAGTCAATGGCTGCATGGCACTTTTTCCTATTATCGTAGCTATAATTGCAGATAGTTTTCTTGGCTGCTTTGCTGTCATCTGGATTTCTTCAATCATCTCTTTGCTGGGAATGGTTCTGTTGACTCTAACTGCAACGATTAGTTCACTAAGACCTGCACCATGTAATGAAGGGTCGAGCTTTTGCACAACTCCATCACCATTGGAATATACAAACCTATTTTTGGCTGTGGCTCTGGCATCTATAGGCTGTGCAGGTACTAGTTTCACAGTCGGAACAATGGGAGCAGATCAACTGGATAATCCTGAGCATCAAGAGAATTTCTTTAACTGGTTTCTCTTTGTTTGGAATGCTGCTTCAATAATTAGTGCTACTGTGATTGTCTATGTTCAAGATAATGTGAGTTGGGGACTGGGGTATGGACTATGTGCTGCAGCAAATTTGTTAGGATTAATTAGTTTTTTGCTGGGAAAGCATTACTATCGCTATGTTCAGCCACAAGGGAGTCCATTCAAGGATATAGCTCGTGTACTATTTGCGGCCTTCTCTAAGAGGAAGGTCTTTTTGTCAACAAGAAATGAAGATTATTTTAGTGAATTACATCTTGAAGTCGATGGCCAACATGATGGTATCAAAGAATTGGCAGCATCAGCAACACCACATAAAGAAACATTCAAGTTTCTGAACCATGCGGCCTTAATAAGCCAAGCAGACATCCAATCAGACGGATCAATCAGGCAATCGTGGAAGTTGTGCACAGTACAACAAGTGGAAGATCTTAAAACCTTACTTAGAATTGCCCCAATTTGGGCAACCGGTATTTTCTTAACCACACCAATGGGTATGCTATCTACCTTAACAGTCCTTCAGGCTCTAACAATGGACACTTGTATTGTATCCAATTTCAAATTCCCGGTAGGTTCCCTGGTAGTCTTTTCACTACTTTCTGGTGCCATCTCTCTCACCATAGTAGACCGATTGATATTCCCTTTATGGCAAAAAACATTTGGAAAAACTCCAACACCCCTCCAACGACTAGGCACGGGTCATGTCTTGAATGTACTGAGCATAGTCATTGCAGCCCTGGTGGAATCAAAGCGGCTCCAAATAGCTCGAGCCAGCTACATTGTCCAAGAATTGACCAGTTCCACCGTGCCAATGTCTGTTTTCTGGTTAGTTCCGCAGCTTGCACTTTCGGGAATGGGAGAAGCATTTCATTTTCCAGGACAAGCTTCATTATACTATCAAGAATTCCCTGCATCCCTTAAAAGCACGTCGACTGCAATGGTTGCACTGCTTATAGCAATTGGATACTATTTGAGCACGGCCTTGACAGATTTTGTGCGGAAGGTAACGAATTGGTTGCCAGATGATTTGAATCATGGAAGGCTCAACTATTTGTATTGGGTGCTGGCTATGATTGGTGCATTGAATTTTGGCCTTTATTTAACATCTGCTGGGGTCTACAAGTATAGAAATGATCAGGATGACAAGACGGTCGATAATAGTTCAAACCAAGGGGATGCAACGTTGTACTACTAG |
| *Cp*THAS1 | ATGGCTGAAAAATCAGCAATAGAAGCAAAGCCAGTGGAGGCCTTCGGATGGGCAGCTAGAGACCCTTCTGGTCTCCTCTCTCCTTTCAAGTTCTTAAGAAGATCAACTGGTGAGCATGATGTGCAGCTTAAAATATTGTATTGTGGGATCTGTGATTGGGATCTGAATGTGGTCAGGAATGGCTTTGGGACAACTAACTATCCTATCGTGCCTGGGCATGAGGTTGTGGGTGAGGTAACTGAAGTGGGTAGCCAAGTGCAGAAATTCAAAGTAGGGGATAAAGTAGGCGTGGGTGCCTTGGTGGGCTCTTGTGGAAAATGCAGGAACTGTATGGAAGGTCTTGAAAACTACTGCCCAAGAATGAAAACAAGTGATGGCACATGTTATAGTGATGGAAACGCGACA |

| | |
|---|---|
| | TTTTTTGACCCGACAGGCCACATAGTAACAACGGATAATGCCAATGATCAT TTAACCAACAAGATATATGGTGGCTATTCAAATGTCATGGTAGTCGATGAG CAGTTTGTGATACTTTGGCCTGACAACTTAGATCTACTACGGGCCGGGCCT CCTCTACTTTGTGCTGGCATTGTTCCATACAGCCCGATGAGATTCTTTGGAC TTGATAAACCTGGAATGCATATTGGTGTTGTTGGTCTCGGTGGGCTTGGCC ATCTTGCTGTCAAATTTGGTAAAGCTTTTGGGTCACATGTCACGGTCATAA GCACATCCATTAGCAAGAAGCAGGAGGCCATTGAAAAATATGGTGTAGAC GCATTTTTGCTAGTTAGTGATACCAACCAGATGCAGGCTGCAGCAGATTCG ATGGATGGGATTATCGATACTGTTGGGAAAATTCATCCTCTTTTGCCATTA ATCAATTTGTTAAAGCGTGACGGGAAGTTAGTTTTGCTTGGTCCACCAGAA GAACCATTCGAGTTGCCAGCTGCTCCTCTGATTATGGGGAGGAAGATGGTG GTTGGGAGTGTCTCGGGAAGCATAAAGGAAACACAGGAGATGGTAGATTT TGCAGCAGAACACGGCATAGTGCCTGATGTGGAGATTATTCCGATCGACTA TGTAAACACTGCAATGGAGCGCATAGAGAAAGGAGACGTCAAATATCGAT TTGTAATTGACATTGGAAAAACGCTCAAATCTGATTGA |
| *Cp*THAS2 | ATGGCTGAAAAACCAGCATTCGGAGTAGCAGGACAGCGAGTGGAGGCCTT TGGATGGGCAGCTAGAGACGCTTCTGGCGTCCTCTCTCCTTTCAAGTTCTCA AGGAGAGCAACGGGCGACCATGATGTTCAGTTCAAAGTGTTGTATTGTGG GATTTGCGATTGGGATATGAACGTGGTGAGAAATGTGTTTGGGACGACCA ATTATCCAGTCGTGCCTGGGCACGAGGCTGTGGGCGAGGTAACTGAAGTG GGTAGACGAGTGCAGAAGAATTTCAAAGTTGGAGATAAAATAGGCTTGGG TGGGTTGGTTGGAACGTGTGGCAAATGCAGGAGCTGTATAGAAGGTCTTG AAAATTACTGCCCGAGCCTGAAAAGTGCTGATGGCACTGGCTTCAGCATCG GAGACATGGTATTTTTTGACGGTACCAATAATCGTGTAACCGACAACAAAC AATATGGTTGCTACTCAAATATTATGGTGGTTGATGAGAAGTTTGCGATTC GTTGGCCCGAAAACTTGGATCTCCGAGCCGGACCTCCTCTGCTTTGTGCTG GGATTGTTCCATATGCCGCGATGAGATACTTCGGACTTGATAAACCAGGAA TGCATGTTGGTGTTGTCGGTCTTGGTGGGATTGGCCATCTTGCTGTCAAATT TGGTAAAGTTTTTGGGGCACATATAACGGTCATCAGCACGTCCATTAGCAA GAAGCAAGAGGCCATTGAAAAATATGGTGCCGACTCATTTTTGCTCGTTAG TGATTCTAGCCAGATGGAGGCTGCAGCTGATTCGATGGATGGGATACTTGA TACTGTTGGGAAAATTCATCCACTTTTGCCATTGATAAATATGTTGAGACG TGACGGGAAGCTGGTTTTGCTTGGTCCACCAGAAGAGCCGTTTGAGTTGCC AGCAGCTCCTCTGATTATGGGGAGGAAGACGGTGGTTGGGAGTACCGCGG GAAGTATAAAGGAAATACAGGAGATGGTATATTTTGCAGCAAAGCACAAC ATACTGCCTGATGTGGAGATAATCCCGATTGACTATGCAAACACTGCAATG GACCGCATTGAGAAAGGAGATATCAAATATCGATTTGTAATCGACATTGG AAACACGCTTAAATGTGATTGA |
| *Cp*TDC | ATGGGCAGCATTGATGCTAATAACGGTGCTTATACAGCTTCACCTGTTGCC CCATTCAAGTCACTTGATCCTGAAGACTTCAGAAAACAAGCCCATCGTATG GTGGATTTCATAGCCGATTATTACAAAAACATCGAAAACTATCCAGTTCTC AGCCAAGTTGAGCCCGGATATCTCCGAAGCAAACTGCCCGAAACCGCCCC TTACCTGCCCGAATCGTTCGAGAACATTTTGAATGATATTCAGAAAGATAT AATCCCTGGAATGACCCACTGGTTGAGCCCTAACTTTTTTGCATACTTTCCA GCTACAGTTAGCTCTGCTGCCTTTCTTGGAGAAATGTTGTGCACTGGCTTTA ACTCTGTGGGGTTCAACTGGCTTGCTTCCCCGGCGGCGACCGAGCTGGAGA TGGTGGTGATGGACTGGCTGGCTAATATGCTTAAGCTCCCTGAGTCCTTTA TGTTTTCAGGCACTGGTGGTGGTGTTCTCCAAGGAACCACCAGTGAGGCTA TTCTTTGTACACTAATTGCCGCCCGCGACCGTGTTCTTGGGGAAATCGGCG TCGAGAATGTTGGCAAGCTTGTTGTTTATGGATCTGATCAAACACATTCTTT |

| | |
|---|---|
| | TTTCATCAAGACATGCAAATTGGCGGGCATTTTCCCATGCAATATAAGAAT AATCCCTACTACAGCTGAAGCCAACTTTTCCATGTGCCCTAGTGCTCTACGT AAACAAATTGAAGCTGACGTCGCAGATGAGCTGGTCCCACTTTTTCTTTGT GCTACTGTGGGTACCACTTCCACCACTGCCATTGACCCAGTGAGTCTGCTA GCTGAGGTAGCAAATGAATTCAATGTGTGGATTCATGTTGATGCTGCTTAT GCTGGTAGTGCGTGCATATGTCCTGAGTTTCGGCAATACTTGGATGGCGTT GAGCGAGTTGACTCGTTGAGTTTGAGCCCTCATAAATGGTTTCTTTGTTTTT TGGATTGCTGTTGTTTGTGGGTTAGGAAACCCGACTTGATGGTCAGGGCAC TAAGTACTAATCCTGAGTACTTGAAAAATAAACGAAGCGAATTGGATAAT GTTGTAGATTTTAAAGATTGGCAAATTGGTACTGGCAGACGATTCAGAGCA CTCCGACTATGGCTAATAATGCGCAGTTATGGTATTGCAAATCTCCAAAGC CACATCCGATCGGACGTTCAAATGGCCAAAATGTTCGAAGGGTTCGTGAA ATCCGACCCAAGGTTTGAAATAGTTGTACCACGTGCATTTTCACTTGTATG CTTTAGGCTTAATCCTTTGGGAGGAAACAACGCAATTTACTTGGAGCTTTT GAACAAGAAACTACTCGAGCTGATCAACTCAACTGGCCGAGTTTACATGA CCCACACCAAGGTTGGTGGGGTCTACATGTTGAGATTTGCAGTGGGGGCCA CGTTTACAGAGGATCGCCATGTGTATGCTGCTTGGGAGTTGATAAAAGAGT GCACTAATGCCTTGCTTAAGGAGAATCATTAG |
| *Cp*DCS (GenBank accession: MW456554.1) | ATGGCCGGAAAATCTCAAGAAGATGGGCAGACGGTAAAGGCTCTAGGATG GGCCGCTAGGGAAGTTTCTGGGGCGATCTCTCCTTTCGATTTCTCAAGAAG GGCCCCAGGAGAGCGCGATGTGCAGGTTAAAATACTATATTGTGGAATCT GTAGTTTTGACACAGAAATGATCAATAACAAGTTTGGCTTTACCAGATATC CCTTTGTACTCGGGCATGAGATTGTGGGAGTGGTATCTGAAGTTGGTAGAA AGGTGCAAAAATTCAAGATTGGGGATAAAGTTGGTGTAGGAACCATGATT GGATCTTGTCGCACTTGTTATAGCTGCACTCACAATCTCGAAAATTACTGC CCAAAAGTTACATTAACAGAAGCAACTTCTGGTGGTTGTTCTAATCTTGTG ATAGCAGATGAAGACTTTGTGTTCCATTGGCCGGTGAATTTGCCTCTTGAT CTTGGAGCTCCTCTCCTTTGTGCTGGGATTACTGTTTATAGCCCTTTGAAAA ATTTTGAACTTGATAAGCCTGGATTGCGTATTGGTGTGGTTGGTCTTGGTGG TATTGGCCATATAGCTGTAAAATTTGCCAAGGCTTTTGGGGCTAAGGTGAC AGTGATTAGTTCATCAGAAAGTAAAAAGGTTGAAGCCATTGAAAAATATG GTGCAGATTCCTTTTTGGTTAGCAGTGATCCAGGGCAGATGCTGGCAGCTG CCGGAACCTTGGATGGTGTCATTGATACCGTCCCAGCACCTCACTCTATTTT GCCATTCCTTGATTTACTCTTGCCTCGTGGAAAGCTAATTATATTAGGTGCA CCAATGGAGCCATTTGTACTGCCAATCTATCCCCTGCTTCAAGGTGGGAGA GTAGTTGCTGGGAGTGCCACTGGAGGATTGAAACAAATCCAAGAAATGCT TCATTTTGCAGCAGAGCACAACATAGTAGCAGATGGCGAGGTTATCCCAAT CGACGACATTAACACTGCGATAAAGCGCATTGAGAAAGGCGATGTCAAAT ATCGATTTGTGGTTGACATTGGCAATACCTTAAAATCTGCTTGA |
| *Cp*DCE (GenBank accession: MW456556.1) | ATGGAGAAACATTTTGTGCTAATCCATGGAGGTTGTTTCGGGGCATGGGCA TGGTACAAAGTGGTGACAATCTTGGAATCCAACGGCTACAAAGCCACTGC CCTTGATATGGCTTCTTCTGGGATCAATCCCAGACGTACAGACGAGGTGAA ATCCTACTCCGATTATTCTGAGCCGTTGATCAAGTTCATGGAGGATTTACC ATCAAATGAGAGAGTGGTTTTGGTTGGGCACAGTTTGGCTGGAGTTATTGT TTCTTTGGCTATGGAAAAATTCCCTCAGAAAATTGCTGCTGGCGTTTTCCTC ACTGCTGTCATGCCTGGTCCTGAGATCACTATGGCAACGCTTCATGAGGAG CACAAGGAACAAATTGATACTTTCATGGACTGCCAAATGATCCACGGCAAT GGCGATGATAATCCTCCGACTGCTTTCCTCTTTGGTCTCGAGTACTTAAAAT CCAAAGTGTTTCAACTCTGTCCTCCTGAGGACATGACACTCGCATCCCTTTT GGTAAGGCCAATATCTCTGGCAATCGAGCAAGGAGAACTAGACATCCCAC |

| | |
|---|---|
| | TCACCCAGATTAACTACGGTTCGGTTCCTCGTATTTACCTCATATCTGAAAA TGACAACGTGACAAAGGTCGAAGTTCAGAGATGGATGGTCGACAAAAATC CACCCGAAGAAGTCTTCGTGATCCCTAGTTGCGATCATATGGTCATGTTAT CCAATCCCAAAGATCTAAGCTCTCGTTTGCTGGAAATTGCCCAGAAATATG ACTAA |
| *Cr*SGD (GenBank accession: AF112888.1) | ATGGGATCTAAAGATGATCAGTCCCTTGTTGTTGCCATTTCTCCAGCTGCTG AACCAAATGGAAATCATTCTGTCCCCATCCCATTCGCCTACCCCAGTATCC CCATTCAACCTAGAAAGCACAACAAGCCCATCGTTCATCGTCGAGATTTCC CCTCAGATTTCATCTTGGGTGCCGGAGGATCTGCTTATCAGTGTGAGGGTG CATATAATGAAGGCAACCGCGGTCCCAGTATATGGGATACTTTCACAAACC GATATCCAGCCAAAATAGCTGATGGATCTAATGGCAATCAAGCCATCAATT CTTACAATTTGTACAAGGAAGATATCAAGATTATGAAGCAAACAGGCTTG GAATCATATAGGTTTTCAATTTCATGGTCAAGAGTATTGCCAGGTGGGAAT CTATCCGGTGGAGTGAATAAAGATGGTGTCAAGTTCTATCATGACTTTATA GATGAGCTTCTAGCCAATGGCATCAAACCCTTTGCAACTCTCTTCCACTGG GATCTTCCCCAAGCTCTTGAAGACGAGTATGGAGGCTTCTTGAGTGATCGA ATTGTGGAAGATTTTACGGAGTATGCAGAATTTTGCTTTTGGGAATTCGGT GACAAAGTAAAATTTTGGACGACTTTCAATGAACCACATACTTATGTTGCA AGTGGATATGCCACTGGTGAATTTGCACCAGGAAGAGGTGGTGCAGATGG CAAGGGGGAACCTGGCAAAGAACCCTATATAGCGACACATAATTTACTTCT TTCTCACAAAGCTGCTGTGGAAGTATATAGGAAAAATTTTCAGAAATGTCA AGGAGGTGAAATTGGAATTGTACTTAATTCAATGTGGATGGAGCCTCTCAA TGAAACCAAAGAAGATATTGATGCTCGGGAAAGGGGTCTTGATTTCATGCT CGGATGGTTCATAGAGCCATTAACAACGGGTGAATACCCAAAATCCATGA GAGCTCTTGTAGGAAGCCGTCTTCCAGAATTTTCAACAGAAGTTTCCGAAA AATTAACAGGATGCTATGATTTTATCGGAATGAATTATTATACAACTACTT ATGTTTCTAATGCAGACAAAATTCCCGATACTCCGGGTTACGAAACAGATG CTCGAATTAATAAGAATATTTTTGTCAAAAAAGTTGATGGGAAGGAAGTGC GCATTGGTGAACCGTGCTATGGGGGATGGCAGCATGTTGTTCCATCTGGAC TCTACAATCTCTTGGTTTACACTAAGGAGAAATACCATGTTCCAGTGATTT ATGTCTCAGAATGTGGTGTGGTTGAGGAAAATAGAACCAACATATTACTTA CAGAAGGTAAAACCAACATATTACTTACAGAAGCTCGTCACGATAAACTC AGGGTTGATTTTCTACAAAGTCATCTCGCTAGCGTGCGAGATGCTATTGAT GATGGTGTGAATGTAAAAGGATTCTTTGTTTGGTCATTCTTCGACAACTTCG AATGGAATTTGGGATATATATGCCGTTATGGAATTATCCATGTTGATTATA AAACTTTTCAAAGATATCCAAAGGATTCTGCCATATGGTACAAGAATTTCA TTAGTGAAGGATTTGTTACGAATACAGCTAAAAAGAGATTCCGAGAAGAA GATAAACTAGTTGAGTTAGTCAAGAAGCAAAAATACTAA |

**Table S2.** Primers used for genes cloning in this study.

| Gene | Plasmid | Primer direction | Sequence |
|---|---|---|---|
| *Cp*T5H1 | 3Ω1 | Forward | TTTATGAATTTTGCAGCTCGATGGAGAATTTTACAGTTGAAATC |
| | | Reverse | GACAACCACAACAAGCACCGCTAATTCTTGATCATTGTAGGA |
| *Cp*T5H2 | 3Ω1 | Forward | TTTATGAATTTTGCAGCTCGATGGAGAATTTTACAGTTGAAGTC |
| | | Reverse | GACAACCACAACAAGCACCGCTAATTCTTGATCATTGTAGGA |
| *Cp*OMT1 | 3Ω1 | Forward | TTTATGAATTTTGCAGCTCGATGGCAACATCTGAGAATTCTGCTGAGC |
| | | Reverse | GACAACCACAACAAGCACCGCTAAGGATAAACCTCAATGATACT |
| *Cp*STR | 3Ω1 | Forward | TTTATGAATTTTGCAGCTCGATGCACATTTCTGAAAATATG |
| | | Reverse | GACAACCACAACAAGCACCGCTAGAAAGAAAACGACTCTCCTTTC |
| *Cp*STTr | 3Ω1 | Forward | TTTATGAATTTTGCAGCTCGATGGAAGCTGCAGTGATGTCT |
| | | Reverse | GACAACCACAACAAGCACCGCTAGTAGTACAACGTTGCATCC |
| *Cp*THAS1 | 3Ω1 | Forward | TTTATGAATTTTGCAGCTCGATGGCTGAAAAATCAGCAATAG |
| | | Reverse | GACAACCACAACAAGCACCGCTAATCAGATTTGAGCGTTTTTC |
| *Cp*THAS2 | 3Ω1 | Forward | TTTATGAATTTTGCAGCTCGATGGCTGAAAAACCAGCATTC |
| | | Reverse | GACAACCACAACAAGCACCGCTAATCACATTTAAGCGTGTTTC |
| *Cp*DCS | 3Ω1 | Forward | TTTATGAATTTTGCAGCTCGATGATGGCCGGAAAATCTCAAG |
| | | Reverse | GACAACCACAACAAGCACCGCTAGCCTAGGTCTCTCG |
| *Cp*DCE | 3Ω1 | Forward | TTTATGAATTTTGCAGCTCGATGGAGAAACATTTTGTGCTAATC |
| | | Reverse | GACAACCACAACAAGCACCGCTAGTCATATTTCTGGGCAATTTC |
| *Cp*TDC | 3Ω1 | Forward | TTTATGAATTTTGCAGCTCGATGGGCAGCATTGATGCTAATA |
| | | Reverse | GACAACCACAACAAGCACCGCTAATGATTCTCCTTAAGCAAGG |
| *Cp*OMT1 | pOPINF | Forward | AAGTTCTGTTTCAGGGCCCGGCAACATCTGAGAATTCTGCTGAGC |
| | | Reverse | ATGGTCTAGAAAGCTTTAAGGATAAACCTCAATGATACT |
| Primers for sequencing | | | |
| 3Ω1 | | Forward | GATGAAAAAGCCCTAAAATTGGAG |
| | | Reverse | ATTATTCACAAATGAGAAACAGAATGG |
| pOPINF | | Forward | CGAAATTAATACGACTCACTATAGG |
| | | Reverse | TAGCCAGAAGTCAGATGCT |

**Table S3.** Names, plant species and accession numbers for the enzymes used in the phylogenetic tree of the *Cp*OMT1 (Figure S13).

| | | | |
|---|---|---|---|
| Cr16OMT | 16-hydroxytabersonine O-methyltransferase | *Catharanthus roseus* | EF444544 |
| Vm16OMT | 16-hydroxyvincadifformine 16-O-methyltransferase | *Vinca minor* | MH010798 |
| CrFlav4OMT | flavonoid 4 -O-methyltransferase | *Catharanthus roseus* | Q6VCW3 |
| CrOMT2 | Myricetin O-methyltransferase | *Catharanthus roseus* | Q8GSN1 |
| TiN10OMT | noribogaine 10-O-methyltransferase | *Tabernanthe iboga* | MH454075 |
| Ci67OMT | 7 -O-demethylcephaeline O-methyltransferase | *Carapichea ipecacuanha* | BAJ72588 |
| IpeOMT1 | O-methyltransferase | *Psychotria ipecacuanha* | AB527082 |
| GsRH11OMT | rankinidine/humantenine-11-O-methyltransferase | *Gelsemium sempervirens* | MF401947 |
| SnvOMT | strychnine O-methyltransferase | *Strychnos nux-vomica* | OM304297 |
| Ca10OMT | 10-hydroxycamptothecin O-methyltransferase | *Camptotheca acuminata* | MG996006 |
| CaResOMT | resveratrol O-methyltransferase | *Camptotheca acuminata* | MG996007 |
| EjOMT4 | O-methyltransferase | *Eriobotrya japonica* | LC127204 |
| RhcOrcOMT1 | orcinol O-methyltransferase | *Rosa hybrid cultivar* | AF502433 |
| MtI4OMT | Isoflavone 4 -O-methyltransferase | *Medicago truncatula* | Q29U70 |
| CjCo4OMT | 3 -hydroxy-N-methyl-(S)-coclaurine 4 -O-methyltransferase | *Coptis japonica* | Q9LEL5 |
| Ps4OMT2 | 3 -hydroxy-N-methyl-(S)-coclaurine 4 -O-methyltransferase 2 | *Papaver somniferum* | Q7XB10 |
| TfNc6OMT | (S)-norcoclaurine 6-O-methyltransferase | *Thalictrum flavum* | AAU20765 |
| PsN6OMT | (R S)-norcoclaurine 6-O-methyltransferase | *Papaver somniferum* | AAQ01669.1 |
| PsN7OMT | norreticuline-7-O-methyltransferase | *Papaver somniferum* | ACN88562.1 |
| CjCoOMT | Tetrahydrocolumbamine 2-O-methyltransferase | *Coptis japonica* | Q8H9A8 |
| HvFlavOMT | flavonoid 7-O-methyltransferase | *Hordeum vulgare* | CAA54616 |
| OsASMT1 | Acetylserotonin O-methyltransferase | *Oryza sativa* | Q6EPG8J |
| PsR7OMT | (R S)-reticuline 7-O-methyltransferase | *Papaver somniferu* | AAQ01668 |
| AtASMT | Acetylserotonin O-methyltransferase | *Arabidopsis thaliana* | Q9T003 |
| OsOMT1 | Flavone 3 -O-methyltransferase | *Oryza sativa* | Q6ZD89 |
| AtOMT1 | Flavone 3 -O-methyltransferase | *Arabidopsis thaliana* | Q9FK25 |
| LwOMT10 | O-methyltransferase | *Lophophora williamsii* | OQ831042 |
| Hp9OMT | corynanthe 9-O-methyltransferase | *Hamelia patens* | WKU83439 |
| LwOMT2 | O-methyltransferase | *Lophophora williamsii* | OQ831037 |
| PsCoNMT | coclaurine N-methyltransferase | *Papaver somniferum* | AAP45316 |
| TfCoNMT | coclaurine N-methyltransferase | *Thalictrum flavum* | AAU20766.1 |
| PsTNMT | tetrahydroprotoberberine-cis-N-methyltransferase | *Papaver somniferum* | AAY79177 |
| CrNMT | hydroxytabersonine n-methyltransferase | *Catharanthus roseus* | AHH33092 |
| ASMT | Acetylserotonin O-methyltransferase | *Homo sapiens* | P46597 |
| VpOMT4 | O-methyltransferase | *Vanilla planifolia* | EF444544 |
| MsCaffCoA3OMT | trans-caffeoyl-CoA 3-O-methyltransferase | *Medicago sativa* | Q40313 |
| NpN4OMT | norbelladine 4 -O-methyltransferase | *Narcissus sp.* | AIL54541 |
| OsMTS1 | O-methyltransferase | *Oryza sativa* | Q6YSY5 |
| CarMXNMT | 7-methylxanthine N-methyltransferase | *Coffea arabica* | BAB39216 |
| CrLAMT | loganic acid methyltransferase | *Catharanthus roseus* | ABW38009 |
| MsEnOMT | corynanthe C17 O-methyltransferase | *Mitragyna speciosa* | WKU61913 |

[10] M. Wang, J. J. Carver, V. V. Phelan, L. M. Sanchez, N. Garg, Y. Peng, D. D. Nguyen, J. Watrous, C. A. Kapono, T. Luzzatto-Knaan, C. Porto, A. Bouslimani, A. V. Melnik, M. J. Meehan, W.-T. Liu, M. Crüsemann, P. D. Boudreau, E. Esquenazi, M. Sandoval-Calderón, R. D. Kersten, L. A. Pace, R. A. Quinn, K. R. Duncan, C.-C. Hsu, D. J. Floros, R. G. Gavilan, K. Kleigrewe, T. Northen, R. J. Dutton, D. Parrot, E. E. Carlson, B. Aigle, C. F. Michelsen, L. Jelsbak, C. Sohlenkamp, P. Pevzner, A. Edlund, J. McLean, J. Piel, B. T. Murphy, L. Gerwick, C.-C. Liaw, Y.-L. Yang, H.-U. Humpf, M. Maansson, R. A. Keyzers, A. C. Sims, A. R. Johnson, A. M.

Sidebottom, B. E. Sedio, A. Klitgaard, C. B. Larson, C. A. Boya P, D. Torres-Mendoza, D. J. Gonzalez, D. B. Silva, L. M. Marques, D. P. Demarque, E. Pociute, E. C. O'Neill, E. Briand, E. J. N. Helfrich, E. A. Granatosky, E. Glukhov, F. Ryffel, H. Houson, H. Mohimani, J. J. Kharbush, Y. Zeng, J. A. Vorholt, K. L. Kurita, P. Charusanti, K. L. McPhail, K. F. Nielsen, L. Vuong, M. Elfeki, M. F. Traxler, N. Engene, N. Koyama, O. B. Vining, R. Baric, R. R. Silva, S. J. Mascuch, S. Tomasi, S. Jenkins, V. Macherla, T. Hoffman, V. Agarwal, P. G. Williams, J. Dai, R. Neupane, J. Gurr, A. M. C. Rodríguez, A. Lamsa, C. Zhang, K. Dorrestein, B. M. Duggan, J. Almaliti, P.-M. Allard, P. Phapale, et al., *Nat. Biotechnol.* **2016**, *34*, 828-837.

[11] http://gnps.ucsd.edu

[12] L.-F. Nothias, D. Petras, R. Schmid, K. Dührkop, J. Rainer, A. Sarvepalli, I. Protsyuk, M. Ernst, H. Tsugawa, M. Fleischauer, F. Aicheler, A. A. Aksenov, O. Alka, P.-M. Allard, A. Barsch, X. Cachet, A. M. Caraballo-Rodriguez, R. R. Da Silva, T. Dang, N. Garg, J. M. Gauglitz, A. Gurevich, G. Isaac, A. K. Jarmusch, Z. Kameník, K. B. Kang, N. Kessler, I. Koester, A. Korf, A. Le Gouellec, M. Ludwig, C. Martin H, L.-I. McCall, J. McSayles, S. W. Meyer, H. Mohimani, M. Morsy, O. Moyne, S. Neumann, H. Neuweger, N. H. Nguyen, M. Nothias-Esposito, J. Paolini, V. V. Phelan, T. Pluskal, R. A. Quinn, S. Rogers, B. Shrestha, A. Tripathi, J. J. J. van der Hooft, F. Vargas, K. C. Weldon, M. Witting, H. Yang, Z. Zhang, F. Zubeil, O. Kohlbacher, S. Böcker, T. Alexandrov, N. Bandeira, M. Wang, P. C. Dorrestein, *Nat. Methods* **2020**, *17*, 905-908.
